# Multiple horizontal mini-chromosome transfers drive genome evolution of clonal blast fungus lineages

**DOI:** 10.1101/2024.02.13.580079

**Authors:** A. Cristina Barragan, Sergio M. Latorre, Angus Malmgren, Adeline Harant, Joe Win, Yu Sugihara, Hernán A. Burbano, Sophien Kamoun, Thorsten Langner

## Abstract

Crop disease pandemics are often driven by clonal lineages of plant pathogens that reproduce asexually. How these clonal pathogens continuously adapt to their hosts despite harboring limited genetic variation, and in absence of sexual recombination remains elusive. Here, we reveal multiple instances of horizontal chromosome transfer within pandemic clonal lineages of the blast fungus *Magnaporthe* (Syn. *Pyricularia) oryzae*. We identified a horizontally transferred 1.2Mb supernumerary mini-chromosome which is remarkably conserved between *M. oryzae* isolates from both the rice blast fungus lineage and the lineage infecting Indian goosegrass (*Eleusine indica*), a wild grass that often grows in the proximity of cultivated cereal crops. Furthermore, we show that this mini-chromosome was horizontally acquired by clonal rice blast isolates through at least nine distinct transfer events over the past three centuries. These findings establish horizontal mini-chromosome transfer as a mechanism facilitating genetic exchange among different host-associated blast fungus lineages. We propose that blast fungus populations infecting wild grasses act as genetic reservoirs that drive genome evolution of pandemic clonal lineages that afflict cereal crops.

## Introduction

Coevolutionary dynamics between plants and their pathogens date back millions of years and are a central force shaping both sets of genomes (Barragan and Weigel 2021). In such antagonistically interacting organisms, a cycle of adaptation and counter-adaptation must occur to avoid extinction (Van Valen 1973). This evolution relies not only on the acquisition of novel mutations but also on the preservation of long-standing genetic variation; together, these components provide the genetic foundations upon which selective pressures act (Nei 2007; Barrett and Schluter 2008). In eukaryotes, one of the major sources of genetic variation is recombination through sexual mating, yet many organisms, including fungal plant pathogens, preferentially reproduce asexually (Barrett 2010; Möller and Stukenbrock 2017). The absence of sexual recombination necessitates alternative mechanisms for generating genetic variability, including mutations, genomic rearrangements, transposon insertion, and gene duplication or loss (Seidl and Thomma 2014; Oggenfuss et al. 2023). However, these processes rely primarily on pre-existing genetic variation, and without the introduction of new genetic material, the adaptive potential of an asexual population is constrained. How clonal plant pathogens adapt to their hosts and avoid extinction despite harboring limited genetic variation is an important research question with practical implications, as clonal lineages of plant pathogens often drive disease pandemics in crops (Drenth et al. 2019).

One mechanism for acquiring genetic variation which does not require sexual mating, is horizontal gene transfer (HGT). This process, consisting of the transmission of genetic material from a donor to a recipient organism within the same generation, is considered a major force in preventing extinction in asexual organisms (Takeuchi et al. 2014). In prokaryotes, HGT is well-established as a source of genetic diversity, occurring through known mechanisms such as conjugation, transformation, or transduction (Sun 2018). The prevalence of HGT in eukaryotes has also become more apparent in recent years (Gabaldón 2020), particularly within the fungal kingdom – one of the most extensively studied eukaryotic lineage (Fitzpatrick 2012; Mohanta and Bae 2015; Sahu et al. 2023). In fungi, parasexuality, a mechanism enabling chromosome reassortment independent of sexual reproduction (Nieuwenhuis and James 2016), is a plausible avenue for HGT.

Fungal genes acquired by HGT are often part of the non-essential accessory genome which is variable between individuals of the same species and contrasts to the core genome, which contains genes essential to housekeeping functions (McCarthy and Fitzpatrick 2019). This is in line with the “two-speed” genome model observed in some filamentous plant pathogens (fungi and oomycetes), where indispensable genomic regions are under higher evolutionary constraints and may appear as slow-evolving, while variable genomic regions are under more relaxed constraints or positive selection, and can appear as rapidly-evolving (Dong et al. 2015). Rapidly-evolving or dynamic genome compartments are characterized by the presence of virulence genes, high sequence diversification, presence/absence variation, structural changes, and segmental duplications (Torres et al. 2020; Huang et al. 2023). An extreme form of structural variation are mini-chromosomes (mChr), also referred to as supernumerary, accessory, or B chromosomes, which exist in addition to core chromosomes and have been found in 15% of eukaryotic species (Covert 1998). While mChr emergence has been associated with genomic rearrangements at repeat- and effector-rich subtelomeric ends of core chromosomes (Bertazzoni et al. 2018; Peng et al. 2019; Langner et al. 2021; van Westerhoven et al. 2023), the exact molecular mechanism remain an area of ongoing investigation. By being physically unlinked from core chromosomes, mChr can diversify rapidly and could serve as a cradle for adaptive evolution without compromising genomic integrity (Croll and McDonald 2012).

The adaptive role of mChr in plant pathogenic fungi is underpinned by their correlation to virulence in various pathogen-host systems (Miao et al. 1991; Kistler 1996; Han et al. 2001; Akagi et al. 2009; Ma et al. 2010; Chuma et al. 2011; Balesdent et al. 2013; van Dam et al. 2017; Habig et al. 2017; Bhadauria et al. 2019; Henry et al. 2021; Asuke et al. 2023). In addition, variation in virulence has been partly attributed to the horizontal transfer of mChr (Mehrabi et al. 2011). This is exemplified in the case of *Fusarium oxysporum*, where the horizontal acquisition of a mChr in laboratory settings transformed a non-pathogenic strain into a virulent pathogen (Ma et al. 2010). Similarly, in the insect pathogen *Metarhizium robertsii,* strains with a horizontally acquired mChr were more virulent compared to those without this mChr (Habig et al. 2023).

A notorious plant pathogenic fungus where asexually reproducing clonal lineages underlie crop pandemics, is the blast fungus *Magnaporthe oryzae* (Syn. *Pyricularia oryzae*) (Latorre et al. 2020; Latorre et al. 2023). The blast fungus is one of the most devastating plant pathogens worldwide and is the causal agent of blast disease in dozens of wild and cultivated grasses (Islam et al. 2023). As a species, *M. oryzae* is differentiated into genetic lineages that tend to be host-associated, with occasional gene flow observed between certain lineages (Couch et al. 2005; Gladieux, Condon, et al. 2018). The highly destructive rice blast fungus lineage reproduces mostly asexually in nature, with limited traces of sexual reproduction having been found (Saleh et al. 2012; Thierry et al. 2022). Nevertheless, the rice blast fungus lineage remains genetically isolated, with no gene flow detected from other *M. oryzae* lineages so far (Gladieux, Condon, et al. 2018). To date, three globally prevalent clonal lineages affecting rice and one affecting wheat have been identified as the underlying cause of persistent blast pandemics (Latorre et al. 2020; Latorre et al. 2023). Despite their restricted genetic diversity, clonal *M. oryzae* lineages readily evolve to counteract host defenses, posing a challenge to the development of durable blast-resistant crop varieties (Younas et al. 2023). The mechanism that allows clonal blast fungus populations to adapt to new host germplasm, despite an apparent lack of avenues for genetic innovation, remains elusive.

Structural variation in both mChr and core chromosomes contribute to genomic diversity in the blast fungus (Talbot et al. 1993; Orbach et al. 1996). Recent genomic analysis of a wheat blast fungus isolate revealed multi-megabase insertions from a related species, suggesting HGT (Kobayashi et al. 2023). In addition, the postulated horizontal transfer of the avirulence gene AVR-Pita2 among related species to the blast fungus substantiates the hypothesis that HGT is occurring (Chuma et al. 2011). While these instances highlight HGT as a possible driver of genetic variation in the blast fungus, the exact mechanisms facilitating HGT remain unclear. In addition to gene transfer, mChr have been associated with virulence gene reshuffling and recombination with core chromosomes (Kusaba et al. 2014; Peng et al. 2019; Langner et al. 2021; Asuke et al. 2023; Gyawali et al. 2023), indicating that horizontal mChr transfer could be instrumental in driving genomic innovation.

In this study, we provide evidence for multiple horizontal mini-chromosome transfer events involving clonal lineages of the rice blast fungus *M. oryzae* that occurred under field conditions. We identified a 1.2Mb supernumerary mini-chromosome, mChrA, which is remarkably conserved across *M. oryzae* isolates from lineages infecting the wild host species, Indian goosegrass (*Eleusine indica*), and rice. We show that mChrA was acquired by clonal rice blast fungus lineages through at least nine independent horizontal transfer events over the past three centuries. This establishes horizontal mChr transfer as a naturally occurring genetic exchange mechanism among different host-associated blast fungus lineages. Our findings lead us to propose that blast fungus lineages infecting wild grasses serve as genetic reservoirs, driving genome evolution of pandemic asexual clonal lineages that afflict crops.

## Results

### Clonal rice blast fungus isolates display variable mChr content

We have previously shown that genetically diverse *M. oryzae* isolates exhibit variable mChr content (Langner et al., 2021). Here, we set out to analyze the extent to which mChr variation contributes to genomic diversity in a set of genetically related isolates belonging to a single clonal lineage. To this end, we selected nine rice blast fungus isolates collected from Italy (Win et al. 2020) (**Fig S1A** and **Table S1**). Using genome-wide single-nucleotide polymorphism (SNP) data we confirmed that the nine isolates belong to a single clonal lineage (clonal lineage II), which is predominant in Europe (Latorre et al. 2020; Thierry et al. 2022) (**Fig 1A**, **S1B** and **Table S2**).

**Fig 1.**
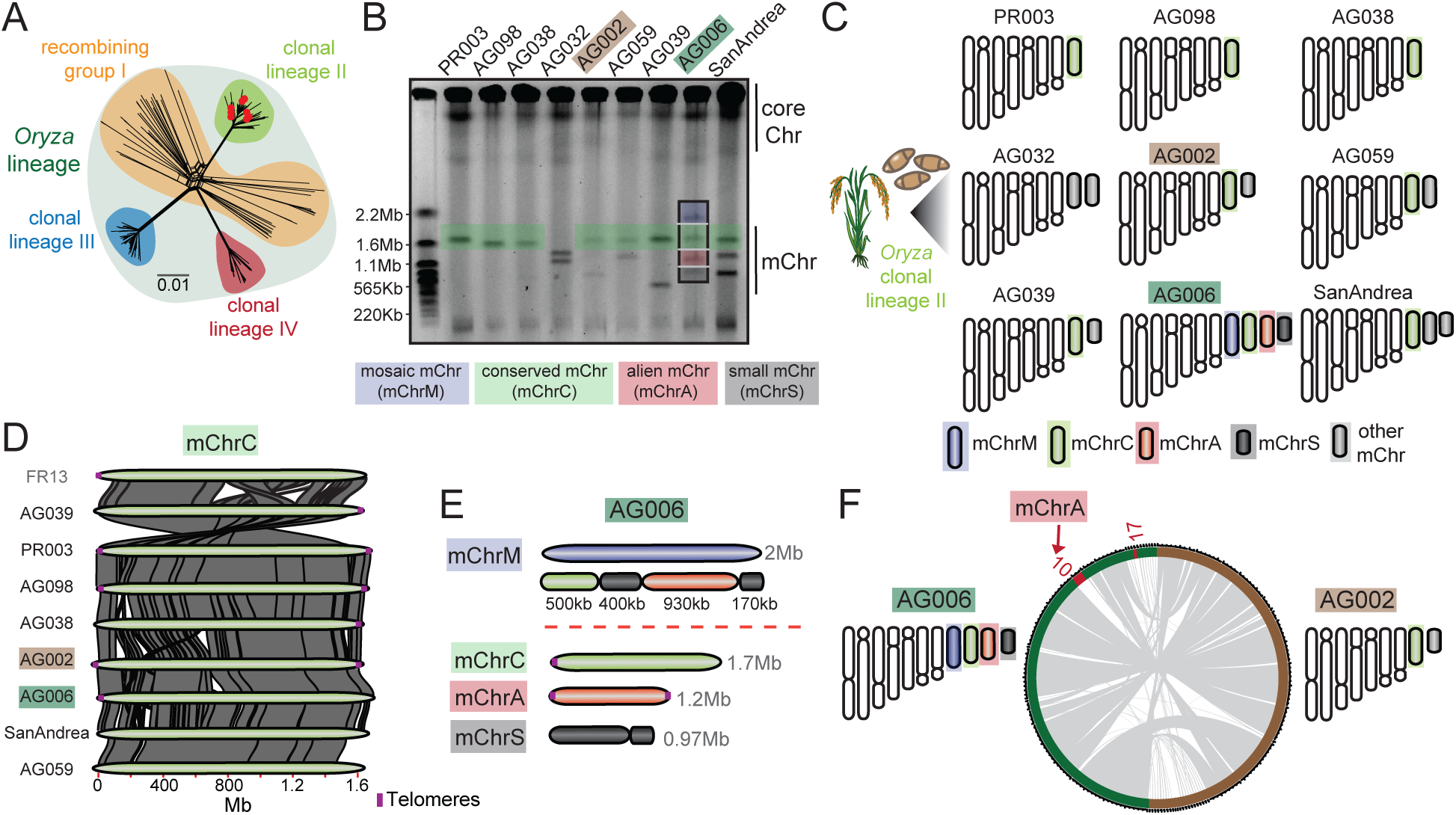
Clonal rice blast fungus isolates display variable mChr content. **A.** Genome-wide SNP-based NeighborNet analysis confirms the nine rice blast fungus isolates (red dots) belong to clonal lineage II (green) (Latorre et al. 2020). **B.** CHEF gel karyotyping reveals variable mChr content. A conserved 1.7Mb mChr (mChrC, green) is found in eight of nine isolates. A 2Mb mChr (mChrM, blue), present in isolate AG006, is a mosaic composed of fragments from three other mChr (mChrC, mChrA, and mChrS; see panel E) from the same isolate. A third 1.2Mb mChr (mChrA, red) found in AG006, is absent from the genomes of the other isolates (see panels E and F). **C**. Schematic karyotype of Italian isolates. Core chromosomes are shown in white. mChr studied in detail are highlighted in colors, while the rest are in gray. **D.** mChrC exhibits high synteny across isolates and is also found in isolate FR13 (Langner et al. 2021). Telomeric sequences are indicated by a vertical line (purple). **E.** Inferred mChrM sequence composition. **F.** Whole-genome alignment between AG006 (green) and AG002 (brown). mChrA (AG006_Contig10) and AG006_Contig17 (in red) are absent from AG002.

To determine the karyotype of the selected isolates, we performed contour-clamped homogeneous electric field based (CHEF) electrophoresis. This revealed variable numbers and sizes of mChr, with each isolate exhibiting one to four mChr, each varying from 0.5 to 2Mb in size (**Fig 1B** and **1C**). To genetically characterize individual mChr, we performed mini-chromosome isolation sequencing (MCIS) on all eighteen mChr found across the nine isolates (Langner et al. 2019; Langner et al. 2021). Reads obtained from each individual mChr were mapped to the reference assembly corresponding to its originating isolate. Contigs exhibiting high MCIS coverage and high repeat density, a characteristic trait of mChr, and were <2Mb in size, were identified as mChr contigs (**Fig S2**, **S3**, **Table S3** and **S4**).

Next, we compared mChr contigs across the studied clonal isolates. Reciprocal sequence homology searches revealed the presence of a conserved 1.7Mb mChr (mChrC) in eight out of the nine isolates. mChrC corresponds to a previously identified mChr found in the rice blast fungus isolate FR13, which also belongs to clonal lineage II (Langner et al. 2021). We aligned mChrC contigs and confirmed high synteny across isolates (**Fig 1D**). To obtain an overview of how common mChrC is in the global *M. oryzae* population, we examined the presence of this sequence across 413 *M. oryzae* and *Magnaporthe grisea* isolates (**Table S5**). We performed short-read mapping to the AG006 genome, known from karyotyping to possess the highest mChr diversity. Subsequent breadth of coverage calculations (see Methods) revealed that mChrC is particularly conserved among rice blast fungus isolates, especially those belonging to clonal lineage II (**Fig S4A**-**C** and **Table S6**).

*M. oryzae* isolate AG006 stood out within the examined set of isolates as it contained three additional mChr named mChrS, mChrA, and mChrM, in addition to mChrC. These mChr exhibited sizes ranging from 0.97 to 2Mb. Notably, the largest of these mChr, which we termed the mosaic mini-chromosome (mChrM), was composed of segments derived from the three smaller mChr, namely, mChrC, mChrA, and mChrS (**Fig 1B, 1C, 1E** and **S2**). The presence of this mosaic mChr reveals that recombination among mChr occurs, and plays a role in generating novel genetic combinations.

Upon closer examination of mChrA, we found it exhibited low sequence similarity to the genomes of the other Italian isolates, as evidenced by reciprocal sequence homology searches (**Fig 1B, 1C, 1E** and **Table S7** and **S8**). There were two exceptions to this, a duplicated fragment within mChrM (**Fig 1E**), and a small 0.1Mb contig (AG006_Contig17) which aligned to a specific region of mChrA (AG006_Contig10) (**Fig S3A**). The latter may have originated from a sequence duplication event or be an assembly artifact. mChrA displayed high MCIS coverage and canonical telomeric repeats at both ends, indicating it is linear and largely assembled into a single contig (**Fig S5**). Lastly, to reinforce these findings, we conducted a whole-genome alignment between AG006 and AG002, an isolate genetically highly similar to AG006 (**Fig S1B**), confirming the absence of mChrA in AG002 (**Fig 2F**).

**Fig 2.**
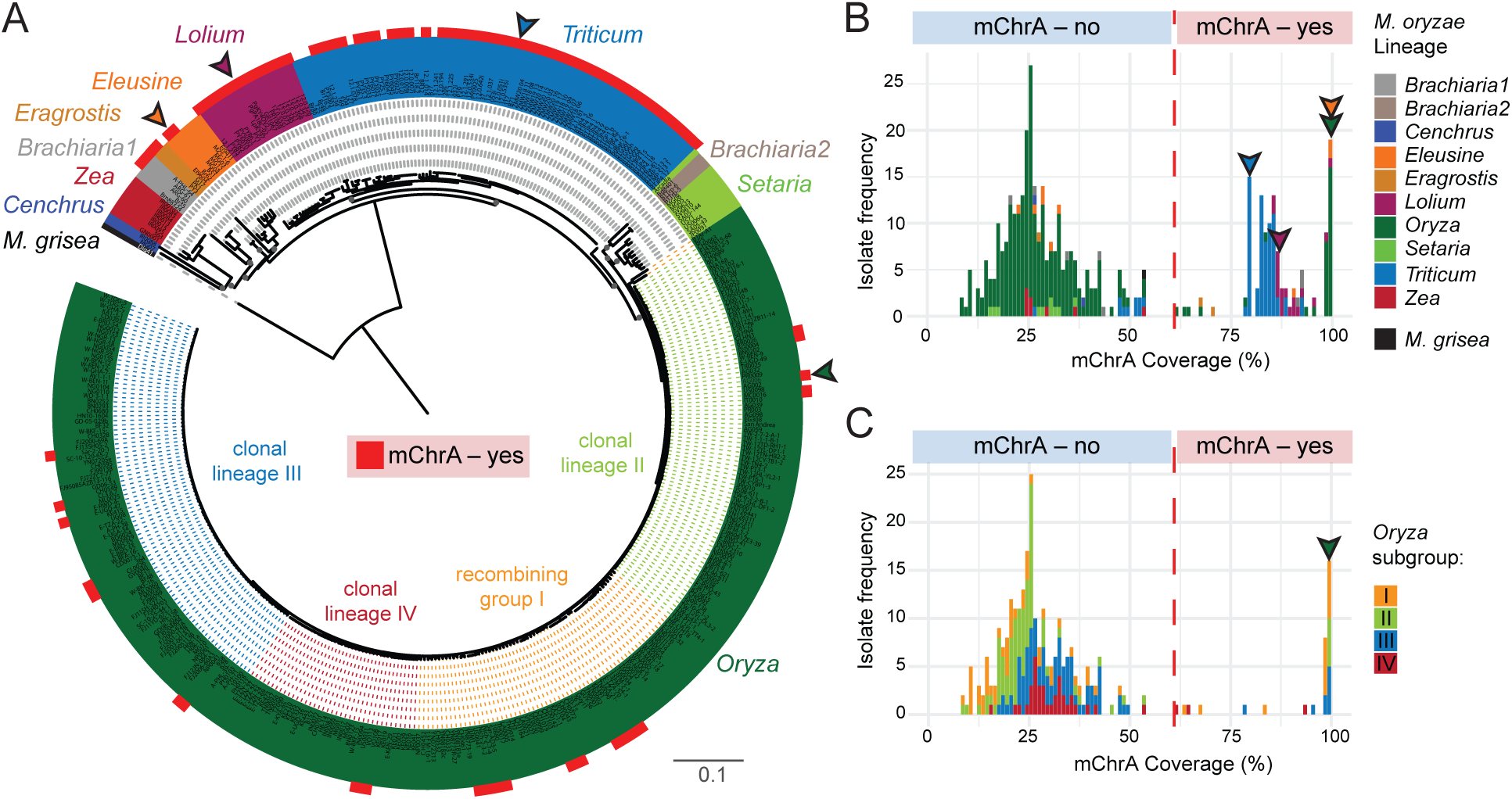
mChrA sequences are present across multiple host-associated blast fungus lineages. **A.** Genome-wide SNP-based NJ tree of 413 *M. oryzae* and *M. grisea* isolates. *M. oryzae* isolates are color-coded by lineage and *M. grisea* is in black. The 126 isolates defined as mChrA carriers (Table S9) are highlighted by a red square and belong to six different *M. oryzae* lineages. Arrows indicate isolates with mChrA-related karyotyping information (Peng et al. 2019; Rahnama et al. 2020), see Fig 4). Colors of dotted lines across the rice blast lineage represent different genetic subgroups (three clonal lineages and a recombining group) (Latorre et al. 2020). Scale bar represents nucleotide substitutions per position. **B.** Bimodal distribution of mChrA breadth of coverage across 413 *M.oryzae* and *M. grisea* isolates. The coverage cutoff (61%) for mChrA presence or absence is indicated by the dotted red line. Arrows as in A. **C.** mChrA breath of coverage across 276 rice blast fungus isolates. Colors represent different genetic subgroups in the *Oryza* lineage. Arrows and coverage cutoff as in B.

Taken together, we found high mChr diversity in a collection of nine clonal rice blast fungus isolates. Remarkably, we identified a unique mChr, mChrA, which does not display sequence similarity to the other nine rice blast fungus isolates. This finding underscores the unique genetic variation present even among closely related blast fungus isolates.

### mChrA sequences are present across multiple host-associated blast fungus lineages

To determine the origin of mChrA, we assessed the presence of the mChrA sequence (AG006_Contig10) across a set of 413 *M. oryzae* isolates belonging to ten different host-associated lineages and to *M. grisea* (**Fig 2A** and **Table S5**). For this purpose, we calculated the breadth of coverage for mChrA (AG006_Contig10) in each isolate, and observed it followed a bimodal distribution (**Fig 2B**). Model-based clustering established 126 isolates as mChrA carriers (**Fig S6A**-**B** and **Table S9**, see Methods). These isolates belonged to six different host-associated *M. oryzae* lineages (**Fig 2A, B** and **S6D**). Notably, only 12% of rice blast fungus isolates (32 of 276) were identified as mChrA carriers (**Fig 2A, B** and **Table S9**). These rice blast fungus isolates were genetically diverse, belonging to all three clonal lineages and the recombining group (defined here as *Oryza* subgroups) (**Fig 2C** and **S6E**).

In contrast to rice blast fungus isolates, the mChrA sequence was common across isolates belonging to the *Lolium* and *Triticum* lineages, with most isolates carrying 80-90% of the mChrA sequence (**Fig 2B** and **S6D**). Previous karyotyping and sequencing efforts identified isolate B71 from the *Triticum* lineage and isolate LpKY97 from the *Lolium* lineage to carry mChr (Peng et al. 2019; Rahnama et al. 2020). Given that these isolates also carry a substantial portion of the mChrA sequence (79% and 87% mapping to mChrA in AG006, respectively), it suggests that the mapped mChrA sequences may share a common ancestry with the mChr in these two isolates. However, the most striking sequence identity was observed in two isolates from the *Eleusine* blast fungus lineage, Br62 and B51. These exhibited mChrA coverage comparable to the rice blast fungus isolate AG006, suggesting high similarity in the mChrA sequences between these isolates (**Fig 2B** and **Table S6**). Taken together, mChrA sequences were found in blast fungus isolates belonging to six different host-associated lineages, with members of the *Eleusine* and *Oryza* lineages carrying nearly identical mChrA sequences.

### Discordant genetic clustering between the core genome and mChrA

Given the patchy distribution of mChrA sequences across isolates from different host-associated blast fungus lineages, we investigated the evolutionary relationships of their core genome and mChrA sequences. Using SNP-based phylogenies and principal component analyses (PCA), we found a clear discordance between the core genome and mChrA (**Fig 3A-D**). For the core genome, isolates are clustered by lineage, whereas for mChrA, lineage-dependent clustering becomes less evident. Most strikingly, whereas the core genomes of isolates from the *Oryza* and *Eleusine* lineages form two distinct groups (**Fig 3A, B** and **S7A**), these two fall within a single group for mChrA (**Fig 3C, D** and **S7B**). This shows that the mChrA sequence in isolates from these two lineages is highly similar, but their core genome is divergent. We note that for the mChrA clustering, three clonal rice blast fungus isolates did not group with other isolates from this lineage, possibly reflecting mChrA sequence dissimilarity.

**Fig 3.**
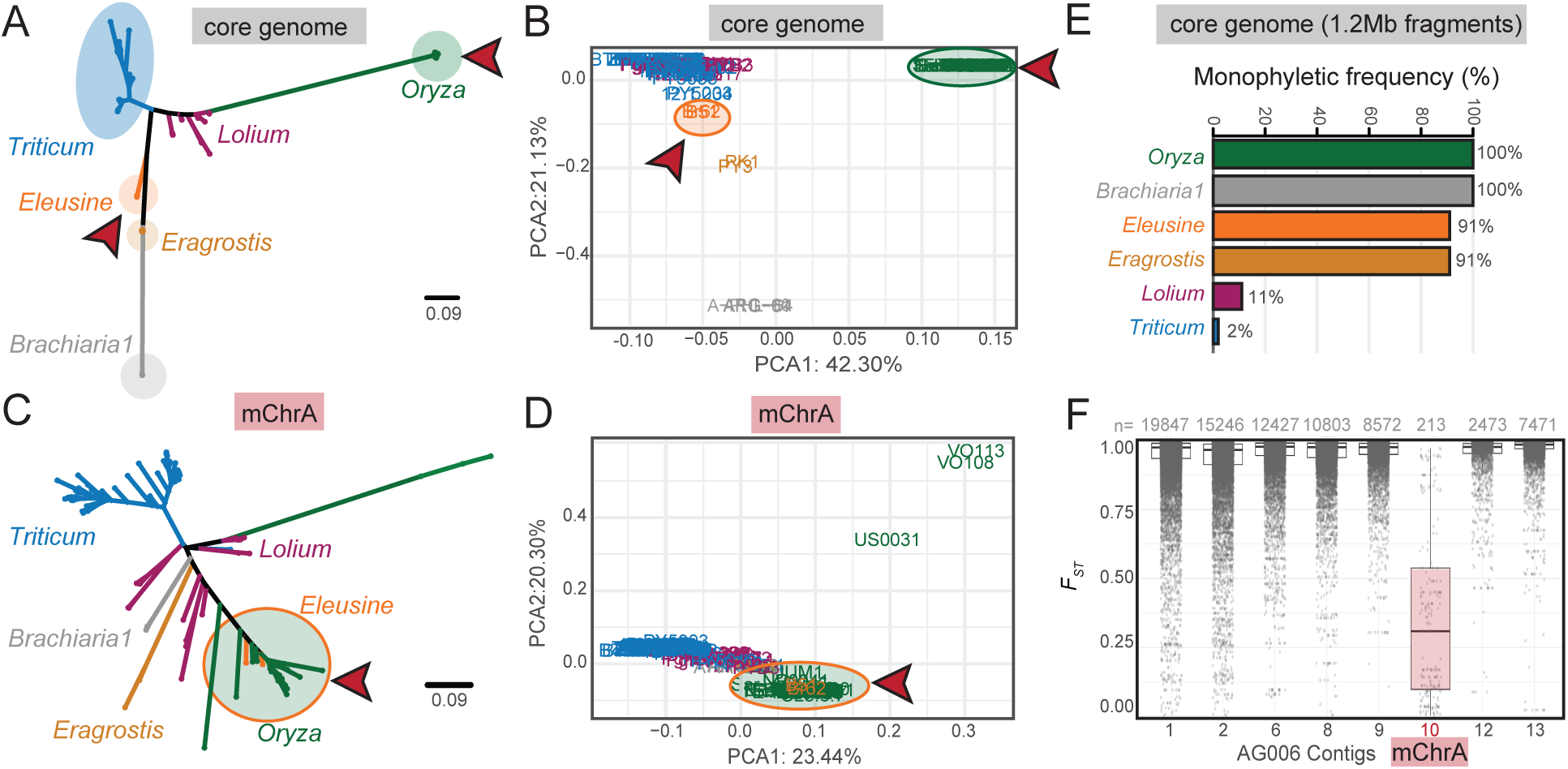
Discordant genetic clustering between the core genome and mChrA. **A-D.** SNP-based NJ trees (A and C) and Principal Component Analyses (PCA, B and D) of 126 *M. oryzae* isolates carrying the mChrA sequence. Discordance between core genome (A and B) and mChrA (C and D) genetic clustering is observed (red arrows). Scale bar represents nucleotide substitutions per position. **E.** Percentage of tree topologies where a monophyletic relationship was observed for 100 randomly selected 1.2Mb core-chromosomal regions. **F.** *F_ST_* between rice blast fungus isolates (n=32) and isolates from the *Eleusine* lineage (Br62 and B51) both carrying mChrA. Each dot (gray) indicates the weighted *F_ST_* per 5kb window using a step size of 500bp. The number of windows per contig are at the top of each box. Core chromosome contigs >2Mb and mChrA are shown.

To ascertain the robustness of the observed genetic clustering of mChrA between members of the *Oryza* and *Eleusine M. oryzae* lineages, we generated phylogenies using 100 randomly selected genomic regions of the same size as mChrA (1.2Mb) across the core genome of all 126 isolates. In all instances, the *Oryza* lineage was monophyletic, and in zero instances did the *Oryza* and *Eleusine* lineages cluster together (**Fig 3E**). This demonstrates that the clustering of members of the *Eleusine* and rice blast fungus lineages is highly unusual and limited to the mChrA sequence. To complement this analysis, we evaluated genetic differentiation between isolates belonging to the rice and *Eleusine* blast fungus lineages carrying the mChrA sequence by calculating the fixation index (F*_ST_*) from genome-wide SNP data (Wright 1951). This analysis confirmed high levels of inter-lineage genetic differentiation in the core genome, but low differentiation levels for mChrA (**Fig 3E** and **S8**). We conclude that mChrA shows discordant genetic clustering when compared to the core genome, which indicates contrasting evolutionary trajectories.

### *Eleusine* isolate Br62 and *Oryza* isolate AG006 carry an intact and highly syntenic mChrA

Following the identification of highly similar mChrA sequences in two isolates from the *Eleusine* blast fungus lineage, we aimed to determine whether these sequences originate from an intact mChr, or whether they are embedded within the core genome, as observed for mChrC segments in *M. oryzae* isolate 70-15 (Langner et al. 2021). To test this, we performed CHEF-gel based karyotyping, and found that Br62 possesses a single mChr of the same size (1.2Mb) as mChrA in AG006 (**Fig 4A**). We performed whole-genome sequencing of Br62 using both Illumina short reads and Nanopore long reads, followed by *de novo* whole-genome assembly (**Table S10**). Subsequent whole-genome alignment between AG006 and Br62 revealed that Br62_Contig07 corresponds to mChrA (AG006_Contig10) and AG006_Contig17 (**Fig 4B**). Furthermore, the alignment of the mChrA in both isolates revealed a high level of synteny, with a single re-arrangement in the center of mChrA (**Fig 4C**).

**Fig 4.**
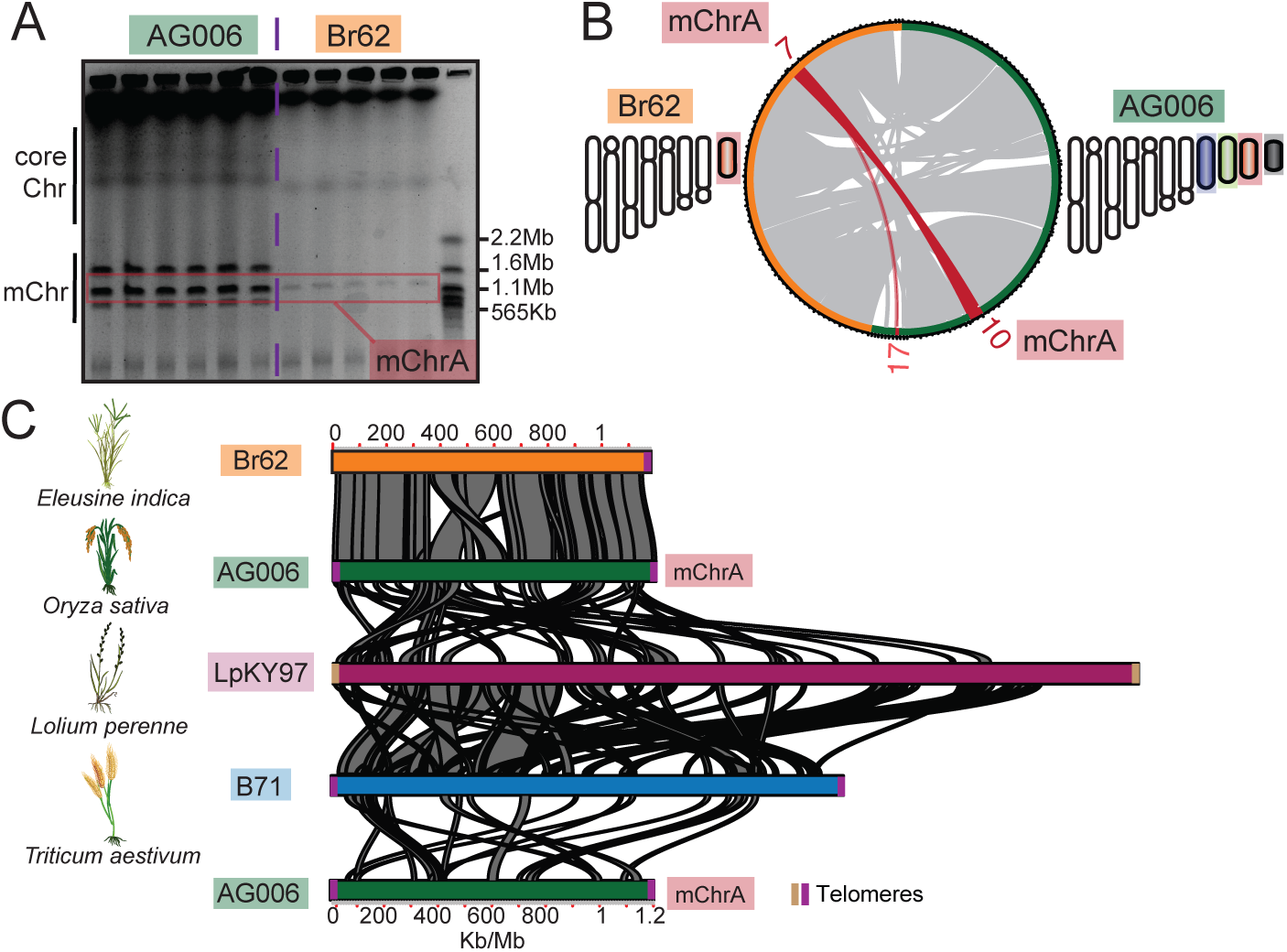
*Eleusine* isolate Br62 and *Oryza* isolate AG006 carry an intact and highly syntenic mChrA. **A.** CHEF-gel karyotyping of AG006 and Br62. Six gel lanes per isolate are shown representing a single biological replicate. Br62 carries a 1.2Mb mChr, the same size as mChrA in AG006 (in red). **B.** Whole-genome alignment of Br62 (orange) and AG006 (green). Br62_Contig07 aligns exclusively to mChrA (AG006_Contig10) and AG006_Contig17 (red). For both isolates a schematic karyotype is depicted. **C.** Alignment of mChrA in AG006 and Br62 reveal high synteny, except for a rearrangement in the central region. Alignments covering a fraction of mChrA are seen among mChrA in AG006 and the mChrA-like mChr1 in *Lolium* isolate LpKY97 (magenta) and the mChr in *Triticum* isolate B71 (blue). Telomeric sequences are indicated by vertical lines (purple/brown). The host plant of each isolate is shown on the left.

To independently validate these findings, we took advantage of a Br62 isolate that lost mChrA after subculturing, as determined by CHEF gel electrophoresis (**Fig S9A**). To identify contigs that originate from the mChr, we sequenced the genome of the Br62 isolate that lacks the 1.2Mb mChr (referred to as Br62-) using Illumina short-reads and aligned the reads to the Br62 genome. We calculated mapping depth per contig in Br62 and Br62-. Depths were consistent in both isolates except for Contig07, here Br62-displayed a near-zero read depth, indicating this corresponded to mChrA (**Fig S9B**). Additionally, Contig07 exhibited a high repeat content, a characteristic feature of mChr (**Fig S9C**). Together these analyses confirm the presence of an intact mChrA in Br62. Intriguingly, subculturing not only resulted in the loss of mChrA in Br62 but also in the loss of the mosaic mChrM in AG006 (**Fig 4A**), underlining the dynamic nature of mChr (Peng et al. 2019; Langner et al. 2021; Liu et al. 2022).

We next set out to determine whether the mChrA sequence is also found as an intact mChr in isolates belonging to other blast fungus lineages known to carry mChr (Peng et al. 2019; Rahnama et al. 2020), and which we identified as mChrA carriers (**Table S9**). We performed pairwise whole-genome alignments between AG006, isolate LpKY97 from the *Lolium* lineage, and isolate B71 from the *Triticum* lineage. Here, mChrA partially aligned to the mChr of both B71 and LpKY97, and to the end of chromosome 3 in B71, which was previously identified as a potential segmental duplication between the B71 mChr and core chromosomes (Peng et al. 2019; Liu et al. 2022; Gyawali et al. 2023) (**Fig 4C** and **Fig S10A**-**C**). The partial mChrA alignments are in accordance with our genetic clustering and breadth of coverage analyses, indicating that mChrA-like mChr are present in LpKY97 and B71, but these are structurally divergent from mChrA in AG006 and Br62 (**Table S6**). As a negative control, we aligned mChrA from AG006 to the conserved mChrC in the rice blast fungus isolate PR003 using the same parameters and no alignments were retrieved. We conclude that mChrA is present as an intact and highly syntenic mChr in the rice blast fungus isolate AG006 and in *Eleusine* blast fungus isolate Br62.

### Multiple horizontal mChrA transfers occurred in clonal rice blast fungus lineages

To test if sexual mating or horizontal gene transfer (HGT) can explain the presence of mChrA in the *Eleusine* and *Oryza* blast fungus lineages, we evaluated patterns of allele sharing through *D*-statistics (Green et al. 2010; Durand et al. 2011). After the mChrA sequence is removed from the genomes, we hypothesize that sexual mating results in a genome-wide introgression signal, leading to a *D*-statistic significantly different from zero, whereas HGT will not produce such a signal. Consequently, we first removed mChrA sequences, and then compared Br62 with the 32 rice blast fungus isolates carrying mChrA sequences, and 13 rice blast isolates not carrying this sequence (see Methods). For each comparison, rice blast fungus isolates belonging to the same *Oryza* subgroup were chosen. We selected *M. grisea* isolate Dig41 as an outgroup, which is divergent from both the rice and *Eleusine* blast fungus lineages. This resulted in the phylogenetic configuration: (Dig41, Br62; *Oryza* +mChrA, *Oryza* −mChrA). Under this configuration, a 99% confidence interval encompassing *D*=0 indicates there is no genome-wide introgression signal and favors the hypothesis of horizontal mChrA transfer. On the other hand, a 99% confidence interval not encompassing *D*=0 signals genome-wide introgression, supporting the acquisition of mChrA through sexual mating. In all tested configurations except those involving isolate BR0026 (31 of 32 isolates), the 99% confidence interval encompassed *D*=0, supporting the acquisition of mChrA by horizontal transfer (**Fig 5A** and **Table S11**). As a control, we tested the configurations (Dig41, Br62; *Oryza* +mChrA, *Oryza* +mChrA) (**Fig S11A**) and (Dig41, Br62; *Oryza* −mChrA, *Oryza* −mChrA) (**Fig S11B** and **Table S11**). Here, the 99% confidence interval encompassed *D*=0 in all tested configurations.

**Fig 5.**
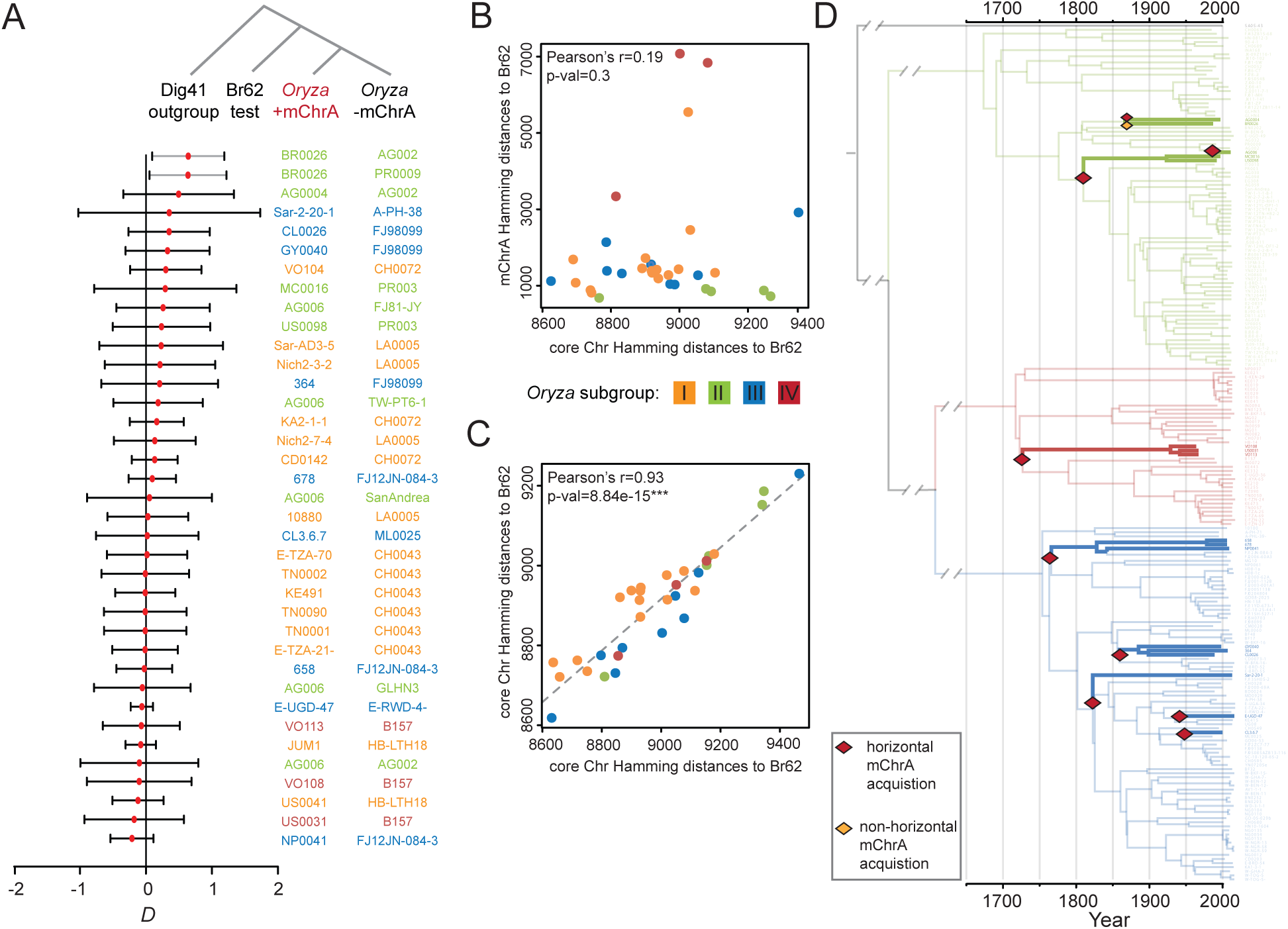
Multiple mChrA transfers occurred in clonal rice blast fungus lineages. **A.** *D*-statistics. Lines depict 99% confidence intervals and the red dot the estimated *D* value. Lines not encompassing *D*=0 are gray and the rest black. Jack-knife blocks were five million base pairs long. **B.** Hamming distances between mChrA and random core chromosomal regions of rice blast fungus isolates compared to Br62. **C.** Genetic distances between two sets of random core chromosomal regions in rice blast fungus isolates compared to Br62. **D.** Ancestral states of mChrA presence or absence along the clonal rice blast fungus phylogeny. Thick lines indicate mChrA is present. Branches are color-coded by lineage. The SA05-43 isolate from the *Setaria* blast fungus lineage was chosen as an outgroup. Branches with evidence for horizontal mChrA acquisition are indicated by a red diamond, the branch where there is evidence for sexual transfer or ILS is indicated by a yellow diamond.

Having established that mChrA was likely horizontally acquired in the large majority of rice blast fungus isolates carrying this sequence (31 of 32 isolates), and given its patchy distribution across the rice blast fungus lineage, we sought to differentiate between a single ancestral mChrA acquisition followed by independent losses, and multiple independent mChrA acquisitions. To test this, we estimated the genetic distance between the mChrA sequence present in the 32 *Oryza* isolates and Br62 and compared it to the genetic distance between random core chromosomal fragments of the same size as mChrA in the same 32 *Oryza* isolates (1.2Mb) and Br62. A significant correlation between the two genetic distances indicates that both the mChrA and the core chromosomes have accumulated mutations in a correlated way. This would support a single ancestral mChrA acquisition by the *Oryza* lineage, followed by multiple mChrA losses, whereas a lack of correlation suggests independent mChrA acquisitions. By analyzing correlations between genetic distances relative to Br62 instead of the magnitude of the distances, our analysis is not confounded by changes in mutation rate or different strengths of purifying selection operating at the mChrA and core chromosome level. We did not find any correlation in genetic distances between mChrA and Br62, and core chromosomes and Br62 (**Fig 5B**). As a control, we compared genetic distances to Br62 among two sets of core chromosomal regions and found a strong correlation (**Fig 5C**). Together, these results favor the hypothesis that the observed mChrA distribution in the rice blast fungus lineage is the result of multiple independent mChrA acquisitions.

In addition, we reconstructed the ancestral states of mChrA presence or absence along the clonal rice blast fungus phylogeny and found evidence for nine independent horizontal mChrA acquisitions. Using a time-scaled phylogeny, we could time these events to have occurred within the past three centuries (**Fig 5D**).

In summary, we provide compelling evidence supporting the scenario of multiple horizontal mChrA transfers involving members of the *Eleusine* and *Oryza* blast fungus lineages. A minimum of nine independent mChrA acquisitions and multiple independent losses occurred across clonal rice blast fungus lineages over the past three centuries.

## Discussion

Crop disease pandemics are frequently caused by clonal lineages of plant pathogens that reproduce asexually. The mechanisms enabling these clonal pathogens to adapt to their hosts, despite their limited genetic variation, remain an area of active research. In our study, we demonstrate that mini-chromosomes (mChr) serve as a source of genetic variation for asexual clonal pathogens. We observed horizontal mini-chromosome transfer occurred in field isolates belonging to clonal populations of the rice blast fungus *M. oryzae*. Our findings demonstrate horizontal acquisition of a 1.2Mb supernumerary mChr by clonal rice blast isolates from a genetically distinct lineage infecting *Eleusine indica*, a wild grass species. We identified a minimum of nine independent horizontal mChr acquisitions over the past three centuries. This establishes horizontal mChr transfer as a process facilitating genetic exchange between host-associated blast fungus lineages in the field. We propose that blast fungus populations infecting wild grasses serve as genetic reservoirs for clonal populations infecting cultivated crops. Horizontal acquisition of mChr by clonal blast fungus isolates appears to increase their genetic diversity, driving genome evolution and potentially aiding in its adaptability.

The genetic mechanisms underlying horizontal minichromosome (mChr) transfer in clonal fungus isolates are intriguing. Under laboratory conditions, horizontal transfer of mChr between fungal isolates has been facilitated through methods such as protoplast fusion (Akagi et al. 2009) or co-culturing (Masel et al. 1996; He et al. 1998; Ma et al. 2010; Vlaardingerbroek et al. 2016; van Dam et al. 2017). Underlying these mChr transfers is parasexual recombination (Soanes and Richards 2014; Vlaardingerbroek et al. 2016). Here, cells from different individuals fuse via anastomosis, forming heterokaryons (Roca et al. 2003; Roca et al. 2005; Ishikawa et al. 2010; Vangalis et al. 2021). These heterokaryons can become unstable polyploid cells, undergoing chromosome reassortment during mitosis (Mela et al. 2020). In the case of *Magnaporthe* spp. parasexual crosses do not exhibit heterokaryon incompatibility and are therefore viable (Crawford et al. 1986). This mechanism has been suggested as a source of genetic variation in the rice blast fungus, potentially occurring under field conditions (Zeigler et al. 1997; Noguchi et al. 2006; Tsujimoto Noguchi 2011; Monsur and Kusaba 2018). Here, we found robust evidence that horizontal mChr transfer occurs under field conditions, a process probably parasexual in nature.

Parasexuality offers fungi an alternative route to enhancing genetic diversity, while maintaining relative genomic stability and avoiding the complexities of sexual reproduction, including pre-mating barriers like reproductive timing and post-mating issues such as hybrid incompatibilities (Roper et al. 2011; Stukenbrock 2013). Recently, it was proposed that chromosome reassortment during parasexual recombination may not be entirely random (Habig et al. 2023). Here, it was suggested that some mChr are preferentially transferred or tend to resist degradation compared to others, resembling the behavior of selfish genetic elements (Ahmad and Martins 2019). This phenomenon could be attributed to distinct chromatin conformations of the mChr. Future research will investigate whether mChrA carries chromatin remodeling elements that could enable its horizontal transfer or shield it from degradation, potentially elucidating the relatively frequent horizontal transfer events observed across the rice blast fungus lineage. Moreover, to better understand the impact of horizontal mChr transfer on *M. oryzae* evolution, it will be crucial to study how frequent and diverse these events are in field populations.

Not all instances of inter-lineage transfer events of the mChrA sequence seem to be the product of parasexually-mediated horizontal transfer. Hybridization through sexual mating is a major player shaping the evolution of fungal plant pathogens, bringing forth a myriad of novel genetic combinations for selection pressures to act on (Stukenbrock 2016). In *M. oryzae*, there is evidence of sexual mating occurring both within and between specific host-associated blast fungus lineages, occasionally facilitating host jumps (Gladieux, Ravel, et al. 2018). In our study, one clonal rice blast fungus isolate, BR0026, exhibited genome-wide introgression signals with *Eleusine* isolate Br62. One plausible hypothesis is that the introgression signals observed in BR0026 may reflect ancient sexual reproduction events involving an isolate from the *Eleusine* lineage. Given that these two isolates were collected in South America, it is possible that sympatry in this region led to sexual reproduction between members of the rice and *Eleusine* lineages.

In addition to isolates belonging to the *Eleusine* and *Oryza* blast fungus lineages, isolates belonging to the *Triticum* and *Lolium* lineages also carry mChrA-like sequences. It remains to be determined whether these were obtained via horizontal transfer or sexual reproduction. The absence of genetic discordance between the core chromosomes and mChrA in these isolates supports sexual reproduction. In addition, substantial admixture has been observed among members of the *Triticum* and *Lolium* lineages (Gladieux, Ravel, et al. 2018), suggesting that sexual reproduction may be the route through which mChrA-like sequences were acquired.

Our study on the prevalence of horizontal gene exchange within local populations underscores the importance of accounting for ecological factors, especially in fungi that tend to specialize in specific hosts. Although our global comprehension of blast fungus populations has expanded, the detailed study of local populations, particularly those that include isolates from both wild and cultivated hosts, remain scarce (Cruz and Valent 2017; Barragan et al. 2022). One question to address will be how horizontal gene exchange through parasexuality is enabled in natural environments. In the case of the blast fungus, one factor offering an avenue for genetic interchange may be the absence of strict host-specialization (Gladieux, Condon, et al. 2018). Laboratory studies have shown that hosts like barley and common millet are susceptible to genetically distinct blast fungus lineages (Kato et al. 2000; Hyon et al. 2012; Chung et al. 2020). In the field, some cases of cross-infection have been reported, but the extent to which these occur in local populations is unknown (Gladieux, Condon, et al. 2018). Such susceptible hosts could serve as hubs for genetic exchanges, potentially contributing to horizontal mChr transfers between isolates from different lineages. Moreover, being a facultative biotroph, the blast fungus possesses the ability to thrive on both living plants and saprophytically on decaying plant matter. This broadens the window for possible genetic interactions, as the pathogen does not require synchronous growth within the same living hosts for this to occur. Understanding gene flow within local blast fungus populations through the study of horizontal gene transfer and other mechanisms, is vital for developing effective disease management strategies. For example, identifying frequent horizontal gene exchange between isolates infecting specific hosts could lead to targeted measures such as strategic weeding or focused fungicide application.

One persistent challenge in pinpointing elements of the accessory genome, such as mChr, has been the biases arising from aligning sequencing reads to a single reference genome. In past comparative genomic approaches, mChrA went unnoticed, as we aligned isolates carrying this sequence to the MG08 reference genome from isolate 70-15, which lacks the mChrA sequence. Leveraging pan-genomes, which are continuously gaining traction across the fungal kingdom (Badet and Croll 2020) or *de novo* assemblies using short read data (Potgieter et al. 2020), coupled with reference-independent genetic clustering approaches like k-mer (Zielezinski et al. 2017; Aylward et al. 2023) or read-based (Dylus et al. 2024) techniques, promises more accurate identification of mChr and of horizontally introgressed regions. The latter could be detected by first identifying mChr and other accessory genomic elements, and then comparing them with the core genome. Studies of this nature have recently detected cases of horizontal introgression in other fungal pathogens (Moolhuijzen et al. 2022; Petersen et al. 2023). In addition to these approaches, the integration of artificial intelligence to distinguish between core chromosomes and mChr using short-read sequencing data presents a timely and innovative approach (Gyawali et al. 2023). The successful implementation of such methodologies will not only facilitate the large-scale identification of candidate mChr regions across isolates, but also help establish whether these regions are preferentially involved in horizontal transfer events.

The horizontal transfer of mChrA from a blast fungus lineage that infects the wild grass *Eleusine indica* to clonal rice blast fungus lineages underscores the intricate ecological interactions involved. Wild grasses can act as potential genetic reservoirs, echoing the dynamics observed in zoonotic diseases where pathogens jump between wild animals and humans (Rahman et al. 2020). This analogy between the plant and animal realms highlights the significance of wild species as reservoirs of pathogens and suggests the possibility of genetic transfers. However, surveys on blast disease often focus on cultivated crops, neglecting wild hosts (Barragan et al. 2022). Therefore, enhanced awareness and surveillance of gene flow dynamics in local blast fungus populations are necessary. This should include investigations into the role of wild grasses as genetic conduits, similar to the concept of zoonoses. Such understanding is crucial for the early identification and prevention of genetic transfers that could initiate new disease outbreaks or intensify existing ones.

## Conclusion

Clonal isolates of the blast fungus are a significant agricultural concern due to their central role in causing crop disease pandemics. Key to tackling this issue is understanding how genetically uniform populations adapt to novel hosts. Our research has revealed that supernumerary mini-chromosomes undergo horizontal transfer in natural field conditions. Notably, we found that mChrA has been transferred horizontally on multiple independent occasions involving isolates from a lineage of blast fungus affecting a wild grass and clonal lineages infecting rice. This finding sheds light on the role of horizontal mini-chromosome transfer in driving the genome evolution of clonal blast fungus populations, potentially aiding in host adaptation. Isolates originating from wild grasses may act as reservoirs of genetic diversity (**Fig 6**). These insights underscore the importance of disease surveillance that encompasses both agricultural crops and adjacent wild grass species.

**Fig 6.**
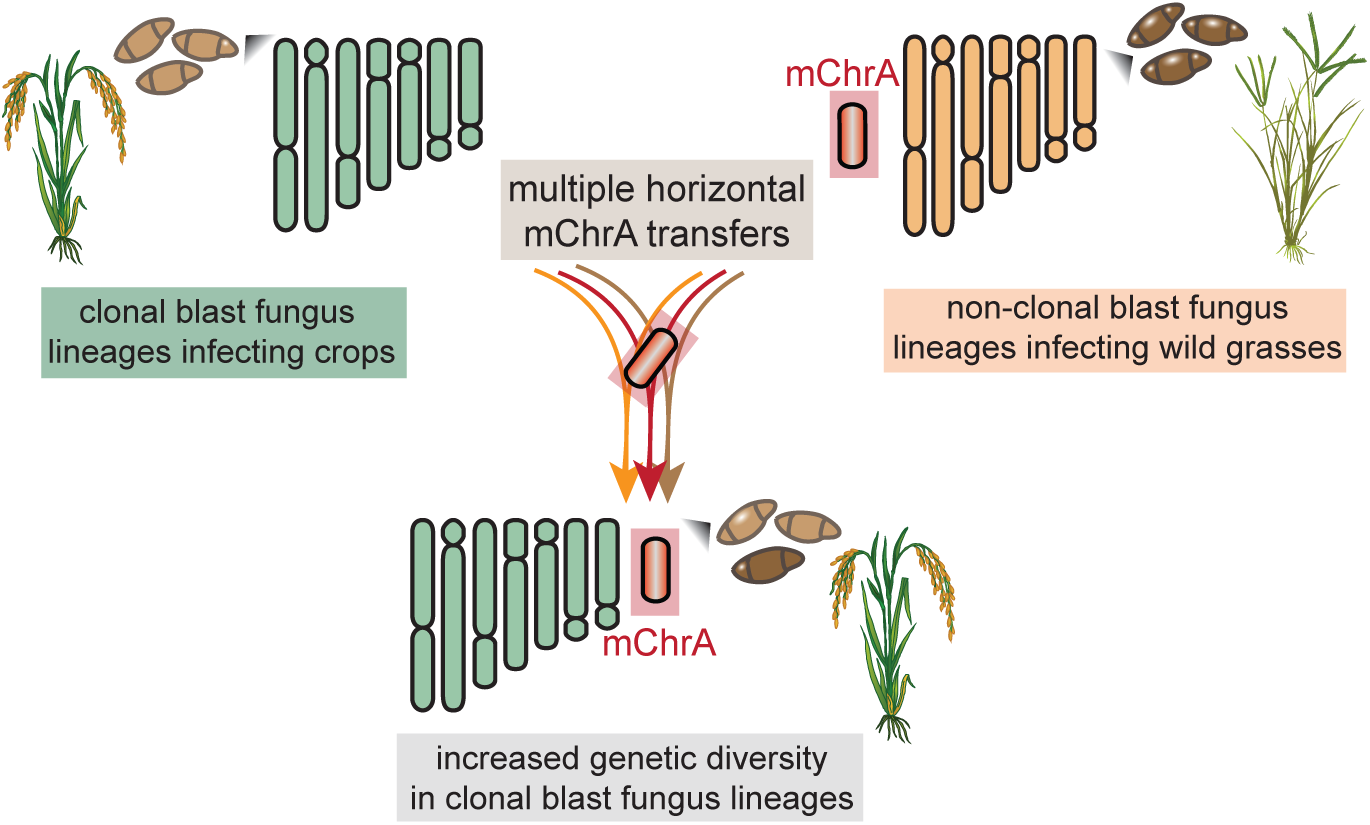
Horizontal mini-chromosome transfers from blast fungus lineages infecting wild grasses drive genome evolution of clonal lineages infecting crops. The recurrent acquisition of mChrA from wild grass-infecting blast fungus lineages by clonal rice blast fungus lineages enhances their evolutionary adaptability and capacity to respond to changing environments and hosts. The coexistence of infected crops and wild hosts facilitates this genetic exchange, posing a challenge to the management of crop disease pandemics.

### Model

## Materials and Methods

### Blast fungus growth conditions

Blast fungus isolates were grown from filter paper stocks by placing these on complete medium (CM) for 7-14 days in a growth chamber at 24°C with a 12 hour light period to induce growth of mycelium and sporulation. For liquid cultures, 8-10 small blocks of mycelium (ca. 0.5 × 0.5cm) were cut out of the edge of fully grown colonies with a sterile spatula, transferred into 150ml of liquid CM medium in a 250ml Erlenmeyer flask and incubated on a rotary shaker at 120 rpm and 24°C for 2-3 days.

### Visualization of worldwide blast fungus distribution

Maps showing the geographical locations of the studied blast fungus isolates were plotted with the R-package ggmap (v3.0) (Kahle and Wickham 2013). In the case of the San Andrea isolate, no exact collection coordinates were available, so the location of the San Andrea Chapel in Ravenna, in Italy’s Po Valley, the region where most other samples were collected from, was chosen.

### Whole-genome and mini-chromosome sequencing and genome assembly

Whole-genome sequencing and assembly of nine Italian blast fungus isolates, including AG006, is described in (Win et al. 2020). Briefly, these isolates were sequenced using the PromethION sequencing platform (Oxford Nanopore Technologies, Oxford, UK) and assembled into contigs using Canu (Koren et al. 2017). Assemblies were then polished with Illumina short reads using Pilon (Walker et al. 2014) and Racon (Vaser et al. 2017) and their completeness assessed using BUSCO (Simão et al. 2015), with a 97.7-98.8% completeness score taking the ascomycota_odb10 database as input (Win et al. 2020). Mini-chromosome isolation sequencing (MCIS) of these isolates was performed as described in (Langner et al. 2019; Langner et al. 2021). In short, mini-chromosomes (mChr) were separated from core chromosomes using CHEF gel electrophoresis. DNA was eluted from gel plugs and sequencing libraries were prepared using a modified version (custom barcodes) of the Nextera Flex library preparation kit (Illumina). Sequencing of mini-chromosomal DNA libraries was carried out on a NextSeq500 system (Illumina). For whole-genome sequencing and d*e novo* assembly generation of the Br62 isolate, high molecular weight DNA was extracted following (Jones et al. 2021). Sequencing runs were then performed by Future Genomics Technologies (Leiden, The Netherlands) using the PromethION sequencing platform (Oxford Nanopore Technologies, Oxford, UK). Long reads were assembled into contigs and corrected using Flye (v2.9-b17680) (Kolmogorov et al. 2019) and polished with long reads using Medaka (v1.7.2) (https://github.com/nanoporetech/medaka), and using Illumina short reads (San Diego, USA) through two consecutive iterations of Pilon (v1.23) (Walker et al. 2014). The resulting assembly was of high quality and contiguity, with a BUSCO (Simão et al. 2015) completeness score of 97.4% using the ascomycota_odb10 database and resulting in ten contigs (**Table S10**).

### Identification of mini-chromosomes in whole-genome assemblies

MCIS read quality was assessed using fastQC (Andrews 2010). Low quality and adapter sequences were removed using trimmomatic (Bolger et al. 2014). mChr reads were mapped to whole-genome assemblies of each strain using BWA-mem (Li 2013) with default parameters. Reads with multiple mappings (mapping quality = 0) and secondary alignments were removed using samtools (Danecek et al. 2021). MCIS read coverage was calculated in 1kb sliding windows with a step size of 500bp using bedtools (Quinlan and Hall 2010). The depth of unambiguously mapping reads was plotted using the R package circlize (Gu et al. 2014). To estimate the repeat content across core and mChr in the nine Italian rice blast isolates, we annotated these using RepeatMasker (http://www.repeatmasker.org/). The input repeat library consisted of the RepBase repeat library for fungi (https://www.girinst.org/repbase/), and repeat libraries from (Chiapello et al. 2015; Peng et al. 2019). For <2Mb contigs repeat content was plotted across 100kb sliding windows and a step size of 50kb, while for >2Mb contigs 10kb windows with a 5kb step size were chosen.

### Whole-genome and mini-chromosome alignments and telomere identification

Whole genome and contig-specific alignments between *M. oryzae* isolates were generated using the nucmer function of MUMMER4 (Marçais et al. 2018). Alignments of a minimum length of 10kb (-I 10000) and >80% percent identity (-i 80) were chosen to retrieve contiguous alignments. Alignment coordinates were extracted and whole genome alignments were plotted using the circlize package (Gu et al. 2014). Alignments between individual contigs were visualized with the karyoploteR package (Gel and Serra 2017). We visually inspected mChr contigs for the presence of (CCCTAA/TTAGGG)n canonical telomeric repeats (Cevernak et al. 2021).

### Genetic analysis of blast fungus isolates: mapping and variant calling

Illumina short reads of 413 *M. oryzae* and *M. grisea* isolates infecting different host plants (**Table S6**) were trimmed using AdapterRemoval (v2.3.1) (Schubert et al. 2016) and then mapped to the AG006 reference genome (Win et al. 2020) using bwa-mem (v0.7.17) (Li 2013) with default parameters. Variant identification was performed using GATK (v4.1.4.0) (McKenna et al. 2010). High-quality SNPs were filtered based on the Quality-by-Depth (QD) parameter using GATK’s VariantFiltration. Only biallelic SNPs within one standard deviation of the median value of QD scores across all SNPs were kept (Latorre et al. 2022). To study the phylogenetic relationship between isolates belonging to the rice blast fungus lineage, we subsetted 274 isolates belonging to this lineage (isolates BF5 and BTAr-A1 were removed due to them being outliers in the rice blast fungus phylogeny) and kept informative SNPs with no missing data using VCFtools (v0.1.14). From this dataset, we created a NeighborNet using Splitstree (Huson and Bryant 2006). We repeated this process for members of the *Oryza* clonal lineage II only, and constructed a Maximum-Likelihood (ML) tree using MEGA (v10.2.4) (Kumar et al. 2018), with 100 bootstraps (see data availability). We repeated the same process for the analysis of all 413 isolates shown in Fig 1 (**Table S6**). Here, two isolates were removed due to the high amount of missing sites (FR13 and 98-06). Based on these SNPs, we created a NJ tree using MEGA (v10.2.4) (Kumar et al. 2018), with 100 bootstraps (see data availability). Isolates deemed as mChrA carriers were highlighted using iTol (Letunic and Bork 2021). To assess for potential discordance in genetic clustering of the core genome and mChrA, we subsetted isolates carrying mChrA (n=126). For both the core genome and mChrA, only SNPs with a maximum of 10% missing data were kept (--max-missing 0.9) using VCFtools (v0.1.11). NJ trees were constructed using IQtree (v2.03) using fast mode (see data availability). SNP-based Principal Component Analyses (PCA) were estimated using the --pca function of PLINK2 (Chang et al. 2015). These were visualized using the R package ggplot2 (v3.4.4, see data availability) (Wickham 2009). To determine the likelihood of the observed genetic discordance being observed by chance, 100 random 1.2Mb regions across the core genome in these 126 isolates were subsetted using a custom python script (see data availability), and NJ trees were computed using IQTree (v2.03) with the fast mode. The number of times each lineage was monophyletic was estimated using a provided custom python script (see data availability). To evaluate genetic differentiation between members of the *Eleusine* and *Oryza* lineages carrying the mChrA sequence, the fixation index (F*_ST_*) based on genome-wide SNPs was calculated. Rice blast isolates carrying mChrA (n=32) were compared to the two *Eleusine* isolates carrying mChrA, Br62 and B51, using only SNPs with no missing data. Weighted F*_ST_* using 5kb window sizes and 500bp step sizes (--fst-window-size 5000 --fst-step-size 500) was calculated using VCFtools (v0.1.14).

### mChrC and mChrA breadth of coverage calculations and mChrA-carrier assignment

To investigate the distribution of the mChrC and mChrA sequence across 413 *M. oryzae* and *M. grisea* isolates, we first calculated the genome-wide breadth of coverage, defined as the percentage of sequence covered by reads from a particular isolate which mapped to the AG006 reference (**Table S6**). To do this, we estimated breadth of coverage per contig using samtools depth (v1.19) (Danecek et al. 2021), and then created a weighted average taking into account contig length. We assessed breadth of coverage for mChrC (AG006_Contig03) and mChrA (AG006_Contig10) across all isolates and then normalized these values by the isolate’s genome-wide breadth of coverage value (**Table S6**). To determine whether an isolate carried the mChrA sequence or not, we performed clustering using a gaussian mixture model (GMM) and estimated the Bayesian Information Criterion (BIC) value for 1-10 clusters using the R-package mclust (v.6.0.0, see data availability) (Scrucca et al. 2023). Using this same package, we also estimated the uncertainty index for mChrA presence (n=126) or absence (n=287) assignment for each isolate (**Table S9**).

### Identification of mChrA in *Eleusine* isolate Br62 and mChrA loss Br62-

*M. oryzae* isolate Br62, belonging to the *Eleusine* lineage, initially carried a single mChr identical in size to mChrA (1.2Mb). To confirm the identity of this mChr, we subcultured Br62 twice via serial passage on Complete Growth Medium (CM), resulting in the loss of mChr as confirmed through CHEF gel electrophoresis. We then sequenced the genome of Br62 without the 1.2Mb mChr (referred to as Br62-) using Illumina short-reads and compared it to the complete Br62 genome sequences. Mapping depth per contig was calculated using the samtools depth function (Danecek et al. 2021). Depths were consistent between Br62 and Br62-except for Contig07, corresponding to mChrA, where Br62-displayed a near-zero read depth. Additionally, repeat content analysis for the Br62 genome, using the same parameters as for the Italian rice blast fungus isolates.

### Differentiation between horizontal mChrA transfer from introgression via sexual mating

To differentiate between horizontal mChr transfer or sexual mating we assessed patterns of allele sharing and calculated *D* statistics (Green et al. 2010; Durand et al. 2011) using popstats (Skoglund et al. 2015) as well as using the custom python script *Dstat.py* (see data availability). We removed the mChrA sequence from the *Eleusine* and *Oryza* mChrA carriers and set the *M. grisea* isolate Dig41 as an outgroup, resulting in the following 4-taxa configuration: (Dig41, Br62; *Oryza* +mChrA, *Oryza* −mChrA). The selection of the non-carrier samples (-mChrA) was random and contingent on their phylogenetic proximity to the tested mChrA carrier (+mChrA) isolate (**Table S11**). In the case of +mChrA isolate AG006, we performed comparisons against 13 different *Oryza* −mChrA isolates, selected throughout along the different clades of the clonal lineage II. As a control, we also tested the 4-taxa configuration: (Dig41, Br62; *Oryza* +mChrA, *Oryza* +mChrA). The tested isolates were selected based on them having phylogenetic proximity. Complementary to this, we included a second control using the 4-taxa configuration: (Dig41, Br62; *Oryza*-mChrA, *Oryza* −mChrA). The testing pair of isolates were chosen randomly and contingent on being part of the same genetic subgroup of the rice blast fungus lineage. In all tested configurations, we only compared rice blast fungus isolates belonging to the same subgroup, to avoid potential unequal drift accumulated between members of different clonal lineages from impacting the analysis. For each configuration we calculated the 99% confidence interval. *D* values were estimated for jack-knife blocks 5 and 10 million base pairs in length (**Table S11**).

### Differentiating between a single and multiple horizontal mChrA transfer events

To differentiate between a single ancestral gain of mChrA followed by independent losses, and independent mChrA gains. We measured Hamming distances between mChrA in all *Oryza* isolates carrying this sequence (n=32) and *Eleusine* isolate Br62, and Hamming distances across random core chromosomal regions (the size of mChrA, 1.2Mb) and Br62 using bcftools (v.1.11) (Danecek et al. 2021), (see data availability). We then assessed if there is correlation between the two hamming distances. Both Pearson’s correlation coefficient and its *p*-value were estimated. As a control, we compared average Hamming distances between two sets of core chromosomal regions to Br62 and again calculated Pearson’s correlation coefficient and its *p*-value.

### Dating of horizontal mChrA transfer Events Across Clonal Rice Blast Fungus Lineages

In order to infer the dating times of horizontal acquisition of the mChrA sequence in the ancestral nodes of the rice blast fungus phylogeny, we performed a bayesian-based dated phylogeny incorporating the isolate collection dates (**Table S6**) using BEAST2 (Bouckaert et al. 2014). We selected the Hasegawa-Kishino-Yano (HKY) nucleotide substitution model. The collection years of the blast fungus isolates served as prior information, providing expected units for the estimated evolutionary rate (substitutions/site/year). In Bayesian analysis, we utilized a log-normal distribution with a mean in real space set at 7.5E-8, based on previous estimations (Latorre et al. 2022). To minimize the effect of demographic assumptions, we chose a Coalescent Extended Bayesian Skyline as a tree prior (Drummond et al. 2005). Isolates without a known collection date were removed from this analysis, and only individuals belonging to rice blast clonal lineages were used to rule out recombination. We ran six independent chains, each spanning a length of 20 million iterations using the CIPRES infrastructure (Miller et al. 2010). To ascertain the ancestral states of presence or absence of the mChrA sequence throughout the rice blast fungus phylogeny, we used the inferred mChrA presence/absence information based on breadth of coverage analyses (**Table S9**). This was done for all rice blast fungus isolates, as well as for the SA05-43 isolate which belongs to the *Setaria* blast fungus lineage, which was set as an outgroup. These values, which were input as discrete states (mChrA = yes/no), were parameterized in a “mugration” analysis, which was implemented in Treetime (v.0.9.0) (Sagulenko et al. 2018) ML-tree as input, generated using IQtree (v2.03) (Minh et al. 2020).

### Textual enhancement

The articulation of text within this manuscript was assisted by the machine learning model ChatGPT-4.

## Supporting information

Barragan_2024_mChrA_SupplementaryData

## Acknowledgements

We thank all members of the Kamoun laboratory and the BLASTOFF team at the Sainsbury Laboratory for valuable discussions. We especially thank Vincent Were for valuable suggestions. We also thank Ana Maria Picco for providing rice blast fungus isolates from Italy, and Alison MacFadyen for managing the public release of sequencing data.

## Data availability

The authors confirm that all data underlying the findings are fully available without restriction. Files and code to perform the analyses described and to generate the plots presented are available as Supplementary Files, and in the git repository: https://github.com/smlatorreo/mChr_Moryzae (Latorre 2024). Sequencing reads were deposited in the European Nucleotide Archive (ENA) under study accession number PRJEB6623 (mini-chromosome sequences from Italian rice blast isolates) and PRJEB67435 (Br62-sequencing). In addition, the Br62 whole-genome assembly is available under GenBank accession number PRJEB66723.

## Author Contributions

**Conceptualization:** ACB, SML, HAB, SK, TL.

**Formal analysis:** ACB, SML, AM, AH, JW, TL.

**Investigation:** ACB, SML, AM, AH, JW, YS, TL.

**Visualization:** ACB, SML, AM, TL.

**Coding:** ACB, SML, AM, TL.

**Supervision:** HAB, SK.

**Writing – original draft:** ACB.

**Writing – review & editing:** ACB, SML, HAB, SK and TL with contributions from all authors.

**Project administration:** ACB, SK.

**Funding acquisition:** ACB, HAB, SK, TL.

**Competing interests**. The authors have declared that no competing interests exist.

