## Supplementary material for "Multiple horizontal mini-chromosome transfers drive genome evolution of clonal blast fungus lineages": Barragan_2024_mChrA_SupplementaryData: Barragan_2024_SupplementaryFigures.pdf

**Fig S1.** Rice blast fungus isolates collected from Italy belong to a single clonal lineage. Related to Fig 1.

**Fig S2.** Identification and analysis of mChr in clonal rice blast fungus isolates. Related to Fig 1.

**Fig S3.** mChr contigs in clonal rice blast fungus isolates display high MCIS coverage and repeat content. Related to Fig 1.

**Fig S4.** mChrC distribution across the global *M. oryzae* and *M. grisea* population. Related to Fig 1.

**Fig S5.** Alignment of mChrA-like contigs from rice blast fungus isolate AG006. Related to Fig 1.

**Fig S6.** mChrA sequences are present across multiple host-associated blast fungus lineages. Related to Fig 2.

**Fig S7.** Principal Components for the core genome and mChrA comparisons. Related to Fig 3.

**Fig S8.** mChrA shows less genome-wide differentiation than the rest of the genome between isolates from the *Eleusine* and *Oryza* blast fungus lineages. Related to Fig 3.

**Fig S9.** Identification of mChrA in *Eleusine* isolate Br62. Related to Fig 4.

**Fig S10.** mChrA in AG006 partially aligns to mChr in isolates LpKY97 and B71. Related to Fig 4.

**Fig S11**. Lack of sexual reproduction or ILS signals between members of the *Eleusine* and *Oryza* lineages. Related to Fig 5.

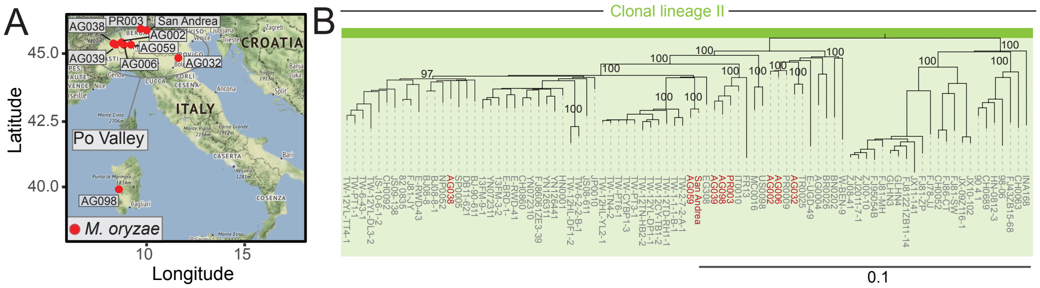


**Fig S1. Rice blast fungus isolates collected from Italy belong to a single clonal lineage. A**. Names and sampling locations of the nine isolates studied (red). **B.** A genome-wide SNP-based Maximum-likelihood (ML) tree confirms the Italian isolates (in red) belong to rice blast fungus clonal lineage II (Latorre et al. 2020). AG006 and AG002 are closely related to each other. Relevant bootstrapping values are shown. Scale bar indicates nucleotide substitutions per position. Related to Fig 1.

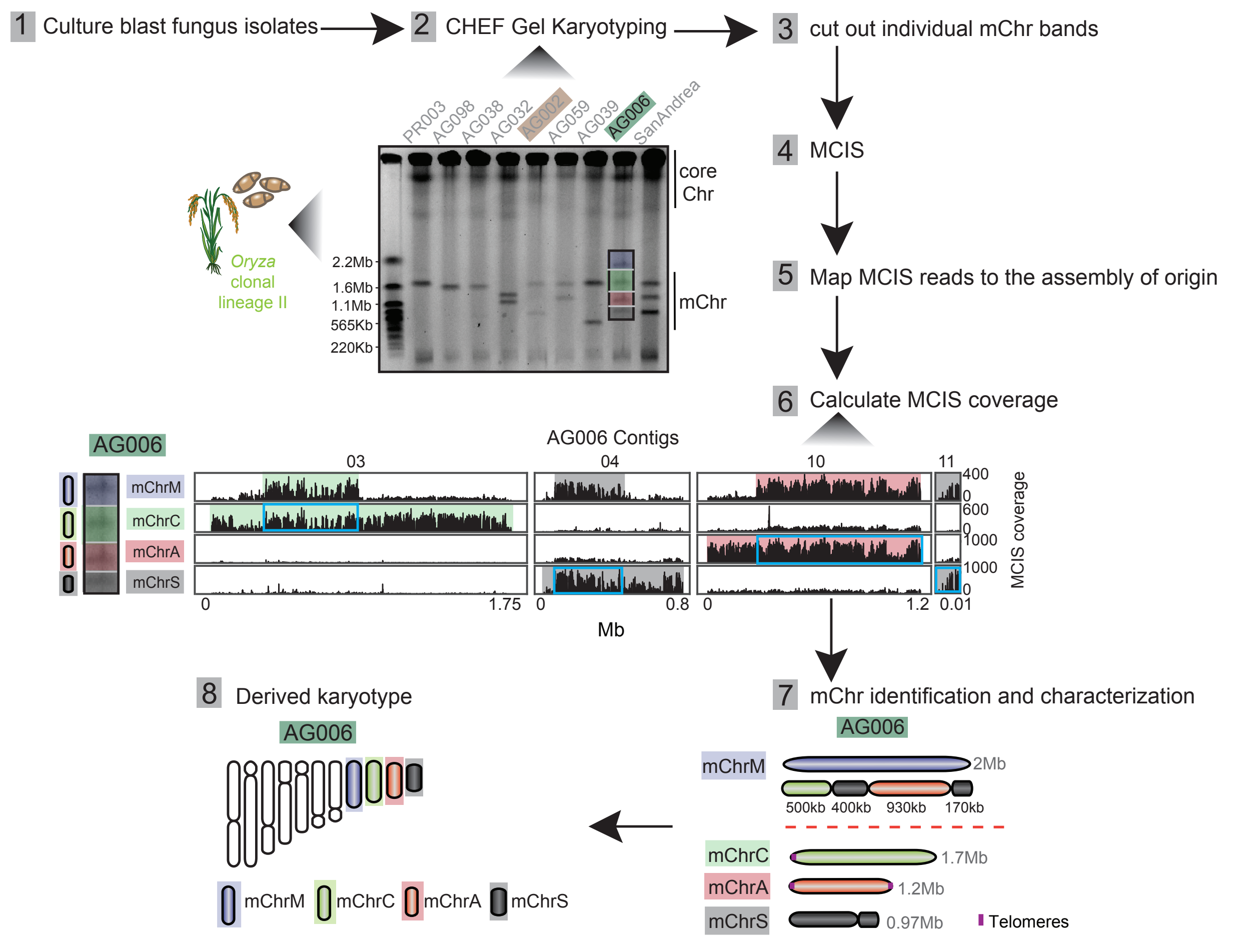


**Fig S2. Identification and analysis of mChr in clonal rice blast fungus isolates.** Nine clonal rice blast fungus isolates were cultured (1). Their mChr content was visualized though CHEF gel-electrophoretic karyotyping (2). To assess the genetic composition of individual mChr, these were excised from the gel individually (3) and sequenced (mini-chromosome isolation sequencing (MCIS), 4). The resulting mChr reads were mapped to the genome assembly of the isolate they originate from (5). MCIS coverage was determined (6, see Fig S3). High MCIS coverage indicates mChr contigs. For example, high MCIS coverage across sections of various mChr contigs (green, gray and red) in AG006 revealed the mosaic nature of mChrM (6 and 7, see Fig 1E). MCIS coverage analysis allows for a detailed representation of mChr content per isolate, as shown for isolate AG006. Regions in mChrC, mChrA and mChrS with overlapping sequences to mChrM are highlighted in light blue. Telomeric sequences are indicated by a vertical line (purple). As a result of this workflow, the karyotype of each *M. oryzae* isolate can be derived (8). Related to Fig 1.

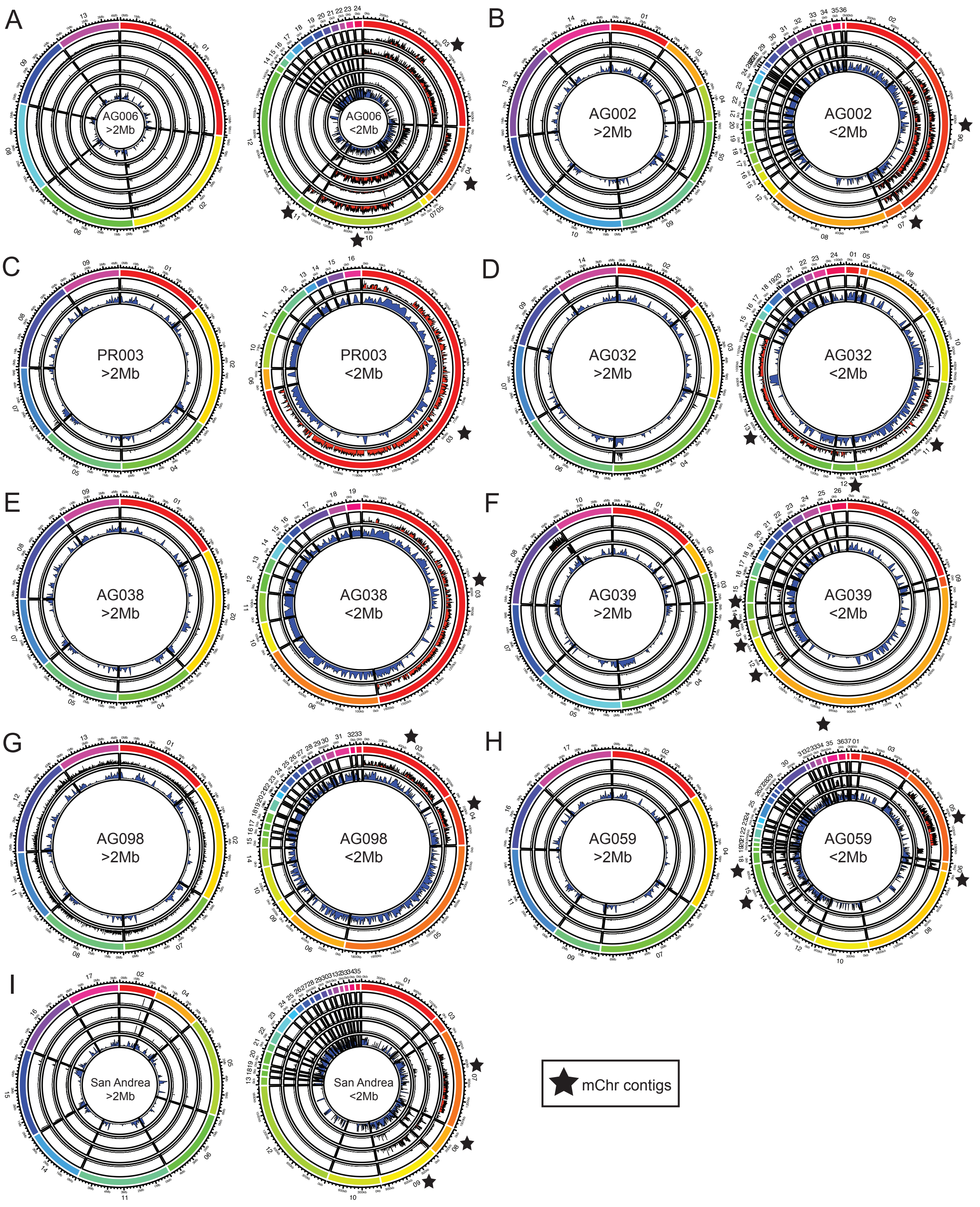


**Fig S3. mChr contigs in clonal rice blast fungus isolates display high MCIS coverage and repeat content. A-I.** Circos plots of mini-chromosome isolation sequencing (MCIS) read coverage and repeat content across nine rice blast fungus isolates. For each isolate, >2Mb contigs are on the left and <2Mb contigs on the right. The outer ring indicates the contigs of each assembly (rainbow colors). mChr contigs are indicated by black stars. Outer track(s) (black/red) indicates MCIS coverage per 10kb sliding window and a step size of 5kb, each track represents a different mChr extracted from an individual gel band. The inner track (black/blue) indicates repeat content. For >2Mb contigs, 100kb sliding windows and a step size of 50kb are shown, and for <2Mb contigs, 10kb windows with a 5kb step size are shown. See supplemental data for individual plots. Related to Fig 1.

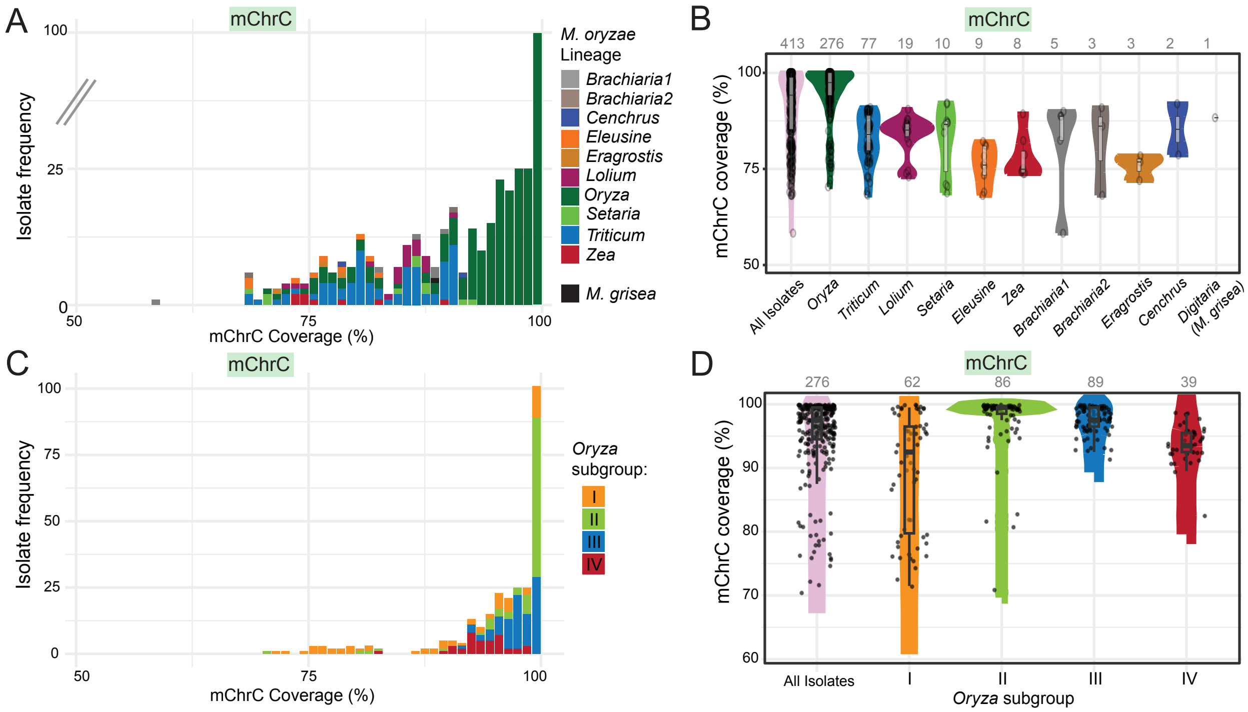


**Fig S4. mChrC distribution across the global *M. oryzae* and *M. grisea* population. A-B**. mChrC (AG006_Contig03) breadth of coverage across 413 *M. oryzae* and *M. grisea* isolates belonging to different host-associated lineages (Table S6). Each isolate is denoted by a black dot and numbers at the top indicate the number of isolates per lineage (B). mChrC is particularly conserved across isolates from the *Oryza* lineage. **C-D.** mChrC breadth of coverage across rice blast fungus isolates. mChrC is particularly prevalent in isolates belonging to the clonal lineage II (green). Each isolate is denoted by a black dot and numbers at the top indicate the number of isolates per lineage (D). Related to Fig 1.

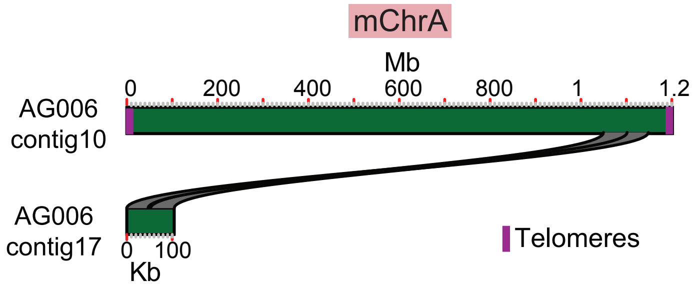


**Fig S5. Alignment of mChrA-like contigs from rice blast fungus isolate AG006.** AG006_Contig17 shares sequence similarity with AG006_Contig10 (mChrA), which displays telomeric sequences at both ends, indicated by a vertical line (purple). Related to Fig 1.

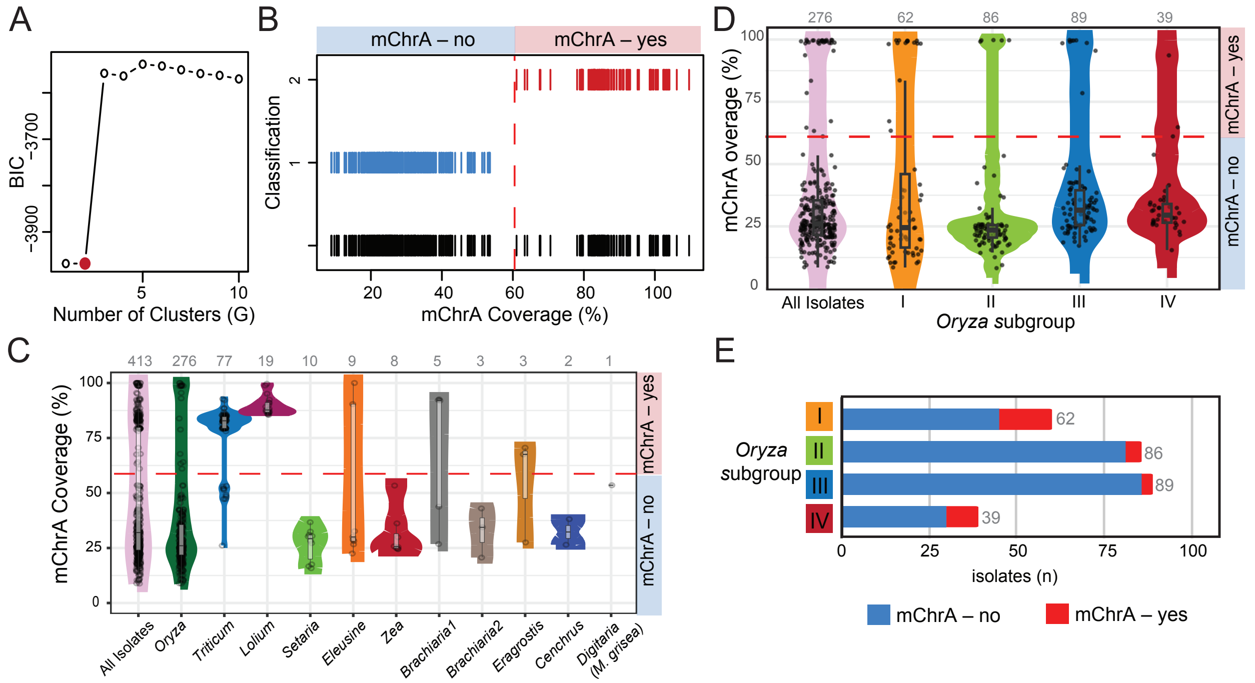


**Fig S6. mChrA sequences are present across multiple host-associated blast fungus lineages. A.** Bayesian Information Criterion (BIC) values for different cluster numbers (G=1-10) show that G=2, marked in red, has a low value, signifying a good fit for the data. **B.** Assignment of isolates as mChrA=yes/no for G=2 using a gaussian mixture model (GMM). The mChrA coverage cutoff (61%) is indicated by the dotted red line. **C.** mChrA breadth of coverage across *M. oryzae* lineages and *M. grisea*. Numbers on top indicate the number of isolates belonging to each lineage, with each isolate denoted by a black dot. **D.** mChrA coverage distribution across the rice blast fungus lineage. Numbers are black dots as in C. **E.** Proportion of isolates carrying mChrA (red) or not (blue) in each genetic subgroup of the rice blast fungus lineage. Numbers next to each bar indicate the isolates belonging to each subgroup. Related to Fig 2.

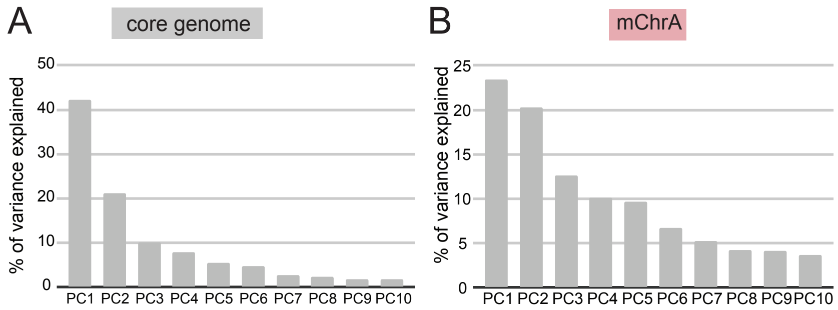


**Fig S7. Principal Components for the core genome and mChrA comparisons. A-B.** SNP-based Principal Component Analyses (PCA). Percentage of variance explained by each individual principal component (PC1-10), for the core genome (A) and mChrA (B). For both cases, PC1 and PC2 are plotted in Fig 3. Related to Fig 3.

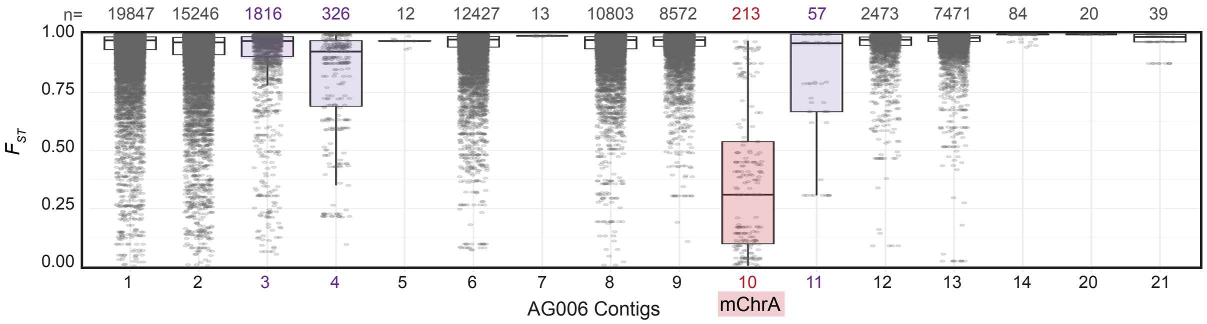


**Fig S8. mChrA shows less genome-wide differentiation than the rest of the genome between isolates from the *Eleusine* and *Oryza* blast fungus lineages. A.** $F_{ST}$ between rice (n=32) and *Eleusine* (Br62 and B51) blast fungus isolates carrying mChrA (AG006_Contig10, in red). mChrA exhibits lower genetic differentiation compared to the rest of the genome. Other mChr contigs in violet. Each dot represents the weighted $F_{ST}$ per 5kb window using a step size of 500bp. The number of windows per contig present at the top of each bar. All informative contigs are shown. Related to Fig 3.

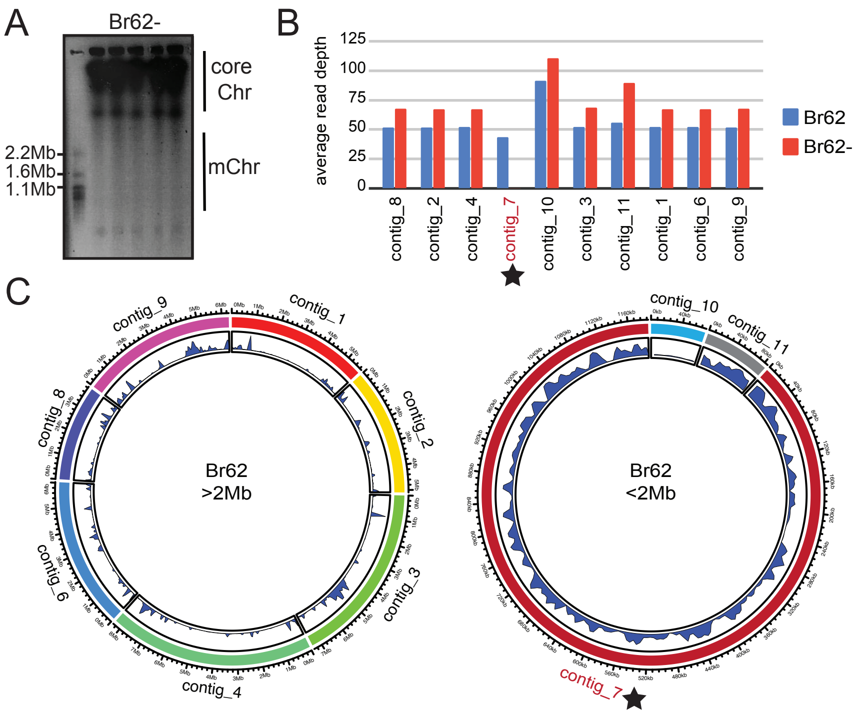


**Fig S9. Identification of mChrA in *Eleusine* isolate Br62. A.** CHEF-gel karyotyping of Br62-, the Br62 isolate without its 1.2Mb mChr. Five gel lanes were dedicated to this isolate, representing one biological replicate. **B.** Illumina whole-genome read sequencing depth per contig of both the original Br62 isolate carrying mChrA, and the Br62 isolate that lost mChrA (Br62-). Br62_Contig07, the mChrA contig, has much lower depth (0.04x) in Br62- than it has in Br62 (43.57x). **C**. Circos plots of repeat content in Br62, >2Mb contigs are on the left and <2Mb contigs on the right. The inner track (black/blue) indicates repeat content. For >2Mb contigs 100kb sliding windows and a step size of 50kb are shown. For <2Mb contigs 10kb windows with a 5kb step size are shown. Contig07 displays high repeat content and is indicated by a black star. Related to Fig 4.

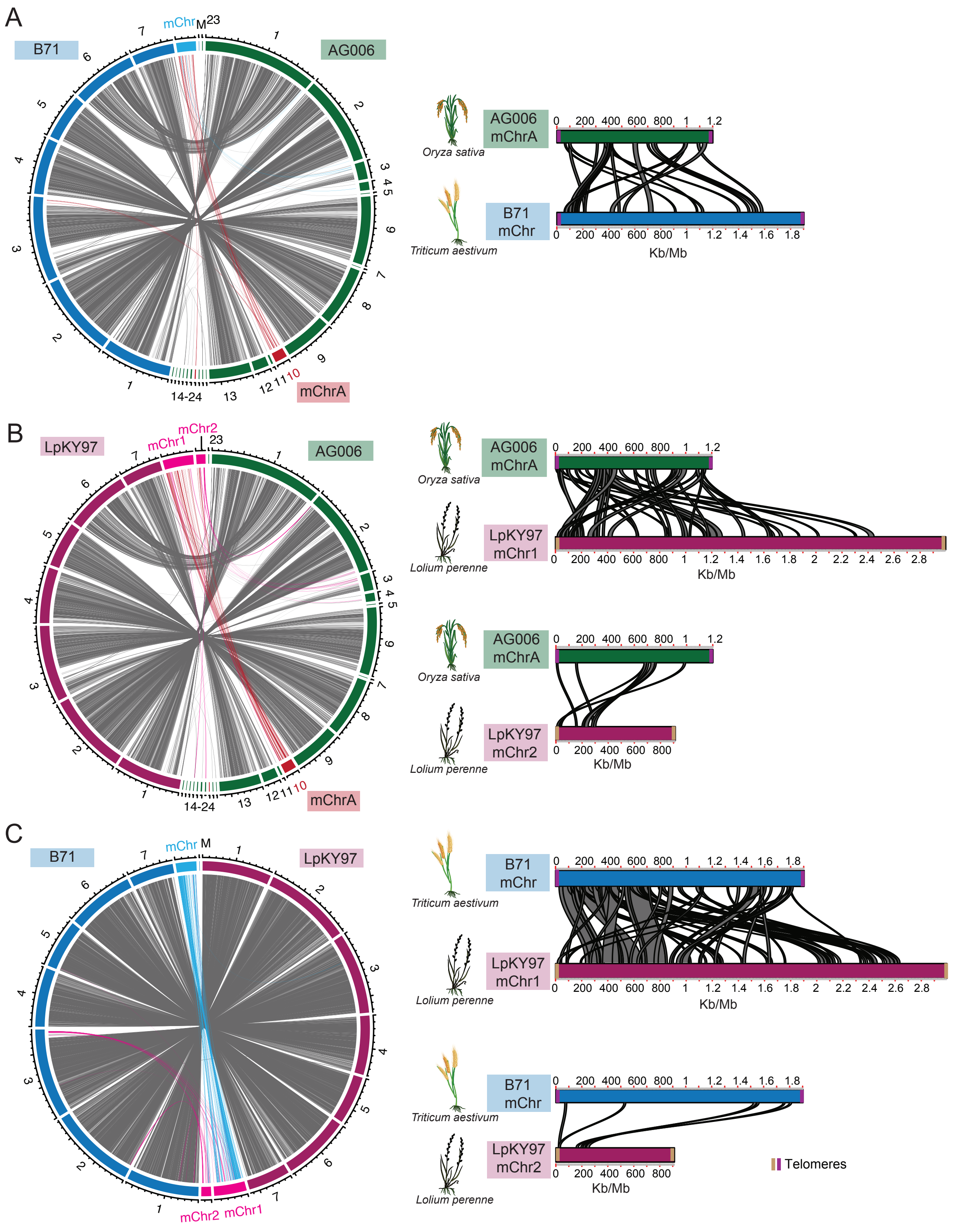


**Fig S10. mChrA in AG006 partially aligns to mChr in isolates LpKY97 and B71. A.** Whole-genome alignment between *Triticum* isolate B71 (blue) and AG006 (green). Each number represents a contig. Red lines indicate alignments from AG006 mChrA to other genomic regions, while blue alignments indicate B71 mChr alignments to regions other than mChrA. **B.** Whole-genome alignment between *Lolium* isolate LpKY97 (magenta) and AG006 (green). Red lines as in A, while pink alignments indicate LpKY97 mChr1 and mChr2 alignments to regions other than mChrA. **C.** Whole-genome alignment of B71 (blue) and LpKY97 (magenta). Blue lines indicate alignments from B71 mChr to mChr1 and mChr2 in LpKY97, while pink indicates alignments from these two mChr to regions other than B71 mChr. Telomeric sequences are indicated by vertical lines (purple/brown). Related to Fig 4.

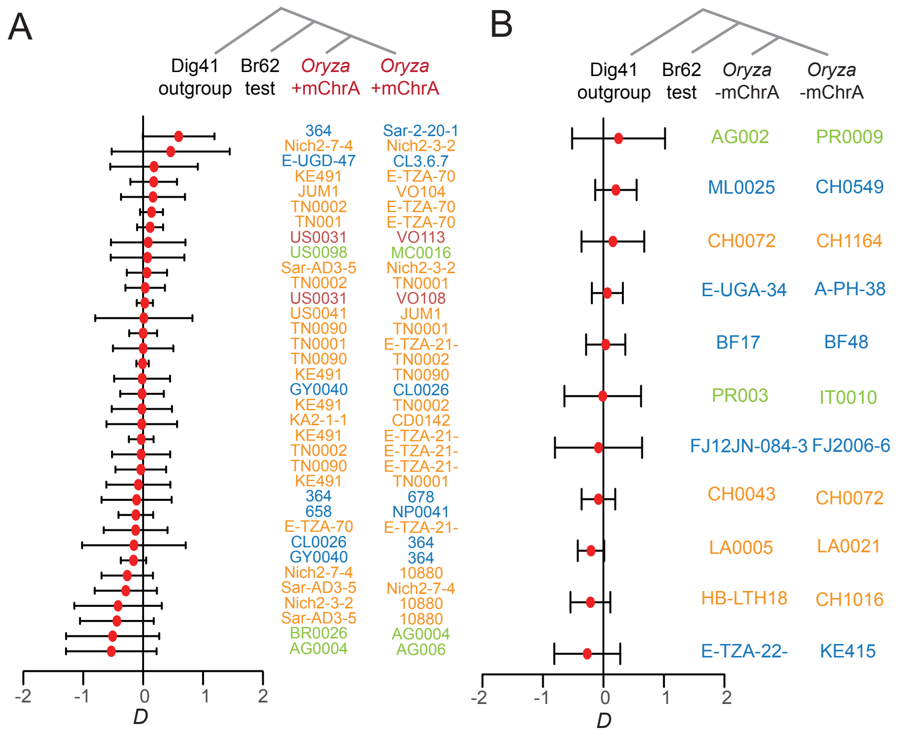


**Fig S11. Lack of sexual reproduction signals between members of the *Eleusine* and *Oryza* lineages. A-B.** *D*-statistics. The mChrA sequence was removed from all isolates carrying mChrA. *M. grisea* isolate Dig41 was set as an outgroup. The resulting phylogenetic configuration was: (Dig41, Br62 ; *Oryza* +mChrA, *Oryza* +mChrA, A) and (Dig41, Br62 ; *Oryza* -mChrA, *Oryza* -mChrA, B). Jack-knife blocks were 5 million base pairs long. In all configurations, D=0 was encompassed in the 99% confidence interval. Related to Fig 5.
