## Supplementary material for "Multiple horizontal mini-chromosome transfers drive genome evolution of clonal blast fungus lineages": Barragan_2024_mChrA_SupplementaryData: riceblast_trees.sh

```
###############################################################################
######### Create genome-wide SNP based NeighborNets or NJ trees from a VCF
########## written by Cristina Barragan 2024 ###################################

#### Fig1A
#subset rice blast fungus individuals from Master VCF
vcftools --vcf /hpc-home/barragan/Italian_blast/pangenome.snps.filtered.bi.recode.vcf --keep 274_ind_rice.txt --recode --recode-INFO-all --out  pangenome.snps.filtered.bi.274rice
#keep only SNPs with no missing data
vcftools --vcf pangenome.snps.filtered.bi.274rice.recode.vcf --max-missing 1.0 --recode --recode-INFO-all --out pangenome.snps.filtered.bi.274rice.md0
#edit vcf
vcf=pangenome.snps.filtered.bi.274rice.md0.recode.vcf

sed -i 's/Contig01/1/g' $vcf
sed -i 's/Contig02/2/g' $vcf
sed -i 's/Contig03/3/g' $vcf
sed -i 's/Contig04/4/g' $vcf
sed -i 's/Contig05/5/g' $vcf
sed -i 's/Contig06/6/g' $vcf
sed -i 's/Contig07/7/g' $vcf
sed -i 's/Contig08/8/g' $vcf
sed -i 's/Contig09/9/g' $vcf
sed -i 's/Contig10/10/g' $vcf
sed -i 's/Contig11/11/g' $vcf
sed -i 's/Contig12/12/g' $vcf
sed -i 's/Contig13/13/g' $vcf
sed -i 's/Contig14/14/g' $vcf
sed -i 's/Contig15/15/g' $vcf
sed -i 's/Contig16/16/g' $vcf
sed -i 's/Contig17/17/g' $vcf
sed -i 's/Contig18/18/g' $vcf
sed -i 's/Contig19/19/g' $vcf
sed -i 's/Contig20/20/g' $vcf
sed -i 's/Contig21/21/g' $vcf
sed -i 's/Contig22/22/g' $vcf
sed -i 's/Contig23/23/g' $vcf
sed -i 's/Contig24/24/g' $vcf

# Create an artificial diplots genome: transform vcf to plink and edit vcf
#vcftools --vcf pangenome.snps.filtered.bi.274rice.md0.recode.vcf --plink --out pangenome.snps.filtered.bi.274rice.md0
#plink2 --vcf  pangenome.snps.filtered.bi.274rice.md0.recode.vcf  --out  pangenome.snps.filtered.bi.274rice.md0.2
#plink2 --pfile pangenome.snps.filtered.bi.274rice.md0.2 --recode vcf --out pangenome.snps.filtered.bi.274rice.md0.2.plink
#sed -i 's/^##fileformat=VCFv4.3/##fileformat=VCFv4.2/' pangenome.snps.filtered.bi.274rice.md0.2.plink.vcf

#prune SNPs
vcftools --vcf pangenome.snps.filtered.bi.274rice.md0.recode.vcf --plink --out pangenome.snps.filtered.bi.274rice.md0
plink --file pangenome.snps.filtered.bi.274rice.md0 --indep-pairwise 50 5 0.5 --out pangenome.snps.filtered.bi.274rice.md0.prune50.5.05
sed -i 's/:/   /g' pangenome.snps.filtered.bi.274rice.md0.prune50.5.05.prune.in
vcftools --vcf pangenome.snps.filtered.bi.274rice.md0.recode.vcf --positions pangenome.snps.filtered.bi.274rice.md0.prune50.5.05.prune.in --recode --recode-INFO-all --out pangenome.snps.filtered.bi.274rice.md0.prune50.5.05

#output file
#pangenome.snps.filtered.bi.274rice.md0.prune50.5.05.recode.vcf 
#kept 10,728 out of a possible 354,307 Sites after pruning

#transform from vcf to fasta
bgzip -c  pangenome.snps.filtered.bi.274rice.md0.prune50.5.05.recode.vcf >  pangenome.snps.filtered.bi.274rice.md0.prune50.5.05.recode.vcf.gz
tabix -p vcf  pangenome.snps.filtered.bi.274rice.md0.prune50.5.05.recode.vcf.gz
zcat pangenome.snps.filtered.bi.274rice.md0.prune50.5.05.recode.vcf.gz | vcf-to-tab > pangenome.snps.filtered.bi.274rice.md0.prune50.5.05.tab
perl  vcf_tab_to_fasta_alignment.pl -i pangenome.snps.filtered.bi.274rice.md0.prune50.5.05.tab > pangenome.snps.filtered.bi.274rice.md0.prune50.5.05.fasta

#Loaded fasta into SplitsTree4 v 4.16.2 (Huson and Bryant, 2006)  -> Fig 1B

#Fig S1B subset members for the green clonal lineage (n=86), similar process

vcftools --vcf /hpc-home/barragan/Italian_blast/pangenome.snps.filtered.bi.recode.vcf --keep 86_ind_green_rice.txt --max-missing 1.0 --recode --recode-INFO-all --out  pangenome.snps.filtered.bi.86green_rice.md0

vcf=pangenome.snps.filtered.bi.86green_rice.md0.recode.vcf
sed -i 's/Contig01/1/g' $vcf
sed -i 's/Contig02/2/g' $vcf
sed -i 's/Contig03/3/g' $vcf
sed -i 's/Contig04/4/g' $vcf
sed -i 's/Contig05/5/g' $vcf
sed -i 's/Contig06/6/g' $vcf
sed -i 's/Contig07/7/g' $vcf
sed -i 's/Contig08/8/g' $vcf
sed -i 's/Contig09/9/g' $vcf
sed -i 's/Contig10/10/g' $vcf
sed -i 's/Contig11/11/g' $vcf
sed -i 's/Contig12/12/g' $vcf
sed -i 's/Contig13/13/g' $vcf
sed -i 's/Contig14/14/g' $vcf
sed -i 's/Contig15/15/g' $vcf
sed -i 's/Contig16/16/g' $vcf
sed -i 's/Contig17/17/g' $vcf
sed -i 's/Contig18/18/g' $vcf
sed -i 's/Contig19/19/g' $vcf
sed -i 's/Contig20/20/g' $vcf
sed -i 's/Contig21/21/g' $vcf
sed -i 's/Contig22/22/g' $vcf
sed -i 's/Contig23/23/g' $vcf
sed -i 's/Contig24/24/g' $vcf

vcftools --vcf pangenome.snps.filtered.bi.86green_rice.md0.recode.vcf --plink --out pangenome.snps.filtered.bi.86green_rice.md0
plink --file pangenome.snps.filtered.bi.86green_rice.md0 --indep-pairwise 50 5 0.5 --out pangenome.snps.filtered.bi.86green_rice.md0.prune50.5.05
vcftools --vcf pangenome.snps.filtered.bi.86green_rice.md0.recode.vcf --positions pangenome.snps.filtered.bi.86green_rice.md0.prune50.5.05.prune.in --recode --recode-INFO-all --out pangenome.snps.filtered.bi.86green_rice.md0.prune50.5.05

bgzip -c pangenome.snps.filtered.bi.86green_rice.md0.prune50.5.05.recode.vcf > pangenome.snps.filtered.bi.86green_rice.md0.prune50.5.05.recode.vcf.gz
tabix -p vcf pangenome.snps.filtered.bi.86green_rice.md0.prune50.5.05.recode.vcf.gz
zcat  pangenome.snps.filtered.bi.86green_rice.md0.prune50.5.05.recode.vcf.gz | vcf-to-tab >  pangenome.snps.filtered.bi.86green_rice.md0.prune50.5.05.tab
perl vcf_tab_to_fasta_alignment.pl -i pangenome.snps.filtered.bi.86green_rice.md0.prune50.5.05.tab > pangenome.snps.filtered.bi.86green_rice.md0.prune50.5.05.fasta

# Created NJ tree (bs=100) using MEGA. 
##### vcf_tab_to_fasta_alignment.pl --> from https://github.com/JinfengChen/vcf-tab-to-fasta
```
