## Supplementary material for "Multiple horizontal mini-chromosome transfers drive genome evolution of clonal blast fungus lineages": Barragan_2024_mChrA_SupplementaryData: Italian_isolates_map.R

```
########################### Plot map of Italian rice blast fungus isolates ########################### 
############################ written by Cristina Barragan Feb 2024 ################################### 
getwd()
setwd("...")

library(tidyverse)
library(dplyr)
library(ggmap)
library(ggplot2)
##Define bounding box for the world, in this case Italy
italy <- c(left=5, bottom=37, right=20, top=48)

map <- get_stamenmap(italy, zoom=6, maptype = "terrain")

ggmap(map)
ustest4 <- read.delim(file="Italian_isolates_locations_Fig1.txt", sep= "\t", header=T)
ustest4

ggmap(map) + geom_point(data = ustest4, aes(x = ustest4$Lon, y = ustest4$Lat), color = "red", size = 2) +  geom_label(data = ustest4, aes(x = ustest4$Lon, y = ustest4$Lat, label = ustest4$Sample), vjust = 1.5, hjust = 0.5, size = 2, color = "black")
```
