## Supplementary material for "Multiple horizontal mini-chromosome transfers drive genome evolution of clonal blast fungus lineages": Barragan_2024_mChrA_SupplementaryData: mChrC_freq.R

```
#####################################################################################
############ Plot mChrC frequency written by Cristina Barragan Feb 2024##############
data <- read.table("mChrC_cov_BAM_normalized.txt", header=T)
data <- read.table("mChrC_freq_riceonly.txt", header=T)

data

library(ggplot2)
library(dplyr)
library(ggplot2)
library(dplyr)

# Bin the data as before
data <- data %>%
  mutate(bin = cut(normalized_mChrC_cov, breaks = seq(0, 100, by = 1), include.lowest = TRUE))
data

#oryza only
data <- data %>%
  mutate(bin = cut(mChrC_cov, breaks = seq(0, 100, by = 1), include.lowest = TRUE))
data

##for both
# Convert 'bin' back to a numeric midpoint for plotting
data$bin_midpoint <- as.numeric(gsub(",.*", "", gsub("\\(|\\[|\\]|\\)", "", data$bin))) + 0.5
data$bin_midpoint

# Plot with bin_midpoint as x -> all lineages
ggplot(data, aes(x = bin_midpoint, fill = HOST)) +
  geom_histogram(stat = "count", binwidth = 1, position = "stack", alpha = 1) +
  scale_x_continuous(breaks = seq(0, 100, by = 25), limits = c(0, 100)) +
  scale_fill_manual(values = c(
    "Triticum" = "#1D76B8",
    "Eleusine" = "#F27423",
    "Brachiaria1" = "#989898",
    "Oryza" = "#056A38",
    "Lolium" = "#9E2064",
    "Eragrostis" = "#CD7F29",
    "Brachiaria2" = "#998379",
    "Cenchrus" = "#3A53A4",
    "Zea" = "#BC2030",
    "Setaria" = "#6ABD45",
    "Digitaria" = "#231F20"
    # Add all the hosts here with their corresponding colors
  )) +
  labs(title = "Frequency of Normalized mChrA Coverage Values by Host",
       x = "Normalized mChrC Coverage Value", 
       y = "Count") +
  theme_minimal() +
  theme(axis.text.x = element_text(angle = 90, hjust = 1)) + geom_vline(aes(xintercept=61), color="red", linetype="dashed", size=1)


### Rice blast fungus lineage only

# Plot with bin_midpoint as x
ggplot(data, aes(x = data$bin_midpoint, fill = group)) +
  geom_histogram(stat = "count", position = "stack", alpha = 1) +
  scale_x_continuous(breaks = seq(0, 100, by = 25), limits = c(0, 100)) +
  scale_fill_manual(values = c(
    "I" = "#F69320",
    "IV" = "#BE1E2D",
    "II" = "#8CC540",
    "III" = "#1D76BB"

    # Add all the hosts here with their corresponding colors
  )) +
  labs(title = "Frequency of Normalized mChrC Coverage Values by Host",
       x = "Normalized mChrC Coverage Value", 
       y = "Count") +
  theme_minimal() +
  theme(axis.text.x = element_text(angle = 90, hjust = 1)) + geom_vline(aes(xintercept=61), color="red", linetype="dashed", size=1)

# Print out the plot
print(ggplot_object)

##########violinplot rice
library(ggplot2)
# Print out the plot
print(ggplot_object)

# Assuming 'data' is your dataframe and it has a column 'group'
group_counts <- data %>%
  group_by(group) %>%
  summarize(count = n())

# View the count of points in each group
print(group_counts)
####### add all samples
# Assuming 'data' is your original dataframe and it has columns 'bin_midpoint' and 'group'
# Create a new data frame for 'All Samples' group
data_all_samples <- data %>% 
  mutate(group = "All Samples")

# Combine with the original data
combined_data <- rbind(data, data_all_samples)

# Plotting
ggplot_object <- ggplot(combined_data, aes(x = group, y = bin_midpoint, fill = group)) +
  geom_violin(trim = FALSE) +
  geom_jitter(width = 0.2, size = 1, alpha = 0.5) +
  geom_boxplot(width = 0.1, outlier.shape = NA, alpha = 0.5) +
  scale_fill_manual(values = c(
    "All Samples" = "#333333",  # Color for the 'All Samples' group
    "I" = "#F69320",
    "IV" = "#BE1E2D",
    "II" = "#8CC540",
    "III" = "#1D76BB"
    # Add all the hosts here with their corresponding colors
  )) +
  labs(title = "Frequency of Normalized mChrA Coverage Values by Host",
       x = "Group",
       y = "Normalized mChrA Coverage Value") +
  theme_minimal() +
  theme(axis.text.x = element_text(angle = 90, hjust = 1)) +
  geom_hline(aes(yintercept = 61), color = "red", linetype = "dashed", size = 1) +
  ylim(0, 100)

# Print out the plot
print(ggplot_object)
```
