## Supplementary material for "Multiple horizontal mini-chromosome transfers drive genome evolution of clonal blast fungus lineages": Barragan_2024_mChrA_SupplementaryData: Horizontal_mini_coverage_AG006_final.py

```
####################################################################################
###########Script written by Angus Malmgren and Thorsten Langner Feb 2024
####################################################################################
library(reticulate)
py_install("matplotlib")
py_install("seaborn")
# Import required Python libraries
plt <- import("matplotlib.pyplot")
pd <- import("pandas")
np <- import("numpy")
sns <- import("seaborn")

#!/usr/bin/env python3
import matplotlib.pyplot as plt
import pandas as pd
import numpy as np
import seaborn as sns

print("Beginning plotting script:")

########### Specify contigs to use:
AG006_contigs=['AG006_Contig03','AG006_Contig04','AG006_Contig10','AG006_Contig11', 'AG006_Contig17']
###########


#Read in AG006 data:
AG006_coords_df=pd.read_csv("FG1846_04_pilon_polished_round2_nonmito_reordered_renamed.fasta.fai.plotting.txt.plotting.lessThanOrEqTo2mb.txt", delimiter=' ', names=['contig','start','end'])
AG006_coords_df=AG006_coords_df[AG006_coords_df.contig.isin(AG006_contigs)]

AG006_coverage_mini009_df=pd.read_csv("All_reads_trimmed_mini009_AG006_bwa_mem_ref_sorted_unique_q1_primary_mapped.bam_vs_Ref_cov_sliding_Wind.txt.sorted.txt", delimiter='\t', names=['contig','window_start','window_end','coverage'])
AG006_coverage_mini009_df=AG006_coverage_mini009_df[AG006_coverage_mini009_df.contig.isin(AG006_contigs)]
AG006_coverage_mini010_df=pd.read_csv("All_reads_trimmed_mini010_AG006_bwa_mem_ref_sorted_unique_q1_primary_mapped.bam_vs_Ref_cov_sliding_Wind.txt.sorted.txt", delimiter='\t', names=['contig','window_start','window_end','coverage'])
AG006_coverage_mini010_df=AG006_coverage_mini010_df[AG006_coverage_mini010_df.contig.isin(AG006_contigs)]
AG006_coverage_mini011_df=pd.read_csv("All_reads_trimmed_mini011_AG006_bwa_mem_ref_sorted_unique_q1_primary_mapped.bam_vs_Ref_cov_sliding_Wind.txt.sorted.txt", delimiter='\t', names=['contig','window_start','window_end','coverage'])
AG006_coverage_mini011_df=AG006_coverage_mini011_df[AG006_coverage_mini011_df.contig.isin(AG006_contigs)]
AG006_coverage_mini012_df=pd.read_csv("All_reads_trimmed_mini012_AG006_bwa_mem_ref_sorted_unique_q1_primary_mapped.bam_vs_Ref_cov_sliding_Wind.txt.sorted.txt", delimiter='\t', names=['contig','window_start','window_end','coverage'])
AG006_coverage_mini012_df=AG006_coverage_mini012_df[AG006_coverage_mini012_df.contig.isin(AG006_contigs)]
print(AG006_coverage_mini009_df)
print(AG006_coverage_mini010_df)
print(AG006_coverage_mini011_df)
print(AG006_coverage_mini012_df)
print()


#Plotting:
sns.set_style("white")

#Plot AG006:
x_vals_m09_c03=AG006_coverage_mini009_df[AG006_coverage_mini009_df["contig"]=="AG006_Contig03"]["window_end"]
y_vals_m09_c03=AG006_coverage_mini009_df[AG006_coverage_mini009_df["contig"]=="AG006_Contig03"]["coverage"]
x_vals_m09_c04=AG006_coverage_mini009_df[AG006_coverage_mini009_df["contig"]=="AG006_Contig04"]["window_end"]
y_vals_m09_c04=AG006_coverage_mini009_df[AG006_coverage_mini009_df["contig"]=="AG006_Contig04"]["coverage"]
x_vals_m09_c10=AG006_coverage_mini009_df[AG006_coverage_mini009_df["contig"]=="AG006_Contig10"]["window_end"]
y_vals_m09_c10=AG006_coverage_mini009_df[AG006_coverage_mini009_df["contig"]=="AG006_Contig10"]["coverage"]
x_vals_m09_c11=AG006_coverage_mini009_df[AG006_coverage_mini009_df["contig"]=="AG006_Contig11"]["window_end"]
y_vals_m09_c11=AG006_coverage_mini009_df[AG006_coverage_mini009_df["contig"]=="AG006_Contig11"]["coverage"]
x_vals_m09_c17=AG006_coverage_mini009_df[AG006_coverage_mini009_df["contig"]=="AG006_Contig17"]["window_end"]
y_vals_m09_c17=AG006_coverage_mini009_df[AG006_coverage_mini009_df["contig"]=="AG006_Contig17"]["coverage"]

x_vals_m10_c03=AG006_coverage_mini010_df[AG006_coverage_mini010_df["contig"]=="AG006_Contig03"]["window_end"]
y_vals_m10_c03=AG006_coverage_mini010_df[AG006_coverage_mini010_df["contig"]=="AG006_Contig03"]["coverage"]
x_vals_m10_c04=AG006_coverage_mini010_df[AG006_coverage_mini010_df["contig"]=="AG006_Contig04"]["window_end"]
y_vals_m10_c04=AG006_coverage_mini010_df[AG006_coverage_mini010_df["contig"]=="AG006_Contig04"]["coverage"]
x_vals_m10_c10=AG006_coverage_mini010_df[AG006_coverage_mini010_df["contig"]=="AG006_Contig10"]["window_end"]
y_vals_m10_c10=AG006_coverage_mini010_df[AG006_coverage_mini010_df["contig"]=="AG006_Contig10"]["coverage"]
x_vals_m10_c11=AG006_coverage_mini010_df[AG006_coverage_mini010_df["contig"]=="AG006_Contig11"]["window_end"]
y_vals_m10_c11=AG006_coverage_mini010_df[AG006_coverage_mini010_df["contig"]=="AG006_Contig11"]["coverage"]
x_vals_m10_c17=AG006_coverage_mini010_df[AG006_coverage_mini010_df["contig"]=="AG006_Contig17"]["window_end"]
y_vals_m10_c17=AG006_coverage_mini010_df[AG006_coverage_mini010_df["contig"]=="AG006_Contig17"]["coverage"]

x_vals_m11_c03=AG006_coverage_mini011_df[AG006_coverage_mini011_df["contig"]=="AG006_Contig03"]["window_end"]
y_vals_m11_c03=AG006_coverage_mini011_df[AG006_coverage_mini011_df["contig"]=="AG006_Contig03"]["coverage"]
x_vals_m11_c04=AG006_coverage_mini011_df[AG006_coverage_mini011_df["contig"]=="AG006_Contig04"]["window_end"]
y_vals_m11_c04=AG006_coverage_mini011_df[AG006_coverage_mini011_df["contig"]=="AG006_Contig04"]["coverage"]
x_vals_m11_c10=AG006_coverage_mini011_df[AG006_coverage_mini011_df["contig"]=="AG006_Contig10"]["window_end"]
y_vals_m11_c10=AG006_coverage_mini011_df[AG006_coverage_mini011_df["contig"]=="AG006_Contig10"]["coverage"]
x_vals_m11_c11=AG006_coverage_mini011_df[AG006_coverage_mini011_df["contig"]=="AG006_Contig11"]["window_end"]
y_vals_m11_c11=AG006_coverage_mini011_df[AG006_coverage_mini011_df["contig"]=="AG006_Contig11"]["coverage"]
x_vals_m11_c17=AG006_coverage_mini011_df[AG006_coverage_mini011_df["contig"]=="AG006_Contig17"]["window_end"]
y_vals_m11_c17=AG006_coverage_mini011_df[AG006_coverage_mini011_df["contig"]=="AG006_Contig17"]["coverage"]

x_vals_m12_c03=AG006_coverage_mini012_df[AG006_coverage_mini012_df["contig"]=="AG006_Contig03"]["window_end"]
y_vals_m12_c03=AG006_coverage_mini012_df[AG006_coverage_mini012_df["contig"]=="AG006_Contig03"]["coverage"]
x_vals_m12_c04=AG006_coverage_mini012_df[AG006_coverage_mini012_df["contig"]=="AG006_Contig04"]["window_end"]
y_vals_m12_c04=AG006_coverage_mini012_df[AG006_coverage_mini012_df["contig"]=="AG006_Contig04"]["coverage"]
x_vals_m12_c10=AG006_coverage_mini012_df[AG006_coverage_mini012_df["contig"]=="AG006_Contig10"]["window_end"]
y_vals_m12_c10=AG006_coverage_mini012_df[AG006_coverage_mini012_df["contig"]=="AG006_Contig10"]["coverage"]
x_vals_m12_c11=AG006_coverage_mini012_df[AG006_coverage_mini012_df["contig"]=="AG006_Contig11"]["window_end"]
y_vals_m12_c11=AG006_coverage_mini012_df[AG006_coverage_mini012_df["contig"]=="AG006_Contig11"]["coverage"]
x_vals_m12_c17=AG006_coverage_mini012_df[AG006_coverage_mini012_df["contig"]=="AG006_Contig17"]["window_end"]
y_vals_m12_c17=AG006_coverage_mini012_df[AG006_coverage_mini012_df["contig"]=="AG006_Contig17"]["coverage"]

#Count number of windows in each contig, to allow scaling of axes:
AG006_contig03_len=AG006_coverage_mini009_df[AG006_coverage_mini009_df.contig=='AG006_Contig03'].shape[0]
AG006_contig04_len=AG006_coverage_mini009_df[AG006_coverage_mini009_df.contig=='AG006_Contig04'].shape[0]
AG006_contig10_len=AG006_coverage_mini009_df[AG006_coverage_mini009_df.contig=='AG006_Contig10'].shape[0]
AG006_contig11_len=AG006_coverage_mini009_df[AG006_coverage_mini009_df.contig=='AG006_Contig11'].shape[0]
AG006_contig17_len=AG006_coverage_mini009_df[AG006_coverage_mini009_df.contig=='AG006_Contig17'].shape[0]


AG006_mini009_max=AG006_coverage_mini009_df[AG006_coverage_mini009_df['contig'].isin(['AG006_Contig17','AG006_Contig11', 'AG006_Contig10', 'AG006_Contig04', 'AG006_Contig03'])]['coverage'].max()
print('max:', AG006_mini009_max)
AG006_mini010_max=AG006_coverage_mini010_df[AG006_coverage_mini009_df['contig'].isin(['AG006_Contig17','AG006_Contig11', 'AG006_Contig10', 'AG006_Contig04', 'AG006_Contig03'])]['coverage'].max()
print('max:', AG006_mini010_max)
AG006_mini011_max=AG006_coverage_mini011_df[AG006_coverage_mini009_df['contig'].isin(['AG006_Contig17','AG006_Contig11', 'AG006_Contig10', 'AG006_Contig04', 'AG006_Contig03'])]['coverage'].max()
print('max:', AG006_mini011_max)
AG006_mini012_max=AG006_coverage_mini012_df[AG006_coverage_mini009_df['contig'].isin(['AG006_Contig17','AG006_Contig11', 'AG006_Contig10', 'AG006_Contig04', 'AG006_Contig03'])]['coverage'].max()
print('max:', AG006_mini012_max)

#Get nearest 100:
AG006_mini009_max_nearest100=(int(AG006_mini009_max/100.0)+1)*100.0
AG006_mini010_max_nearest100=(int(AG006_mini010_max/100.0)+1)*100.0
AG006_mini011_max_nearest100=(int(AG006_mini011_max/100.0)+1)*100.0
AG006_mini012_max_nearest100=(int(AG006_mini012_max/100.0)+1)*100.0

#Use the following to get common y-axis: fig,axs = plt.subplots(4,4, sharex='col', sharey=True,
fig,axs = plt.subplots(5,5, sharex='col', sharey='row', gridspec_kw={'hspace':0.1, 'wspace':0.05, 'width_ratios':[AG006_contig03_len,AG006_contig04_len,AG006_contig10_len,AG006_contig11_len,AG006_contig17_len]}, figsize=(40,8))
fig.suptitle('AG006', fontsize=30)
axs[0,0].plot(x_vals_m09_c03, y_vals_m09_c03,color='black')
axs[0,0].set_ylim([0,AG006_mini009_max_nearest100])
axs[0,1].plot(x_vals_m09_c04, y_vals_m09_c04,color='black')
axs[0,2].plot(x_vals_m09_c10, y_vals_m09_c10,color='black')
axs[0,3].plot(x_vals_m09_c11, y_vals_m09_c11,color='black')
axs[0,4].plot(x_vals_m09_c17, y_vals_m09_c17,color='black')


axs[0,0].fill_between(x_vals_m09_c03, 0, y_vals_m09_c03, color='red')
axs[0,1].fill_between(x_vals_m09_c04, 0, y_vals_m09_c04, color='red')
axs[0,2].fill_between(x_vals_m09_c10, 0, y_vals_m09_c10, color='red')
axs[0,3].fill_between(x_vals_m09_c11, 0, y_vals_m09_c11, color='red')
axs[0,4].fill_between(x_vals_m09_c17, 0, y_vals_m09_c17, color='red')


axs[0,0].set_title('Contig03', fontsize=20)
axs[0,1].set_title('Contig04', fontsize=20)
axs[0,2].set_title('Contig10', fontsize=20)
axs[0,3].set_title('Contig11', fontsize=20)
axs[0,4].set_title('Contig17', fontsize=20)


axs[1,0].plot(x_vals_m10_c03, y_vals_m10_c03,color='black')
axs[1,0].set_ylim([0,AG006_mini010_max_nearest100])
axs[1,1].plot(x_vals_m10_c04, y_vals_m10_c04,color='black')
axs[1,2].plot(x_vals_m10_c10, y_vals_m10_c10,color='black')
axs[1,3].plot(x_vals_m10_c11, y_vals_m10_c11,color='black')#, 'tab:red')
axs[1,4].plot(x_vals_m10_c17, y_vals_m10_c17,color='black')#, 'tab:red')


axs[1,0].fill_between(x_vals_m10_c03, 0, y_vals_m10_c03, color='red')
axs[1,1].fill_between(x_vals_m10_c04, 0, y_vals_m10_c04, color='red')
axs[1,2].fill_between(x_vals_m10_c10, 0, y_vals_m10_c10, color='red')
axs[1,3].fill_between(x_vals_m10_c11, 0, y_vals_m10_c11, color='red')
axs[1,4].fill_between(x_vals_m10_c17, 0, y_vals_m10_c17, color='red')


axs[2,0].plot(x_vals_m11_c03, y_vals_m11_c03,color='black')
axs[2,0].set_ylim([0,AG006_mini011_max_nearest100])
axs[2,1].plot(x_vals_m11_c04, y_vals_m11_c04,color='black')
axs[2,2].plot(x_vals_m11_c10, y_vals_m11_c10,color='black')
axs[2,3].plot(x_vals_m11_c11, y_vals_m11_c11,color='black')
axs[2,4].plot(x_vals_m11_c17, y_vals_m11_c17,color='black')


axs[2,0].fill_between(x_vals_m11_c03, 0, y_vals_m11_c03, color='red')
axs[2,1].fill_between(x_vals_m11_c04, 0, y_vals_m11_c04, color='red')
axs[2,2].fill_between(x_vals_m11_c10, 0, y_vals_m11_c10, color='red')
axs[2,3].fill_between(x_vals_m11_c11, 0, y_vals_m11_c11, color='red')
axs[2,4].fill_between(x_vals_m11_c17, 0, y_vals_m11_c17, color='red')


axs[3,0].plot(x_vals_m12_c03, y_vals_m12_c03,color='black')
axs[3,0].set_ylim([0,AG006_mini012_max_nearest100])
axs[3,1].plot(x_vals_m12_c04, y_vals_m12_c04,color='black')
axs[3,2].plot(x_vals_m12_c10, y_vals_m12_c10,color='black')
axs[3,3].plot(x_vals_m12_c11, y_vals_m12_c11,color='black')
axs[3,4].plot(x_vals_m12_c17, y_vals_m12_c17,color='black')


axs[3,0].xaxis.major.formatter._useMathText = True
axs[3,1].xaxis.major.formatter._useMathText = True
axs[3,2].xaxis.major.formatter._useMathText = True
axs[3,3].xaxis.major.formatter._useMathText = True
axs[3,4].xaxis.major.formatter._useMathText = True


axs[3,0].fill_between(x_vals_m12_c03, 0, y_vals_m12_c03, color='red')
axs[3,1].fill_between(x_vals_m12_c04, 0, y_vals_m12_c04, color='red')
axs[3,2].fill_between(x_vals_m12_c10, 0, y_vals_m12_c10, color='red')
axs[3,3].fill_between(x_vals_m12_c11, 0, y_vals_m12_c11, color='red')
axs[3,4].fill_between(x_vals_m12_c11, 0, y_vals_m12_c17, color='red')


#To remove axis labels from 'inner' plots:
for ax in axs.flat:
	ax.label_outer()


plt.savefig('Horizontal_mini_coverage_AG006_new.pdf')#.png
```
