## Supplementary material for "Multiple horizontal mini-chromosome transfers drive genome evolution of clonal blast fungus lineages": Barragan_2024_mChrA_SupplementaryData: ciclize_AG006_AG002.R

```
## Initial script outline by Thorsten Langner and Angus Malmgren and further developed by Cristina Barragan Feb 2024
#######################################################################################################################

library(tidyr)
library(circlize)
library(dplyr)
library(tidyverse)
getwd()
#################################### based on TLs script 
circos.par("start.degree" = 90)
######AG006 and AG002
chr <- c("AG006_Contig01", "AG006_Contig02", "AG006_Contig03", "AG006_Contig04", "AG006_Contig05", "AG006_Contig06", "AG006_Contig07", "AG006_Contig08", "AG006_Contig09", "AG006_Contig10","AG006_Contig11", "AG006_Contig12", "AG006_Contig13", "AG006_Contig14", "AG006_Contig15", "AG006_Contig16", "AG006_Contig17", "AG006_Contig18","AG006_Contig19", "AG006_Contig20", "AG006_Contig21", "AG006_Contig22", "AG006_Contig23", "AG006_Contig24","AG002_Contig01","AG002_Contig02","AG002_Contig03", "AG002_Contig04", "AG002_Contig05", "AG002_Contig06", "AG002_Contig07", "AG002_Contig08", "AG002_Contig09", "AG002_Contig10","AG002_Contig11", "AG002_Contig12", "AG002_Contig13", "AG002_Contig14", "AG002_Contig15", "AG002_Contig16", "AG002_Contig17", "AG002_Contig18","AG002_Contig19", "AG002_Contig20", "AG002_Contig21", "AG002_Contig22", "AG002_Contig23", "AG002_Contig24", "AG002_Contig25", "AG002_Contig26", "AG002_Contig27", "AG002_Contig28", "AG002_Contig29", "AG002_Contig30", "AG002_Contig31","AG002_Contig32","AG002_Contig33", "AG002_Contig34", "AG002_Contig35","AG002_Contig36")
start <- c(1,1,1,1,1,1,1,1,1,1,1,1,1,1,1,1,1,1,1,1,1,1,1,1,1,1,1,1,1,1,1,1,1,1,1,1,1,1,1,1,1,1,1,1,1,1,1,1,1,1,1,1,1,1,1,1,1,1,1,1)
end <- c(11008917,8645043,1697966,796569,60573,6618557,51574,5731635,4595668,1201610,170974,1464303,4077659,55256,72685,77902,104405,133460,96136,73650,73030,52001,70829,75409,3985893,715898,3545443,2385863,4245495,1477886,154114,1024435,5624068,5692689,4555161,138019,6389779,4598193,72054,68216,88316,76010,96438,71349,78730,93972,109517,82058,20785,1896,7104,35812,87389,111446,74273,138865,78102,70893,66394,23914)
###########
df <- data.frame(chr, start, end)
df
dim(df)
circos.initializeWithIdeogram(df)

#adding rainbow color to chromosomes
circos.track(ylim = c(0, 1), bg.col = c(rainbow(50, alpha = 1)), bg.border = NA, track.height = 0.1)
AG006_coords_df <- read_tsv("AG006_AG002_algn_coords_long.bed", col_names = T)
AG006_coords_df
dim(AG006_coords_df)

Br62_coords_df <- read_tsv("AG002_AG006_algn_coords_long.bed", col_names = T)
Br62_coords_df
dim(Br62_coords_df)

#adding genomic links between two sets of bed files
#circos.genomicLink(AG006_coords_df, Br62_coords_df, col = rand_color(nrow(AG006_coords_df), transparency = 0.5), border = NA)

circos.genomicLink(AG006_coords_df, Br62_coords_df, col = "dimgrey", border = NA)
```
