## Supplementary material for "Multiple horizontal mini-chromosome transfers drive genome evolution of clonal blast fungus lineages": Barragan_2024_mChrA_SupplementaryData: AG006_alignment.R

```
## Initial script outline by Thorsten Langner and Angus Malmgren and further developed by Cristina Barragan Feb 2024
#######################################################################################################################

if (!requireNamespace("BiocManager", quietly = TRUE))
  install.packages("BiocManager")
BiocManager::install("karyoploteR")

#https://bernatgel.github.io/karyoploter_tutorial//Tutorial/PlotTypes/PlotTypes.html

library(karyoploteR)
library(tidyverse)
library(ggplot2)

#AG00G_contig10 and AG006_contig17

custom.genome <- toGRanges(data.frame(chr=c("AG006_Contig10","AG006_Contig17"), start=c(1,1), end=c(1201610,104405)))
kp <- plotKaryotype(genome = custom.genome, plot.type = 2)

kpAddBaseNumbers(kp, tick.dist = 100000, tick.len = 10, tick.col="red", cex=0.7, minor.tick.dist = 10000, minor.tick.len = 5, minor.tick.col = "gray")

AG006_coords_df <- read_tsv("AG006_Contig10_17.txt", col_names = T)
AG006_coords_df

Br62_coords_df <- read_tsv("AG006_Contig17_10.txt", col_names = T)
Br62_coords_df 

AG006_coords_ranges <- toGRanges(data.frame(chr=c(AG006_coords_df$contig), start=c(AG006_coords_df$start), end=c(AG006_coords_df$end), strand=c(AG006_coords_df$strand)))
AG006_coords_ranges

Br62_coords_ranges <- toGRanges(data.frame(chr=c(Br62_coords_df$contig), start=c(Br62_coords_df$start), end=c(Br62_coords_df$end), strand=c(Br62_coords_df$strand)))
Br62_coords_ranges

#plot
karyoploteR::kpPlotLinks(kp, data = AG006_coords_ranges, data2 = Br62_coords_ranges, y= -0.1, col="dimgray", cex=0.2)


#create a dataframe
chr <- c("contig10","contig07")
start <- c(1, 1)
end <- c(1201610, 1185594)
df <- data.frame(chr, start, end)
df

#AG00G_contig10 and Br62_contig07
custom.genome <- toGRanges(data.frame(chr=c("contig10","contig07"), start=c(1,1), end=c(1201610,1185594)))
kp <- plotKaryotype(genome = custom.genome)
kpAddBaseNumbers(kp, tick.dist = 100000, tick.len = 10, tick.col="red", cex=0.7, minor.tick.dist = 10000, minor.tick.len = 5, minor.tick.col = "gray")


for (i in 1:nrow(df)) {
  lines(df$start[i], df$end[i], df$start[i+1], df$end[i+1], col="blue")
}


start.regs <- toGRanges(data.frame("contig10", 1, 300))
end.regs <- toGRanges(data.frame("contig07", 200, 900))

kpPlotLinks(kp, data=start.regs, data2=end.regs)

start.regs <- toGRanges(data.frame(chr=c("contig10"), 100, 800))
end.regs <- toGRanges(data.frame(chr=c("contig07"), 200, 900))
kp <- plotKaryotype(genome = custom.genome) 
kpPlotLinks(kp, data=start.regs, data2=end.regs)


custom.cytobands <- toGRanges("contig10_cytobands.txt")
custom.cytobands
custom.cytobands <- toGRanges("allcontigs_MoTeR.txt")


#AG006
custom.genome <- toGRanges(data.frame(chr=c("contig01", "contig02", "contig03", "contig04", "contig05", "contig06", "contig07", "contig08", "contig09", "contig10","contig11", "contig12", "contig13", "contig14", "contig15", "contig16", "contig17", "contig18","contig19", "contig20", "contig21", "contig22", "contig23", "contig24"), start=c(1,1,1,1,1,1,1,1,1,1,1,1,1,1,1,1,1,1,1,1,1,1,1,1), end=c(11008917,8645043,1697966,796569,60573,6618557,51574,5731635,4595668,1201610,170974,1464303,4077659,55256,72685,77902,104405,133460,96136,73650,73030,52001,70829,75409)))
kp <- plotKaryotype(genome = custom.genome)
custom.cytobands <- toGRanges("contig10_cytobands.txt")
custom.cytobands
custom.cytobands <- toGRanges("allcontigs_MoTeR.txt")

#LpKY97

custom.genome <- toGRanges(data.frame(chr=c("1", "2", "3", "4", "5", "6", "7", "min1", "min2"), start=c(1,1,1,1,1,1,1,1,1), end=c(6359358,7985541,7463082,5442467,4430380,6155580,3866546,2996683,913334)))
kp <- plotKaryotype(genome = custom.genome)

custom.cytobands <- toGRanges("allcontigs_MoTeR_LpKY97.txt")

#B71

custom.genome <- toGRanges(data.frame(chr=c("1", "2", "3", "4", "5", "6", "7", "mini"), start=c(1,1,1,1,1,1,1,1), end=c(6668242,7915472,8237918,5413369,4460276,6133529,4079358,1903245)))
kp <- plotKaryotype(genome = custom.genome)

custom.cytobands <- toGRanges("allcontigs_MoTeR_B71.txt")
custom.cytobands <- toGRanges("allcontigs_allrepeats_B71.txt")

#FR13

custom.genome <- toGRanges(data.frame(chr=c("contig01", "contig02", "contig03", "contig04", "contig05", "contig06", "contig07", "contig08", "contig09", "contig10","contig11", "contig12", "contig13", "contig14", "contig15", "contig16", "contig17", "contig18","contig19", "contig20", "contig21", "contig22", "contig23", "contig24", "contig25", "contig26", "contig27", "contig28", "contig29", "contig30", "contig31"), start=c(1,1,1,1,1,1,1,1,1,1,1,1,1,1,1,1,1,1,1,1,1,1,1,1,1,1,1,1,1,1,1), end=c(7382384,6680418,6077162,5398440,4622961,4094040,2617860,2137937,1647583,1626924,1222279,840477,338330,334587,273887,224111,223742,94828,86637,60288,59180,48370,48775,48310,45575,39710,32743,30392,31084,21987,19414)))
kp <- plotKaryotype(genome = custom.genome)

custom.cytobands <- toGRanges("allcontigs_MoTeR_FR13.txt")

#AG002

custom.genome <- toGRanges(data.frame(chr=c("contig01", "contig02", "contig03", "contig04", "contig05", "contig06", "contig07", "contig08", "contig09", "contig10","contig11", "contig12", "contig13", "contig14", "contig15", "contig16", "contig17", "contig18","contig19", "contig20", "contig21", "contig22", "contig23", "contig24", "contig25", "contig26", "contig27", "contig28", "contig29", "contig30", "contig31","contig32","contig33", "contig34", "contig35","contig36"), start=c(1,1,1,1,1,1,1,1,1,1,1,1,1,1,1,1,1,1,1,1,1,1,1,1,1,1,1,1,1,1,1,1,1,1,1,1), end=c(3985893,715898,3545443,2385863,4245495,1477886,154114,1024435,5624068,5692689,4555161,138019,6389779,4598193,72054,68216,88316,76010,96438,71349,78730,93972,109517,82058,20785,1896,7104,35812,87389,111446,74273,138865,78102,70893,66394,23914)))
kp <- plotKaryotype(genome = custom.genome)

custom.cytobands <- toGRanges("allcontigs_MoTeR_AG002.txt")

#70-15

custom.genome <- toGRanges(data.frame(chr=c("1", "2", "3", "4", "5", "6", "7", "supercont8.8 "), start=c(1,1,1,1,1,1,1,1), end=c(7978604,8319966,6606598,5546968,4490059,4133993,3415785,535760)))
kp <- plotKaryotype(genome = custom.genome)

custom.cytobands <- toGRanges("allcontigs_MoTeR_70-15.txt")

kp <- plotKaryotype(genome = custom.genome, cytobands = custom.cytobands)

#MZ1-6

custom.genome <- toGRanges(data.frame(chr=c("1", "2", "3", "4", "5", "6", "7"), start=c(1,1,1,1,1,1,1), end=c(6129159,8796158,6918261,5706987,4408553,6060913,4683251)))
kp <- plotKaryotype(genome = custom.genome)

custom.cytobands <- toGRanges("allcontigs_MoTeR_MZ1-6.txt")

#AG032
custom.genome <- toGRanges(data.frame(chr=c("contig01", "contig02", "contig03", "contig04", "contig05", "contig06", "contig07", "contig08", "contig09", "contig10","contig11", "contig12", "contig13", "contig14", "contig15", "contig16", "contig17", "contig18","contig19", "contig20", "contig21", "contig22", "contig23", "contig24"), start=c(1,1,1,1,1,1,1,1,1,1,1,1,1,1,1,1,1,1,1,1,1,1,1,1), end=c(87960,5961411,6503571,8843348,47277,6528442,5215895,501660,4620578,504069,916242,158012,1236074,4085146,91926,54323,49253,93912,1925,60970,94420,54403,81516,124577)))
kp <- plotKaryotype(genome = custom.genome)
custom.cytobands <- toGRanges("allcontigs_MoTeR_AG032.txt")

#AG038
custom.genome <- toGRanges(data.frame(chr=c("contig01", "contig02", "contig03", "contig04", "contig05", "contig06", "contig07", "contig08", "contig09", "contig10","contig11", "contig12", "contig13", "contig14", "contig15", "contig16", "contig17", "contig18","contig19"), start=c(1,1,1,1,1,1,1,1,1,1,1,1,1,1,1,1,1,1,1), end=c(7293716,9163141,1627133,5360410,5673907,662615,4867123,6481103,4120698,166626,160299,99710,77874,94636,46127,59796,171549,91142,73564)))
kp <- plotKaryotype(genome = custom.genome)
custom.cytobands <- toGRanges("allcontigs_MoTeR_AG038.txt")

#AG039
custom.genome <- toGRanges(data.frame(chr=c("contig01", "contig02", "contig03", "contig04", "contig05", "contig06", "contig07", "contig08", "contig09", "contig10","contig11", "contig12", "contig13", "contig14", "contig15", "contig16", "contig17", "contig18","contig19","contig20", "contig21", "contig22", "contig23", "contig24","contig25","contig26"), start=c(1,1,1,1,1,1,1,1,1,1,1,1,1,1,1,1,1,1,1,1,1,1,1,1,1,1), end=c(6137354,2406083,2075711,11002933,6249570,677523,5699073,6608653,50769,4154976,1344442,201837,92281,81761,117684,9276,71398,1852,64883,56613,53025,57863,61786,83293,59684,75635)))
kp <- plotKaryotype(genome = custom.genome)
custom.cytobands <- toGRanges("allcontigs_MoTeR_AG039.txt")

#AG059
custom.genome <- toGRanges(data.frame(chr=c("contig01", "contig02", "contig03", "contig04", "contig05", "contig06", "contig07", "contig08", "contig09", "contig10","contig11", "contig12", "contig13", "contig14", "contig15", "contig16", "contig17", "contig18","contig19", "contig20", "contig21", "contig22", "contig23", "contig24", "contig25", "contig26", "contig27", "contig28", "contig29", "contig30", "contig31","contig32","contig33", "contig34", "contig35","contig36", "contig37"), start=c(1,1,1,1,1,1,1,1,1,1,1,1,1,1,1,1,1,1,1,1,1,1,1,1,1,1,1,1,1,1,1,1,1,1,1,1,1), end=c(153748,6004778,897278,7152446,1707998,134113,6740178,1959904,2988346,928142,5900770,356302,370800,180500,753198,4313370,4940171,177180,59092,54239,46598,138085,66015,81746,246483,95971,60618,49338,93215,480560,51209,78430,91891,57607,173827,116606,42369)))
kp <- plotKaryotype(genome = custom.genome)

custom.cytobands <- toGRanges("allcontigs_MoTeR_AG059.txt")

#AG098
custom.genome <- toGRanges(data.frame(chr=c("contig01", "contig02", "contig03", "contig04", "contig05", "contig06", "contig07", "contig08", "contig09", "contig10","contig11", "contig12", "contig13", "contig14", "contig15", "contig16", "contig17", "contig18","contig19", "contig20", "contig21", "contig22", "contig23", "contig24", "contig25", "contig26", "contig27", "contig28", "contig29", "contig30", "contig31","contig32","contig33"), start=c(1,1,1,1,1,1,1,1,1,1,1,1,1,1,1,1,1,1,1,1,1,1,1,1,1,1,1,1,1,1,1,1,1), end=c(6094221,7877299,1104432,548346,1925088,652874,6607814,5678089,200313,385518,4600349,6522306,4085952,192324,97956,54194,66530,98480,500,118216,1902,62607,73675,88201,61877,78194,55802,103578,34020,87549,151152,46122,55346)))
kp <- plotKaryotype(genome = custom.genome)

custom.cytobands <- toGRanges("allcontigs_MoTeR_AG098.txt")

#PR003
custom.genome <- toGRanges(data.frame(chr=c("contig01", "contig02", "contig03", "contig04", "contig05", "contig06", "contig07", "contig08", "contig09", "contig10","contig11", "contig12", "contig13", "contig14", "contig15", "contig16"), start=c(1,1,1,1,1,1,1,1,1,1,1,1,1,1,1,1), end=c(6063740,8582813,1676349,6557505,5682125,78272,5081269,6310446,4048513,131938,93305,104117,42085,41659,59771,61291)))
kp <- plotKaryotype(genome = custom.genome)
custom.cytobands <- toGRanges("allcontigs_MoTeR_PR003.txt")

#San_Andrea
custom.genome <- toGRanges(data.frame(chr=c("contig01", "contig02", "contig03", "contig04", "contig05", "contig06", "contig07", "contig08", "contig09", "contig10","contig11", "contig12", "contig13", "contig14", "contig15", "contig16", "contig17", "contig18","contig19", "contig20", "contig21", "contig22", "contig23", "contig24", "contig25", "contig26", "contig27", "contig28", "contig29", "contig30", "contig31","contig32","contig33", "contig34", "contig35"), start=c(1,1,1,1,1,1,1,1,1,1,1,1,1,1,1,1,1,1,1,1,1,1,1,1,1,1,1,1,1,1,1,1,1,1,1), end=c(1367262,2334244,242184,3024089,6303665,4585811,1639350,475668,1000554,871259,5937455,1964240,120390,3736388,5588979,4122582,3268206,85166,66937,194211,113348,198773,265059,139168,110708,111798,53730,101770,88543,58785,88885,53723,56704,74452,65628)))
kp <- plotKaryotype(genome = custom.genome)
custom.cytobands <- toGRanges("allcontigs_MoTeR_San_Andrea.txt")

kp <- plotKaryotype(genome = custom.genome, cytobands = custom.cytobands)

kpAddBaseNumbers(kp,tick.dist = 100000, tick.len = 10, tick.col="red", cex=0.5,
                 minor.tick.dist = 100000, minor.tick.len = 5, minor.tick.col = "gray")

kpAddCytobandLabels(kp, cex=0.3, force.all = TRUE, srt=90, col="gray")

#AG006 effectors 
custom.genome <- toGRanges(data.frame(chr=c("contig01", "contig02", "contig03", "contig04", "contig05", "contig06", "contig07", "contig08", "contig09", "contig10","contig11", "contig12", "contig13", "contig14", "contig15", "contig16", "contig17", "contig18","contig19", "contig20", "contig21", "contig22", "contig23", "contig24"), start=c(1,1,1,1,1,1,1,1,1,1,1,1,1,1,1,1,1,1,1,1,1,1,1,1), end=c(11008917,8645043,1697966,796569,60573,6618557,51574,5731635,4595668,1201610,170974,1464303,4077659,55256,72685,77902,104405,133460,96136,73650,73030,52001,70829,75409)))
kp <- plotKaryotype(genome = custom.genome)
kpAddBaseNumbers(kp, tick.dist = 1000000, tick.len = 10, tick.col="red", cex=0.5, minor.tick.dist = 100000, minor.tick.len = 5, minor.tick.col = "gray")

custom.cytobands <- toGRanges("AG006_178effectors_original.txt")
custom.cytobands <- toGRanges("AG006_WG_987effectors.txt")
custom.cytobands
kp <- plotKaryotype(genome = custom.genome, cytobands = custom.cytobands)
kp

#Br62 effectors
custom.genome <- toGRanges(data.frame(chr=c("contig01", "contig02", "contig03", "contig04", "contig06", "contig07", "contig08", "contig09", "contig10","contig11"), start=c(1,1,1,1,1,1,1,1,1,1), end=c(5686109,5098061,7337754,8318372,6027403,1185594,3851847,6285418,69893,84649)))
kp <- plotKaryotype(genome = custom.genome)
custom.cytobands <- toGRanges("Br62_178effectors_original.txt")
custom.cytobands <- toGRanges("Br62_WG_937effectors.txt")
custom.cytobands
kp <- plotKaryotype(genome = custom.genome, cytobands = custom.cytobands)
kp

#AG006_contig10
custom.genome <- toGRanges(data.frame(chr=c("contig10"), start=c(1), end=c(1201610)))
kp <- plotKaryotype(genome = custom.genome)
kpAddBaseNumbers(kp, tick.dist = 100000, tick.len = 10, tick.col="red", cex=0.7, minor.tick.dist = 10000, minor.tick.len = 5, minor.tick.col = "gray")

custom.cytobands <- toGRanges("AG006_178effectors_original_contig10.txt")
custom.cytobands
kp <- plotKaryotype(genome = custom.genome, cytobands = custom.cytobands)
kp


#Br62_contig07
custom.genome <- toGRanges(data.frame(chr=c("contig07"), start=c(1), end=c(1185594)))
kp <- plotKaryotype(genome = custom.genome)
kpAddBaseNumbers(kp, tick.dist = 100000, tick.len = 10, tick.col="red", cex=0.7, minor.tick.dist = 10000, minor.tick.len = 5, minor.tick.col = "gray")

custom.cytobands <- toGRanges("Br62_178effectors_original_contig07.txt")
custom.cytobands
kp <- plotKaryotype(genome = custom.genome, cytobands = custom.cytobands)
kp
```
