## Supplementary material for "Multiple horizontal mini-chromosome transfers drive genome evolution of clonal blast fungus lineages": Barragan_2024_mChrA_SupplementaryData: mChrC_alignment.R

```
## Initial script outline by Thorsten Langner and Angus Malmgren and further developed by Cristina Barragan Feb 2024
#######################################################################################################################
####Pairwise alignments

if (!requireNamespace("BiocManager", quietly = TRUE))
  install.packages("BiocManager")
BiocManager::install("karyoploteR")

#https://bernatgel.github.io/karyoploter_tutorial//Tutorial/PlotTypes/PlotTypes.html

library(karyoploteR)
library(tidyverse)
library(ggplot2)

#conserved_mChr
#FR13 and AG039
custom.genome <- toGRanges(data.frame(chr=c("FR13_Cont09", "AG039_Contig08"), start=c(1,1), end=c(1647583,1544363)))
kp <- plotKaryotype(genome = custom.genome, plot.type = 2)
kpAddBaseNumbers(kp, tick.dist = 100000, tick.len = 10, tick.col="red", cex=0.7, minor.tick.dist = 10000, minor.tick.len = 5, minor.tick.col = "gray")
AG006_coords_df <- read_tsv("AG039_Contig08_FR13.txt", col_names = T)
AG006_coords_df
Br62_coords_df <- read_tsv("FR13_AG039_Contig08.txt", col_names = T)
Br62_coords_df
AG006_coords_ranges <- toGRanges(data.frame(chr=c(AG006_coords_df$contig), start=c(AG006_coords_df$start), end=c(AG006_coords_df$end), strand=c(AG006_coords_df$strand)))
AG006_coords_ranges
Br62_coords_ranges <- toGRanges(data.frame(chr=c(Br62_coords_df$contig), start=c(Br62_coords_df$start), end=c(Br62_coords_df$end), strand=c(Br62_coords_df$strand)))
Br62_coords_ranges
karyoploteR::kpPlotLinks(kp, data = AG006_coords_ranges, data2 = Br62_coords_ranges, y= -0.1, col="dimgray", cex=0.2)

#AG039_PR003
custom.genome <- toGRanges(data.frame(chr=c("AG039_Contig08","PR003_Contig03"), start=c(1,1), end=c(1544363,1676349)))
kp <- plotKaryotype(genome = custom.genome, plot.type = 2)
kpAddBaseNumbers(kp, tick.dist = 100000, tick.len = 10, tick.col="red", cex=0.7, minor.tick.dist = 10000, minor.tick.len = 5, minor.tick.col = "gray")
AG006_coords_df <- read_tsv("AG039_Contig08_mchr_PR003.txt", col_names = T)
AG006_coords_df
Br62_coords_df <- read_tsv("PR003_AG039_Contig08_mchr.txt", col_names = T)
Br62_coords_df
AG006_coords_ranges <- toGRanges(data.frame(chr=c(AG006_coords_df$contig), start=c(AG006_coords_df$start), end=c(AG006_coords_df$end), strand=c(AG006_coords_df$strand)))
AG006_coords_ranges
Br62_coords_ranges <- toGRanges(data.frame(chr=c(Br62_coords_df$contig), start=c(Br62_coords_df$start), end=c(Br62_coords_df$end), strand=c(Br62_coords_df$strand)))
Br62_coords_ranges
karyoploteR::kpPlotLinks(kp, data = AG006_coords_ranges, data2 = Br62_coords_ranges, y= -0.1, col="dimgray", cex=0.2)

#PR003_AG098
custom.genome <- toGRanges(data.frame(chr=c("PR003_Contig03", "AG098_Contig03"), start=c(1,1), end=c(1676349, 1676349)))
kp <- plotKaryotype(genome = custom.genome, plot.type = 2)
kpAddBaseNumbers(kp, tick.dist = 100000, tick.len = 10, tick.col="red", cex=0.7, minor.tick.dist = 10000, minor.tick.len = 5, minor.tick.col = "gray")
AG006_coords_df <- read_tsv("PR003_AG098_concat.txt", col_names = T)
AG006_coords_df 
Br62_coords_df <- read_tsv("AG098_concat_PR003.txt", col_names = T)
Br62_coords_df
AG006_coords_ranges <- toGRanges(data.frame(chr=c(AG006_coords_df$contig), start=c(AG006_coords_df$start), end=c(AG006_coords_df$end), strand=c(AG006_coords_df$strand)))
AG006_coords_ranges
Br62_coords_ranges <- toGRanges(data.frame(chr=c(Br62_coords_df$contig), start=c(Br62_coords_df$start), end=c(Br62_coords_df$end), strand=c(Br62_coords_df$strand)))
Br62_coords_ranges
karyoploteR::kpPlotLinks(kp, data = AG006_coords_ranges, data2 = Br62_coords_ranges, y= -0.1, col="dimgray", cex=0.2)
#AG098_AG038 -> re-check
custom.genome <- toGRanges(data.frame(chr=c("AG098_Contig03", "AG038_Contig03"), start=c(1,1), end=c(1652778, 1627133)))
kp <- plotKaryotype(genome = custom.genome, plot.type = 2)
kpAddBaseNumbers(kp, tick.dist = 100000, tick.len = 10, tick.col="red", cex=0.7, minor.tick.dist = 10000, minor.tick.len = 5, minor.tick.col = "gray")
AG006_coords_df <- read_tsv("AG098_AG038_10k.txt", col_names = T)
AG006_coords_df
Br62_coords_df <- read_tsv("AG038_AG098_10k.txt", col_names = T)
Br62_coords_df
AG006_coords_ranges <- toGRanges(data.frame(chr=c(AG006_coords_df$contig), start=c(AG006_coords_df$start), end=c(AG006_coords_df$end), strand=c(AG006_coords_df$strand)))
AG006_coords_ranges
Br62_coords_ranges <- toGRanges(data.frame(chr=c(Br62_coords_df$contig), start=c(Br62_coords_df$start), end=c(Br62_coords_df$end), strand=c(Br62_coords_df$strand)))
Br62_coords_ranges
karyoploteR::kpPlotLinks(kp, data = AG006_coords_ranges, data2 = Br62_coords_ranges, y= -0.1, col="dimgray", cex=0.2)
#AG038_AG002
custom.genome <- toGRanges(data.frame(chr=c("AG038_Contig03", "AG002_Contig06-7"), start=c(1,1), end=c(1627133, 1632000)))
kp <- plotKaryotype(genome = custom.genome, plot.type = 2)
kpAddBaseNumbers(kp, tick.dist = 100000, tick.len = 10, tick.col="red", cex=0.7, minor.tick.dist = 10000, minor.tick.len = 5, minor.tick.col = "gray")
AG006_coords_df <- read_tsv("AG038_AG002_concat.txt", col_names = T)
AG006_coords_df
Br62_coords_df <- read_tsv("AG002_concat_AG038.txt", col_names = T)
Br62_coords_df
AG006_coords_ranges <- toGRanges(data.frame(chr=c(AG006_coords_df$contig), start=c(AG006_coords_df$start), end=c(AG006_coords_df$end), strand=c(AG006_coords_df$strand)))
AG006_coords_ranges
Br62_coords_ranges <- toGRanges(data.frame(chr=c(Br62_coords_df$contig), start=c(Br62_coords_df$start), end=c(Br62_coords_df$end), strand=c(Br62_coords_df$strand)))
Br62_coords_ranges
karyoploteR::kpPlotLinks(kp, data = AG006_coords_ranges, data2 = Br62_coords_ranges, y= -0.1, col="dimgray", cex=0.2)

#AG002_AG006
custom.genome <- toGRanges(data.frame(chr=c("AG002_Contig06-7","AG006_Contig03"), start=c(1,1), end=c(1632000, 1697966)))
kp <- plotKaryotype(genome = custom.genome, plot.type = 2)
kpAddBaseNumbers(kp, tick.dist = 100000, tick.len = 10, tick.col="red", cex=0.7, minor.tick.dist = 10000, minor.tick.len = 5, minor.tick.col = "gray")
AG006_coords_df <- read_tsv("AG002_concat_AG006.txt", col_names = T)
AG006_coords_df
Br62_coords_df <- read_tsv("AG006_AG002_concat.txt", col_names = T)
Br62_coords_df
AG006_coords_ranges <- toGRanges(data.frame(chr=c(AG006_coords_df$contig), start=c(AG006_coords_df$start), end=c(AG006_coords_df$end), strand=c(AG006_coords_df$strand)))
AG006_coords_ranges
Br62_coords_ranges <- toGRanges(data.frame(chr=c(Br62_coords_df$contig), start=c(Br62_coords_df$start), end=c(Br62_coords_df$end), strand=c(Br62_coords_df$strand)))
Br62_coords_ranges
karyoploteR::kpPlotLinks(kp, data = AG006_coords_ranges, data2 = Br62_coords_ranges, y= -0.1, col="dimgray", cex=0.2)

#AG006_San_Andrea
custom.genome <- toGRanges(data.frame(chr=c("AG006_Contig03","San_Andrea_Contig07"), start=c(1,1), end=c(1697966, 1639350)))
kp <- plotKaryotype(genome = custom.genome, plot.type = 2)
kpAddBaseNumbers(kp, tick.dist = 100000, tick.len = 10, tick.col="red", cex=0.7, minor.tick.dist = 10000, minor.tick.len = 5, minor.tick.col = "gray")
AG006_coords_df <- read_tsv("AG006_to_SanAndrea_10k.bed", col_names = T)
AG006_coords_df
Br62_coords_df <- read_tsv("SanAndrea_to_AG006_10k.bed", col_names = T)
Br62_coords_df
AG006_coords_ranges <- toGRanges(data.frame(chr=c(AG006_coords_df$contig), start=c(AG006_coords_df$start), end=c(AG006_coords_df$end), strand=c(AG006_coords_df$strand)))
AG006_coords_ranges
Br62_coords_ranges <- toGRanges(data.frame(chr=c(Br62_coords_df$contig), start=c(Br62_coords_df$start), end=c(Br62_coords_df$end), strand=c(Br62_coords_df$strand)))
Br62_coords_ranges
karyoploteR::kpPlotLinks(kp, data = AG006_coords_ranges, data2 = Br62_coords_ranges, y= -0.1, col="dimgray", cex=0.2)

#AG059_SanAndrea
custom.genome <- toGRanges(data.frame(chr=c("San_Andrea_Contig07", "AG059_Contig05"), start=c(1,1), end=c(1639350, 1697966)))
kp <- plotKaryotype(genome = custom.genome, plot.type = 2)
kpAddBaseNumbers(kp, tick.dist = 100000, tick.len = 10, tick.col="red", cex=0.7, minor.tick.dist = 10000, minor.tick.len = 5, minor.tick.col = "gray")
AG006_coords_df <- read_tsv("AG059_Contig05_San_Andrea.txt", col_names = T)
Br62_coords_df <- read_tsv("San_Andrea_AG059_Contig05.txt", col_names = T)
AG006_coords_ranges <- toGRanges(data.frame(chr=c(AG006_coords_df$contig), start=c(AG006_coords_df$start), end=c(AG006_coords_df$end), strand=c(AG006_coords_df$strand)))
AG006_coords_ranges
Br62_coords_ranges <- toGRanges(data.frame(chr=c(Br62_coords_df$contig), start=c(Br62_coords_df$start), end=c(Br62_coords_df$end), strand=c(Br62_coords_df$strand)))
Br62_coords_ranges
karyoploteR::kpPlotLinks(kp, data = AG006_coords_ranges, data2 = Br62_coords_ranges, y= -0.1, col="dimgray", cex=0.2)
```
