## Supplementary material for "Multiple horizontal mini-chromosome transfers drive genome evolution of clonal blast fungus lineages": Barragan_2024_mChrA_SupplementaryData: FST_mChrA.R

```
############ written by Cristina Barragan Feb 2024
###########

getwd()

library(ggplot2)
library(dplyr)
library(tidyverse)

CR<-read.delim(file="pangenome.snps.filtered.bi.md09.Eleusine_Oryza.2.plink.5k.05k.windowed.weir.fst")

##now only values between 0 and 1
# Exclude negative WEIGHTED_FST values
CR <- CR[CR$WEIGHTED_FST >= 0, ]
CR
# Convert CHROM to factor and ensure its levels are in ascending order
CR$CHROM <- factor(CR$CHROM, levels = unique(sort(CR$CHROM)))
CR$CHROM
# Calculate the number of data points for each CHROM
counts <- as.data.frame(table(CR$CHROM))

# Boxplot with individual data points and count annotations
ggplot(CR, aes(x = as.factor(CHROM), y = WEIGHTED_FST)) +
  geom_boxplot(outlier.shape = NA) +  # Set outlier.shape to NA to hide individual outlier points, as we'll plot them with geom_jitter
  geom_jitter(width = 0.2, alpha = 0.2, size = 1, color = "gray40") +
  geom_text(data = counts, aes(x = Var1, y = max(CR$WEIGHTED_FST, na.rm = F), label = Freq), vjust = -0.5) +
  labs(title = "Boxplot of WEIGHTED_FST by CHROM with Data Points",
       x = "CHROM",
       y = "WEIGHTED_FST") +
  ylim(0, 1) +
  theme_minimal()
########################### only plot some Chromosomes
# Desired chromosomes to plot
desired_chroms <- c(1, 2, 6, 8, 9, 10, 12, 13)
# Filter the data
CR <- CR[CR$CHROM %in% desired_chroms, ]
CR
# Convert CHROM to factor and ensure its levels are in the specified order
CR$CHROM <- factor(CR$CHROM, levels = desired_chroms)
CR$CHROM 
# Calculate the number of data points for each CHROM
counts <- as.data.frame(table(CR$CHROM))
counts
# Boxplot with individual data points and count annotations
ggplot(CR, aes(x = CHROM, y = WEIGHTED_FST)) +
  geom_boxplot(outlier.shape = NA) +
  geom_jitter(width = 0.2, alpha = 0.2, size = 1, color = "gray40") +
  geom_text(data = counts, aes(x = Var1, y = max(CR$WEIGHTED_FST, na.rm = TRUE), label = Freq), vjust = -0.5) +
  labs(title = "Boxplot of WEIGHTED_FST by CHROM with Data Points",
       x = "CHROM",
       y = "WEIGHTED_FST") +
  ylim(0, 1) +
  theme_minimal()
```
