## Supplementary material for "Multiple horizontal mini-chromosome transfers drive genome evolution of clonal blast fungus lineages": Barragan_2024_mChrA_SupplementaryData: PCA_mChrA.R

```
########################################################################
########written by Cristina Barragan Feb 2024 ########################

# Load the eigenvalues and eigenvectors

eigenvals <- read.table("126indv_mChrA_samples.new.mChrA.snps.filtered.maxmiss10.plink.eigenval", header = FALSE)[,1]
eigenvectors <- read.table("126indv_mChrA_samples.new.mChrA.snps.filtered.maxmiss10.plink.edited2.eigenvec", header = T)

eigenvals <- read.table("126indv_mChrA_samples.new.coreChr.snps.filtered.maxmiss10.eigenval", header = FALSE)[,1]
eigenvectors <- read.table("126indv_mChrA_samples.new.coreChr.snps.filtered.maxmiss10.edited2.eigenvec", header = T)

eigenvals
eigenvectors

# Load the ggplot2 library
library(ggplot2)

# Create a PCA plot

############plot with desirec colors

ggplot(eigenvectors, aes(x = PC1, y = PC2, label = IID, color = TOY)) +
  geom_text(hjust = 1.2, vjust = -0.1, size = 4) +  # Adjust position and size of text
  scale_color_manual(values = c("Triticum" = "#1D76B8",
                                "Eleusine" = "#F27423",
                                "Brachiaria1" = "#989898",
                                "Oryza" = "#056A38",
                                "Lolium" = "#9E2064",
                                "Eragrostis"="#CD7F29")) + # Map TOY to colors
  labs(x = "PCA1: 42.30%", y = "PCA2:21.13%", title = "126 indv coreChr") +
  theme_minimal()


# Assuming 'eigenvals' is a vector of eigenvalues

# Calculate the total sum of eigenvalues
total_sum <- sum(eigenvals)

# Calculate the percentage of variation explained by each component
percentage_explained <- (eigenvals / total_sum) * 100

# Print the results
for (i in seq_along(percentage_explained)) {
  cat(paste("PC", i, ": ", percentage_explained[i], "%\n"))
}
```
