## Supplementary material for "Multiple horizontal mini-chromosome transfers drive genome evolution of clonal blast fungus lineages": Barragan_2024_mChrA_SupplementaryData: circlize_AG006_Br62_B71_LpKY97.R

```
## Initial script outline by Thorsten Langner and Angus Malmgren and further developed by Cristina Barragan Feb 2024
#######################################################################################################################
library(tidyr)
library(circlize)
library(dplyr)
library(tidyverse)

############ start
######AG006 and Br62
chr <- c("AG006_Contig01", "AG006_Contig02", "AG006_Contig03", "AG006_Contig04", "AG006_Contig05", "AG006_Contig06", "AG006_Contig07", "AG006_Contig08", "AG006_Contig09", "AG006_Contig10","AG006_Contig11", "AG006_Contig12", "AG006_Contig13", "AG006_Contig14", "AG006_Contig15", "AG006_Contig16", "AG006_Contig17", "AG006_Contig18","AG006_Contig19", "AG006_Contig20", "AG006_Contig21", "AG006_Contig22", "AG006_Contig23", "AG006_Contig24","Br62_Contig01", "Br62_Contig02", "Br62_Contig03","Br62_Contig04", "Br62_Contig06","Br62_Contig07", "Br62_Contig08", "Br62_Contig09","Br62_Contig10","Br62_Contig11")
start <- c(1,1,1,1,1,1,1,1,1,1,1,1,1,1,1,1,1,1,1,1,1,1,1,1,1,1,1,1,1,1,1,1,1,1)
end <- c(11008917,8645043,1697966,796569,60573,6618557,51574,5731635,4595668,1201610,170974,1464303,4077659,55256,72685,77902,104405,133460,96136,73650,73030,52001,70829,75409,5686109,5098061,7337754,8318372,6027403,1185594,3851847,6285418,69893,84649)

#### AG006 for B71
chr <- c("AG006_Contig01", "AG006_Contig02", "AG006_Contig03", "AG006_Contig04", "AG006_Contig05", "AG006_Contig06", "AG006_Contig07", "AG006_Contig08", "AG006_Contig09", "AG006_Contig10","AG006_Contig11", 
         "AG006_Contig12", "AG006_Contig13", "AG006_Contig14", "AG006_Contig15", "AG006_Contig16", "AG006_Contig17", "AG006_Contig18","AG006_Contig19", "AG006_Contig20", "AG006_Contig21", "AG006_Contig22", "
         AG006_Contig23", "AG006_Contig24","CP060337.1chromosome_mini", "CP060330.1chromosome_1", "CP060331.1chromosome_2","CP060332.1chromosome_3", "CP060333.1chromosome_4","CP060334.1chromosome_5", "CP060335.1chromosome_6", 
         "CP060336.1chromosome_7","CP060338.1mitochondrion,_complete_genome")
start <- c(1,1,1,1,1,1,1,1,1,1,1,1,1,1,1,1,1,1,1,1,1,1,1,1,1,1,1,1,1,1,1,1,1)
end <- c(11008917,8645043,1697966,796569,60573,6618557,51574,5731635,4595668,1201610,170974,1464303,4077659,55256,72685,77902,104405,133460,96136,73650,73030,52001,70829,75409,1903245,6668242,7915472,8237918,5413369,4460276,6133529,4079358,34996)

####AG006 and LpKY97
chr <- c("AG006_Contig01", "AG006_Contig02", "AG006_Contig03", "AG006_Contig04", "AG006_Contig05", "AG006_Contig06", "AG006_Contig07", "AG006_Contig08", "AG006_Contig09", "AG006_Contig10","AG006_Contig11", 
         "AG006_Contig12", "AG006_Contig13", "AG006_Contig14", "AG006_Contig15", "AG006_Contig16", "AG006_Contig17", "AG006_Contig18","AG006_Contig19", "AG006_Contig20", "AG006_Contig21", "AG006_Contig22", "
         AG006_Contig23", "AG006_Contig24","CP050920.1", "CP050927.1", "CP050921.1","CP050928.1", "CP050922.1","CP050923.1", "CP050924.1", 
         "CP050925.1","CP050926.1")
start <- c(1,1,1,1,1,1,1,1,1,1,1,1,1,1,1,1,1,1,1,1,1,1,1,1,1,1,1,1,1,1,1,1,1)
end <- c(11008917,8645043,1697966,796569,60573,6618557,51574,5731635,4595668,1201610,170974,1464303,4077659,55256,72685,77902,104405,133460,96136,73650,73030,52001,70829,75409,6359358,2996683,7985541,913334,7463082,5442467,4430380,6155580,3866546)

## B71 and LpKY97
chr <- c("CP050920.1", "CP050927.1", "CP050921.1","CP050928.1", "CP050922.1","CP050923.1", "CP050924.1", 
         "CP050925.1","CP050926.1", "CP060337.1chromosome_mini", "CP060330.1chromosome_1", "CP060331.1chromosome_2","CP060332.1chromosome_3", "CP060333.1chromosome_4","CP060334.1chromosome_5", "CP060335.1chromosome_6", 
         "CP060336.1chromosome_7","CP060338.1mitochondrion,_complete_genome")
start <- c(1,1,1,1,1,1,1,1,1,1,1,1,1,1,1,1,1,1)
end <- c(6359358,2996683,7985541,913334,7463082,5442467,4430380,6155580,3866546,1903245,6668242,7915472,8237918,5413369,4460276,6133529,4079358,34996)


###########
df <- data.frame(chr, start, end)
df
dim(df)
circos.initializeWithIdeogram(df)

#adding rainbow color to chromosomes
circos.track(ylim = c(0, 1), bg.col = c(rainbow(50, alpha = 1)), bg.border = NA, track.height = 0.2)
AG006_coords_df <- read_tsv("AG006_Br62_algn_coords_long.bed", col_names = T)
AG006_coords_df <- read_tsv("AG006_B71_algn_coords_long.bed", col_names = T)
AG006_coords_df <- read_tsv("AG006_LpKY97_algn_coords_long.bed", col_names = T)
AG006_coords_df <- read_tsv("B71_LpKY97_algn_coords_long.bed", col_names = T)


AG006_coords_df
dim(AG006_coords_df)

Br62_coords_df <- read_tsv("Br62_AG006_algn_coords_long.bed", col_names = T)
Br62_coords_df <- read_tsv("Br62_AG006_algn_coords_long_test.bed", col_names = T)
Br62_coords_df <- read_tsv("B71_AG006_algn_coords_long.bed", col_names = T)
Br62_coords_df <- read_tsv("LpKY97_AG006_algn_coords_long.bed", col_names = T)
Br62_coords_df <- read_tsv("LpKY97_B71_algn_coords_long.bed", col_names = T)

Br62_coords_df
dim(Br62_coords_df)

#adding genomic links between two sets of bed files
#circos.genomicLink(AG006_coords_df, Br62_coords_df, col = rand_color(nrow(AG006_coords_df), transparency = 0.5), border = NA)

circos.genomicLink(AG006_coords_df, Br62_coords_df, col = "dimgrey", border = NA)
```
