## Supplementary figures and images for "Multiple horizontal mini-chromosome transfers drive genome evolution of clonal blast fungus lineages"

### AG002_gt2Mb_mini007_008_repeatWind100k.pdf

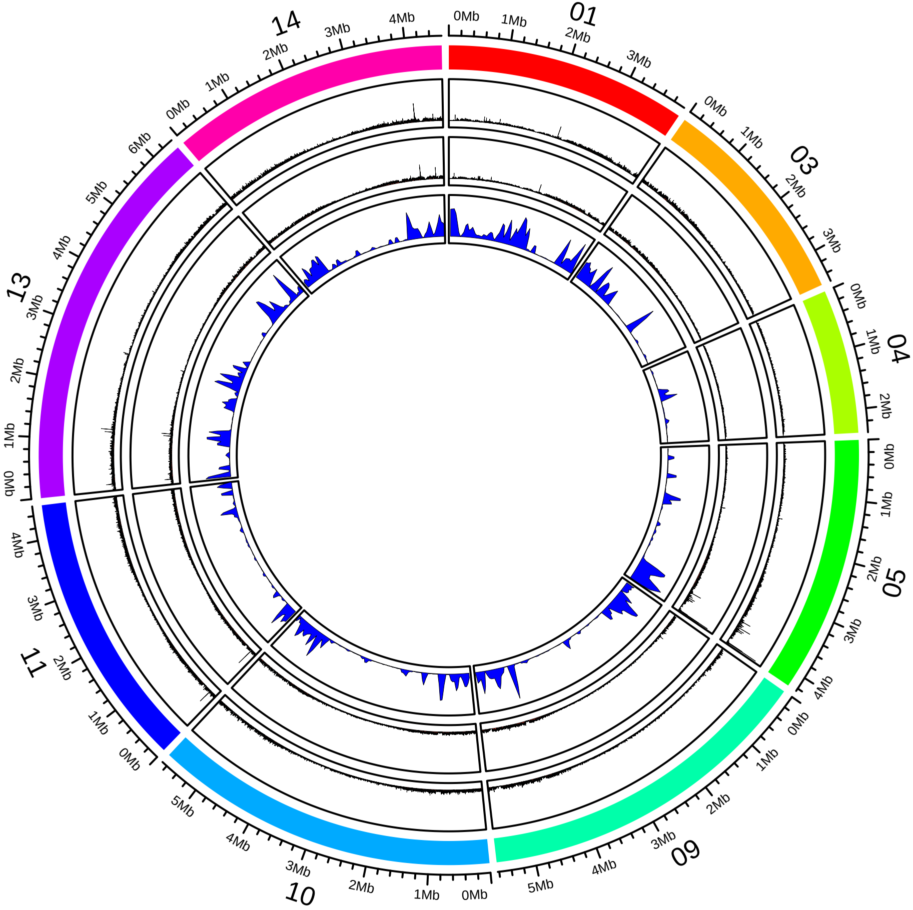

### AG002_lt2Mb_mini007_008_repeatWind10k.pdf

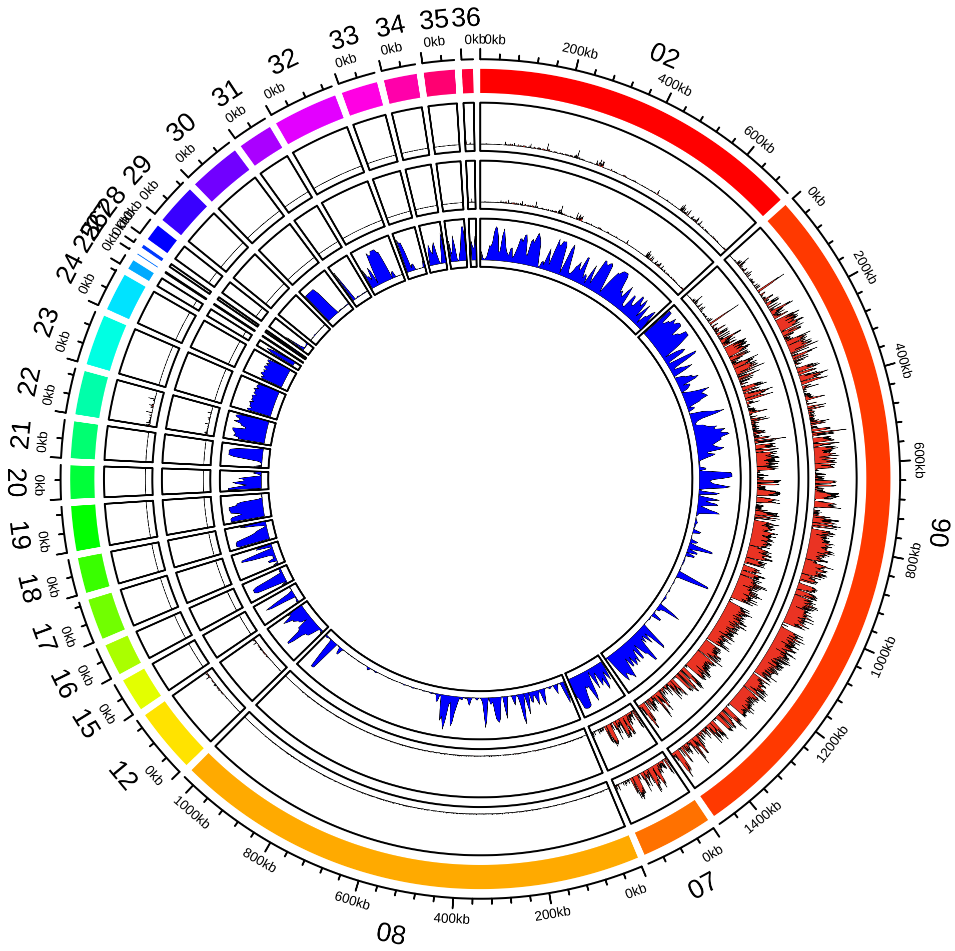

### AG006_gt2Mb_mini009_010_011_012_repeatWind100k.pdf

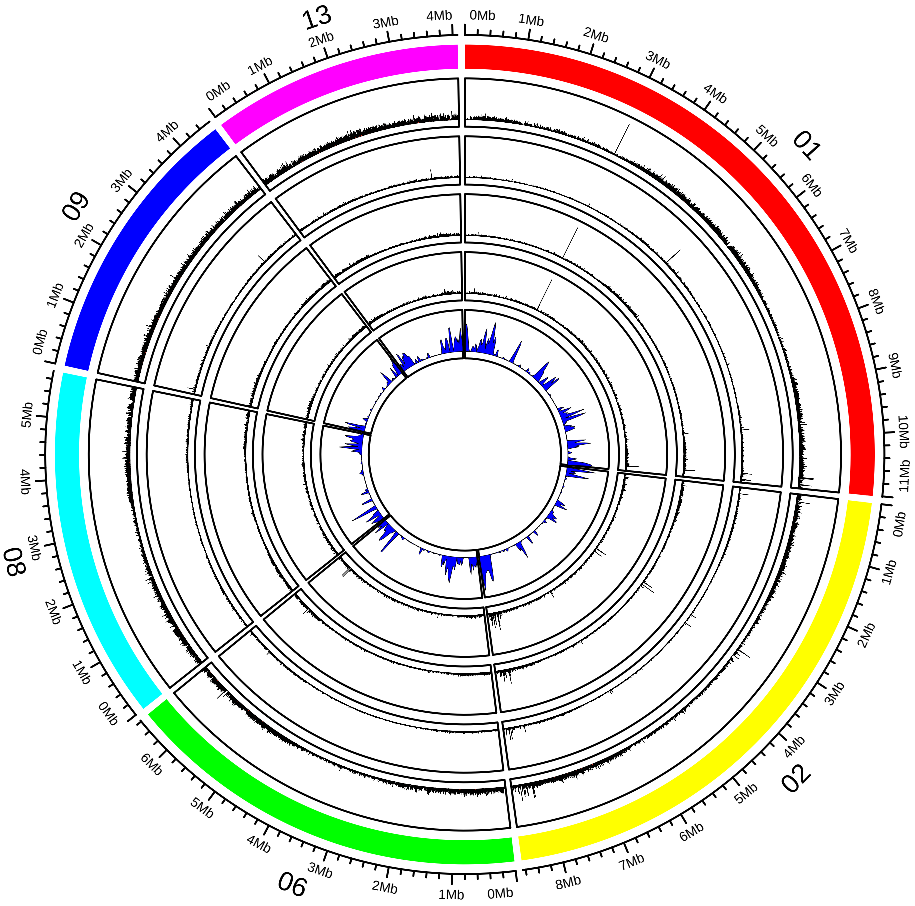

### AG006_lt2Mb_mini009_010_011_012_repeatWind10k.pdf

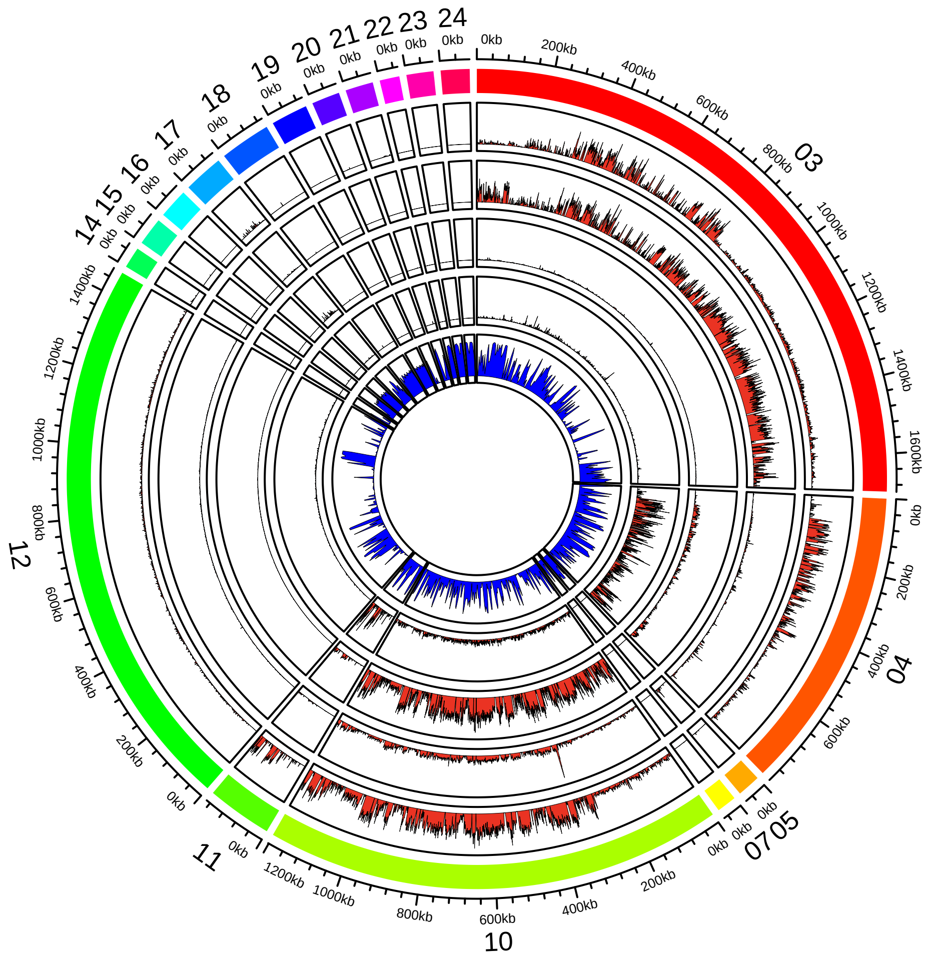

### AG032_gt2Mb_mini013_repeatWind100k.pdf

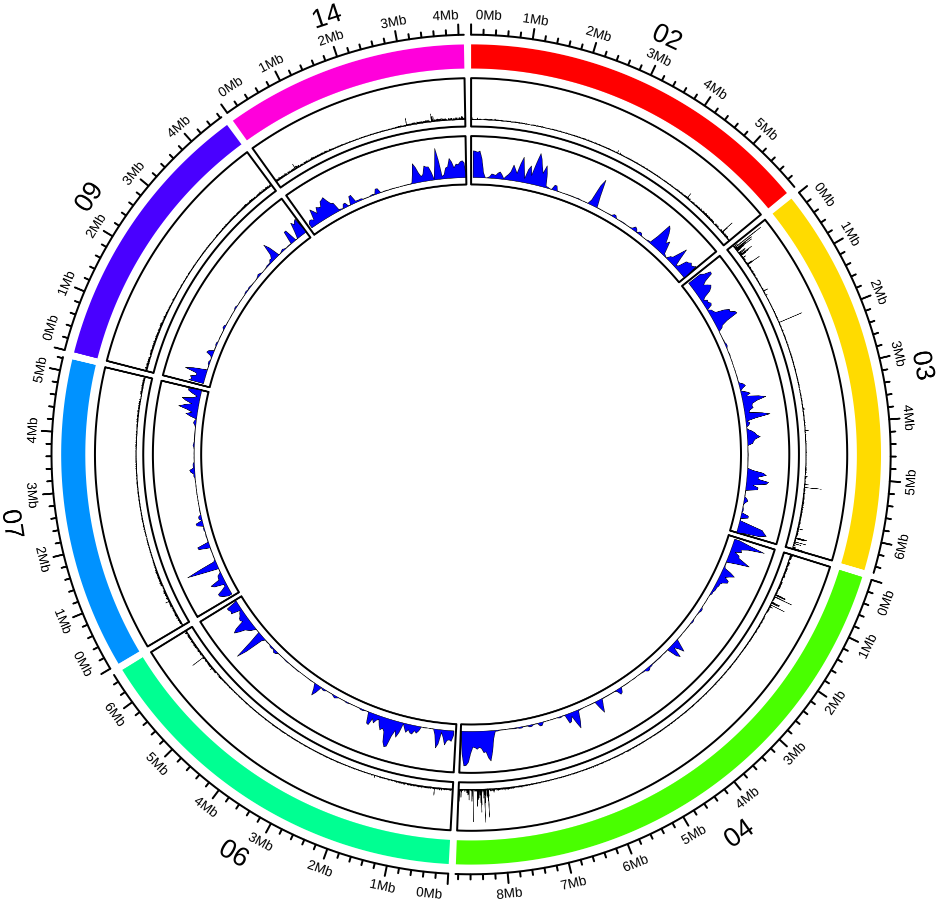

### AG032_lt2Mb_mini013_repeatWind10k.pdf

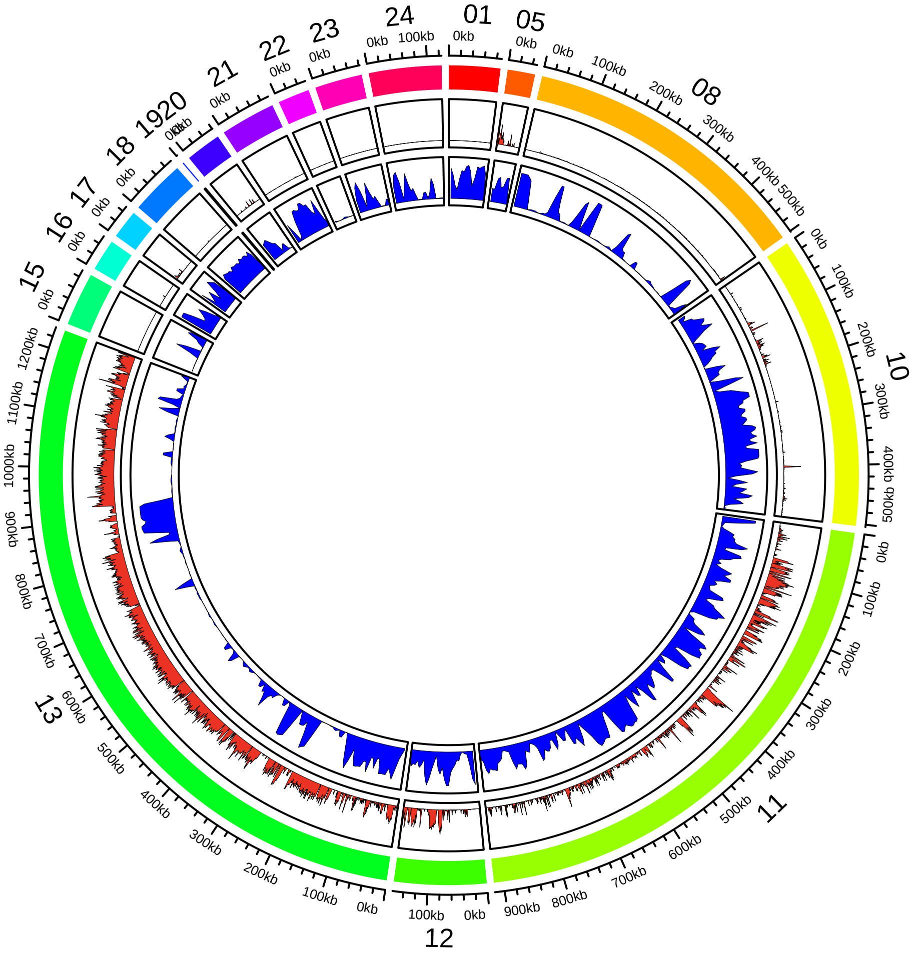

### AG038_gt2Mb_mini015_repeatWind100k.pdf

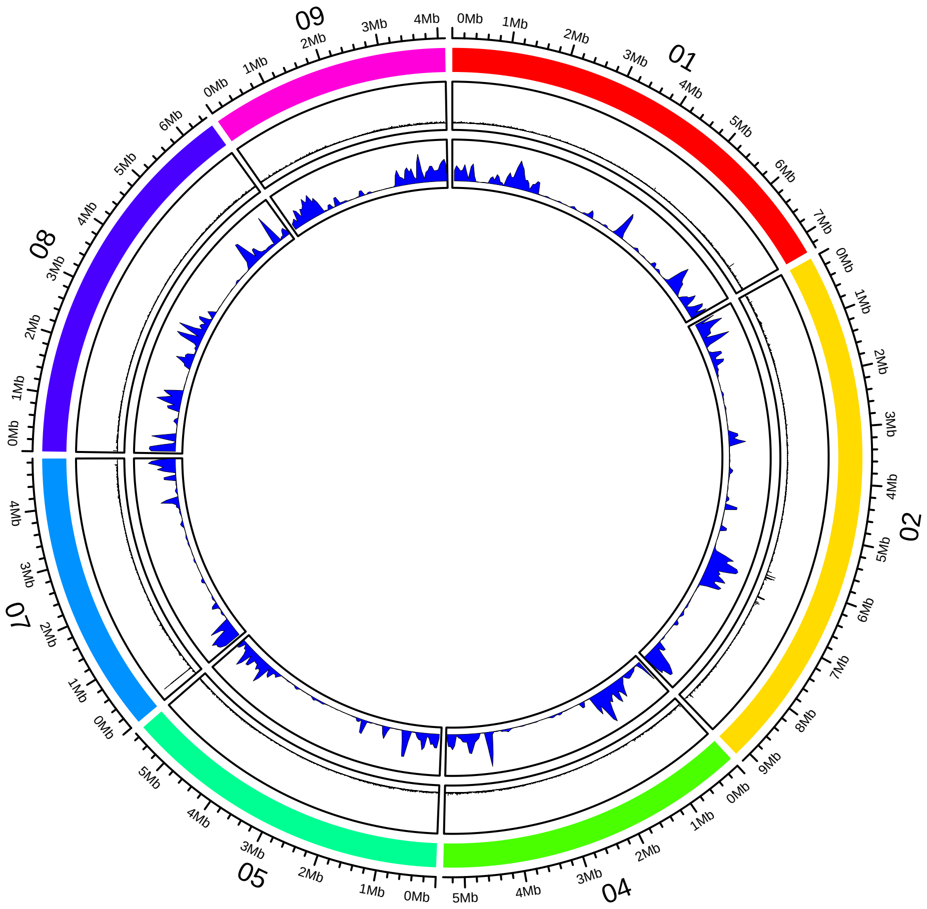

### AG038_lt2Mb_mini015_repeatWind10k.pdf

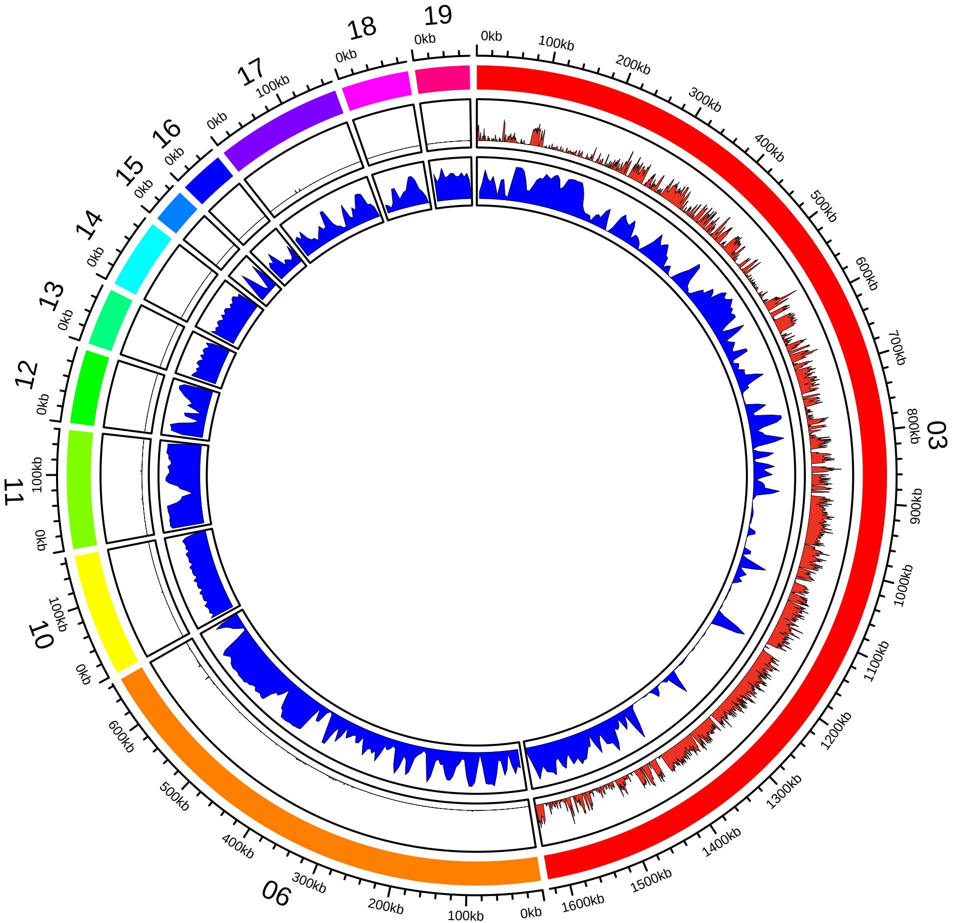

### AG039_gt2Mb_mini019_020_repeatWind100k.pdf

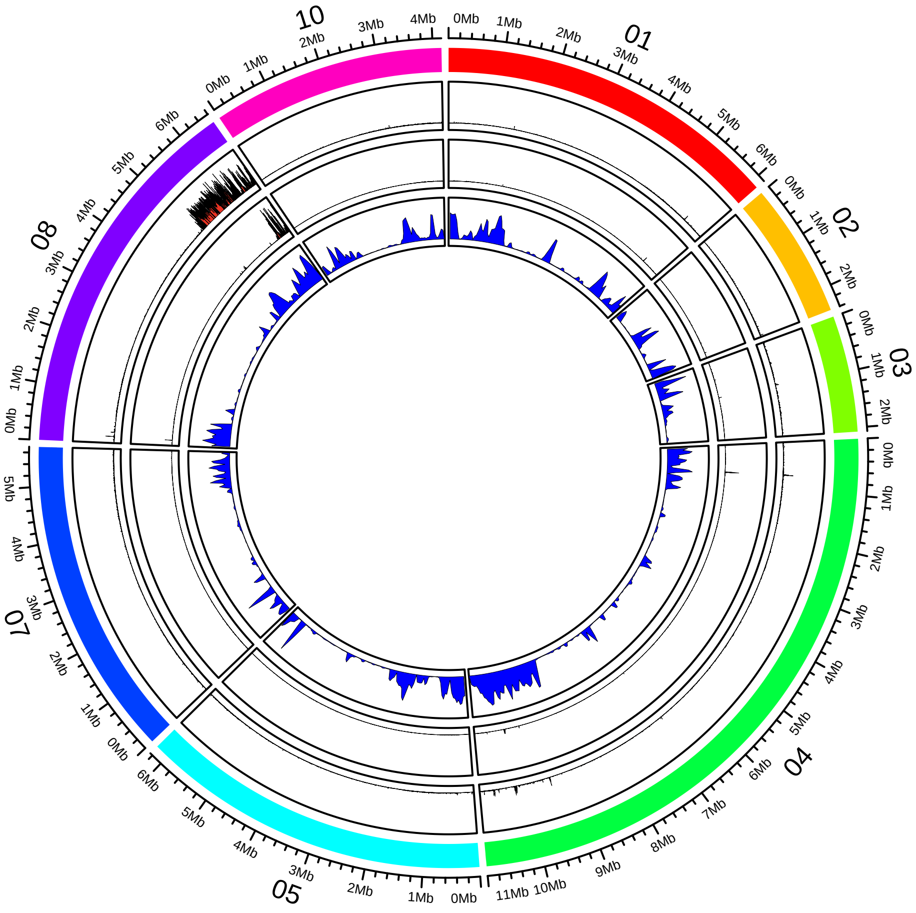

### AG039_lt2Mb_mini019_020_repeatWind10k.pdf

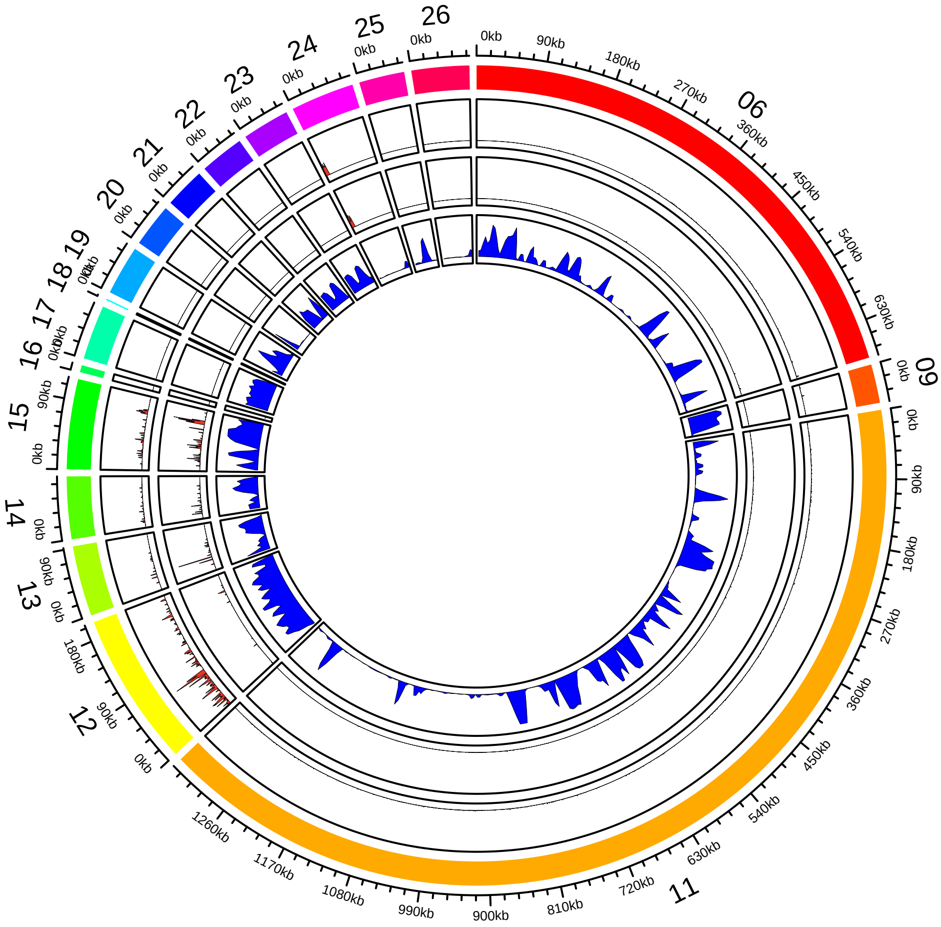

### AG059_gt2Mb_mini016_017_repeatWind100k.pdf

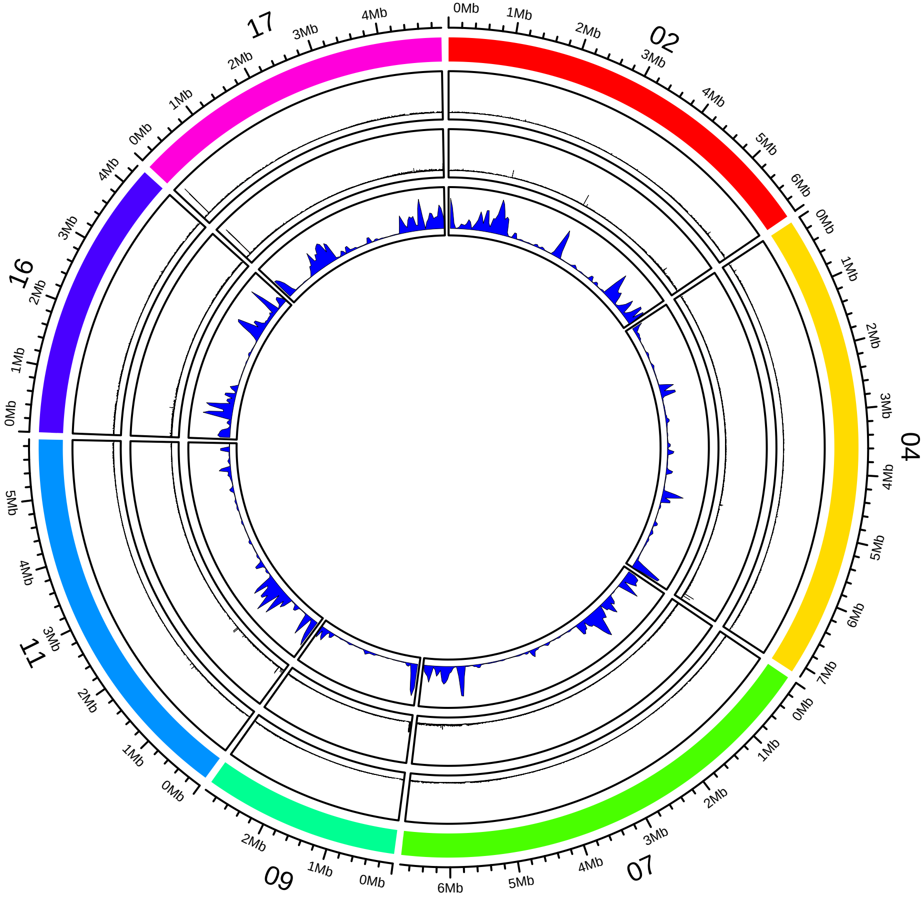

### AG059_lt2Mb_mini016_017_repeatWind10k.pdf

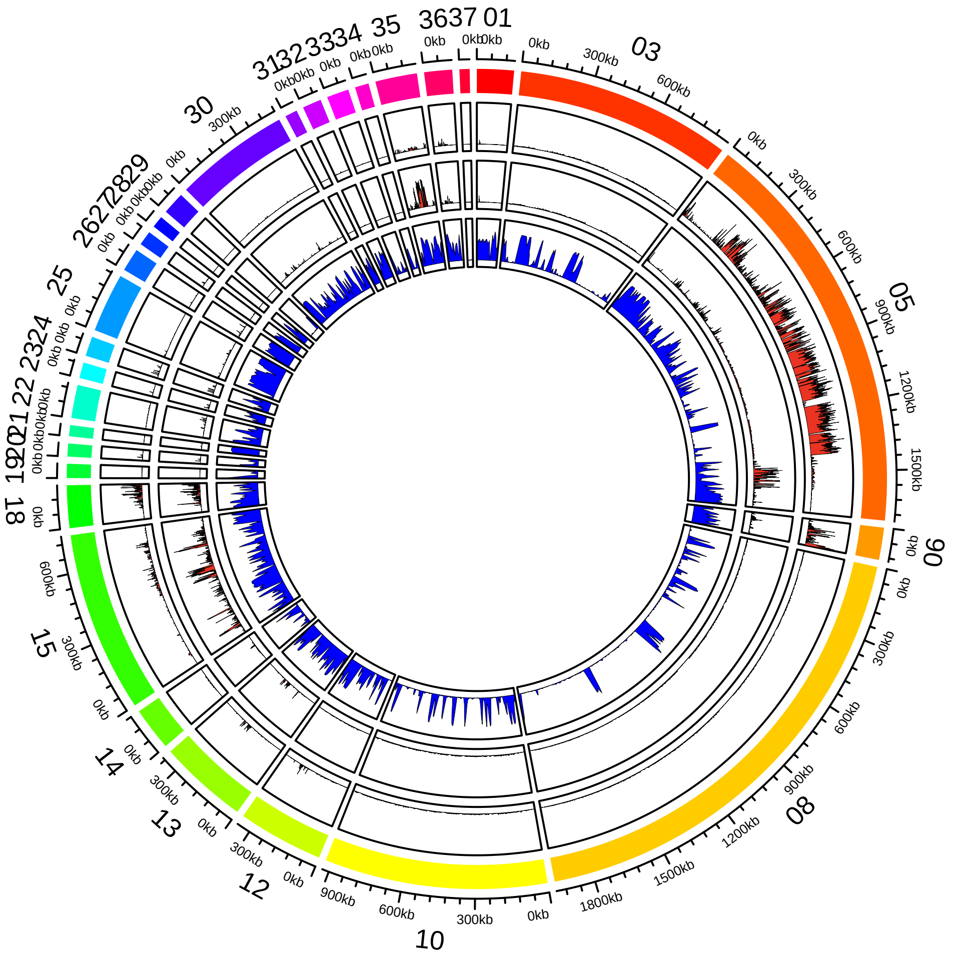

### AG098_gt2Mb_mini018_repeatWind100k.pdf

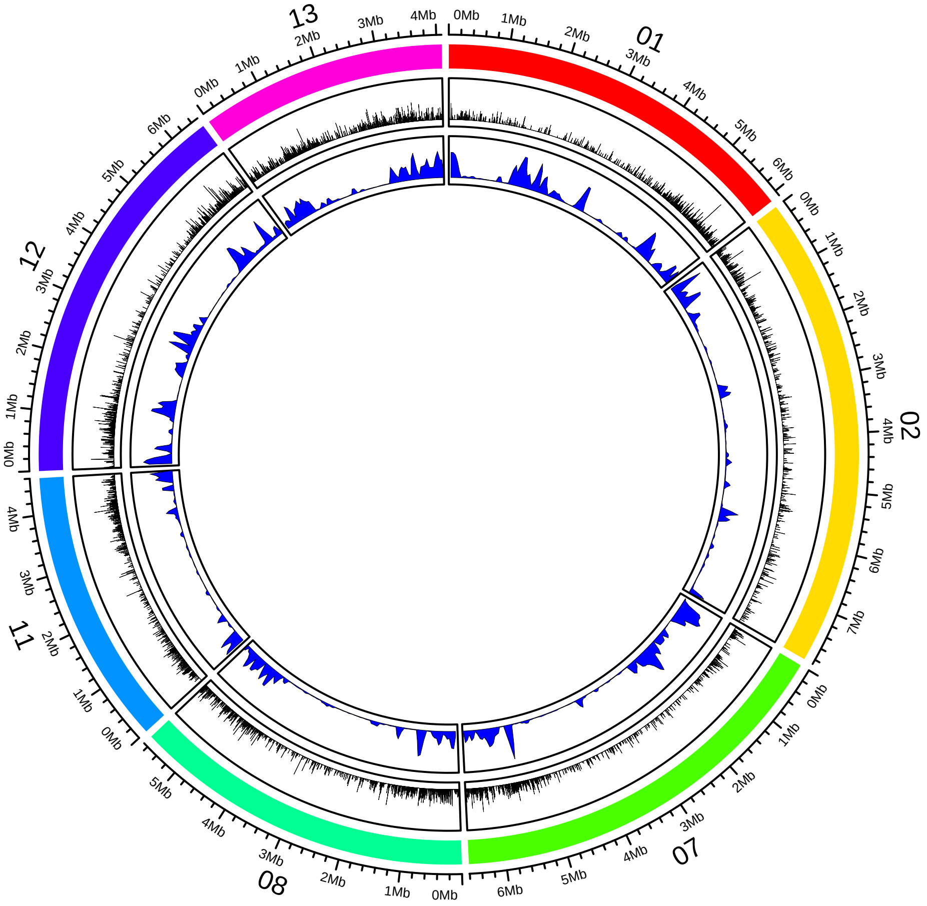

### AG098_lt2Mb_mini018_repeatWind10k.pdf

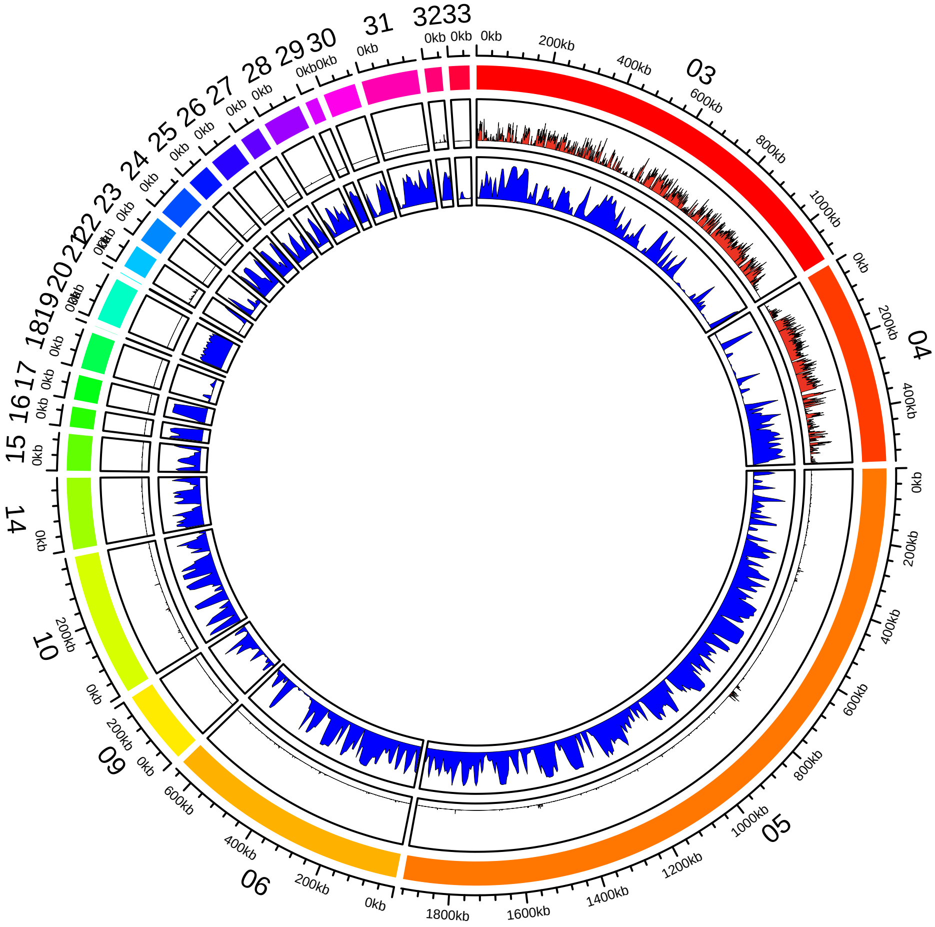

### Horizontal_mini_coverage_AG006_new.pdf

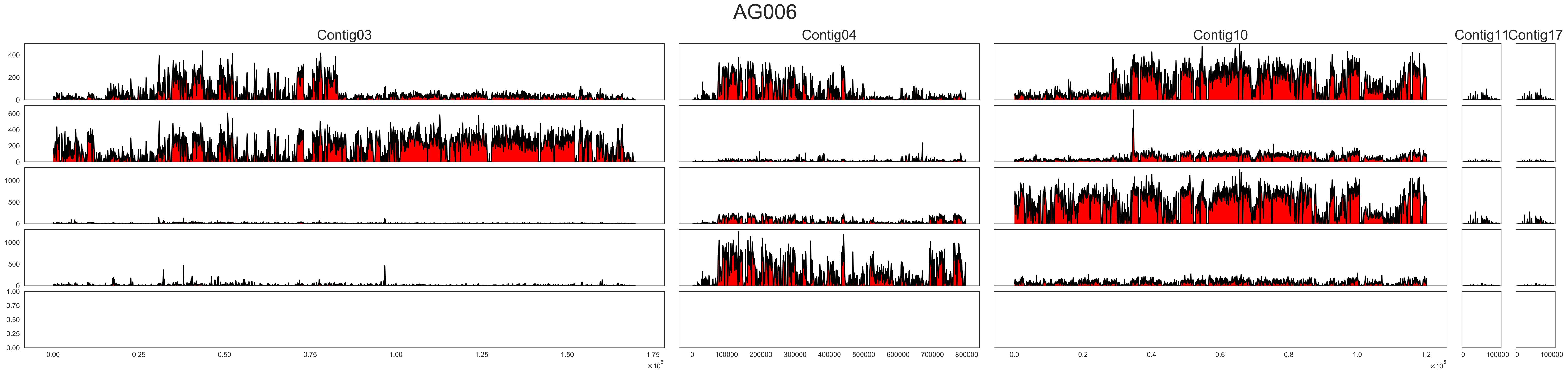

### PR003_gt2Mb_mini014_repeatWind100k.pdf

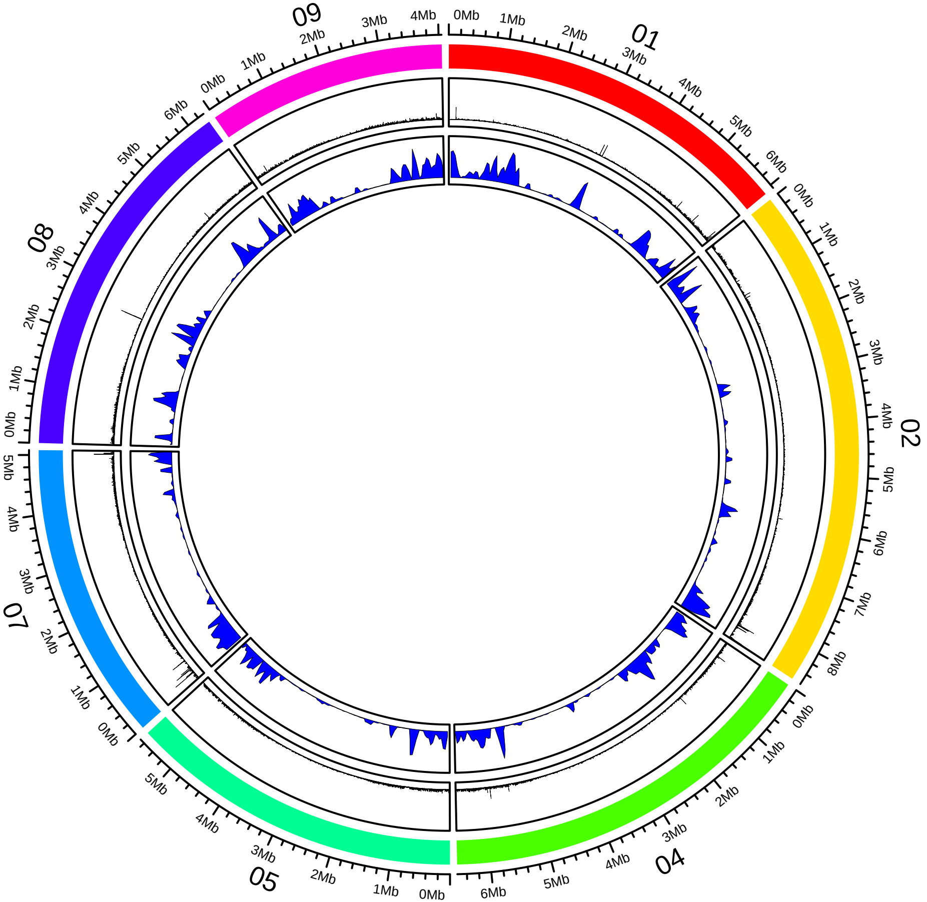

### PR003_lt2Mb_mini014_repeatWind10k.pdf

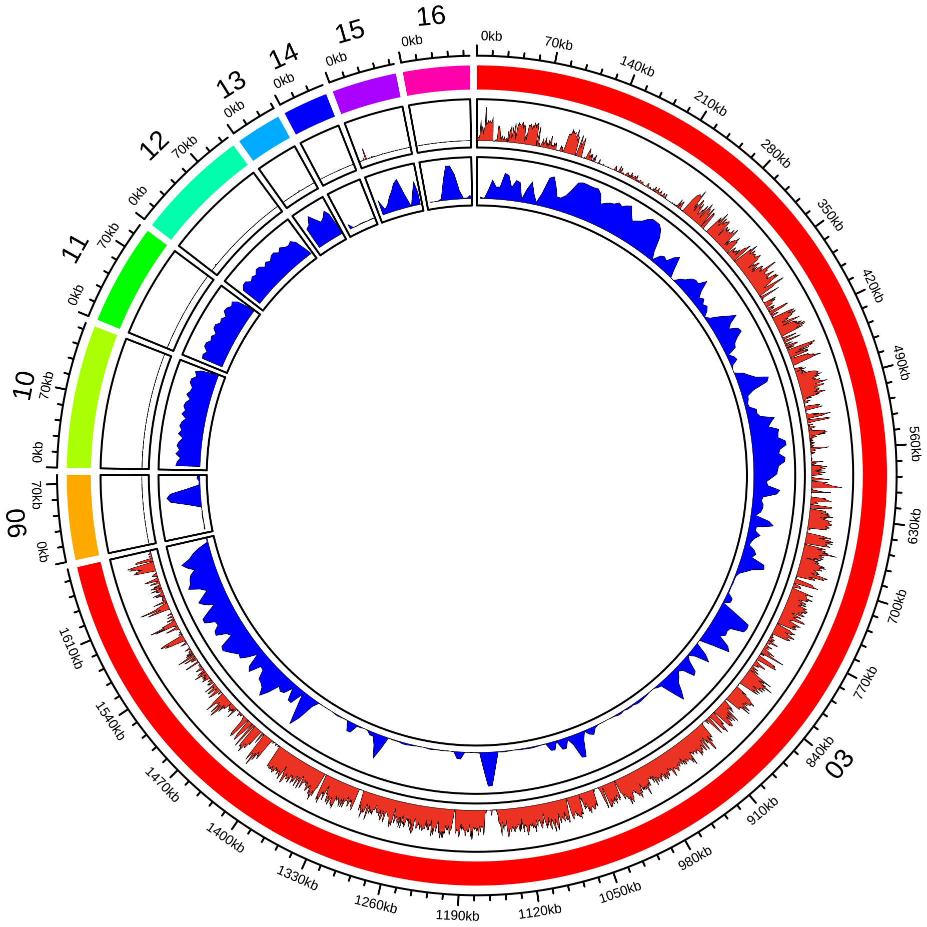

### San_Andrea_gt2Mb_mini021_022_023_repeatWind100k.pdf

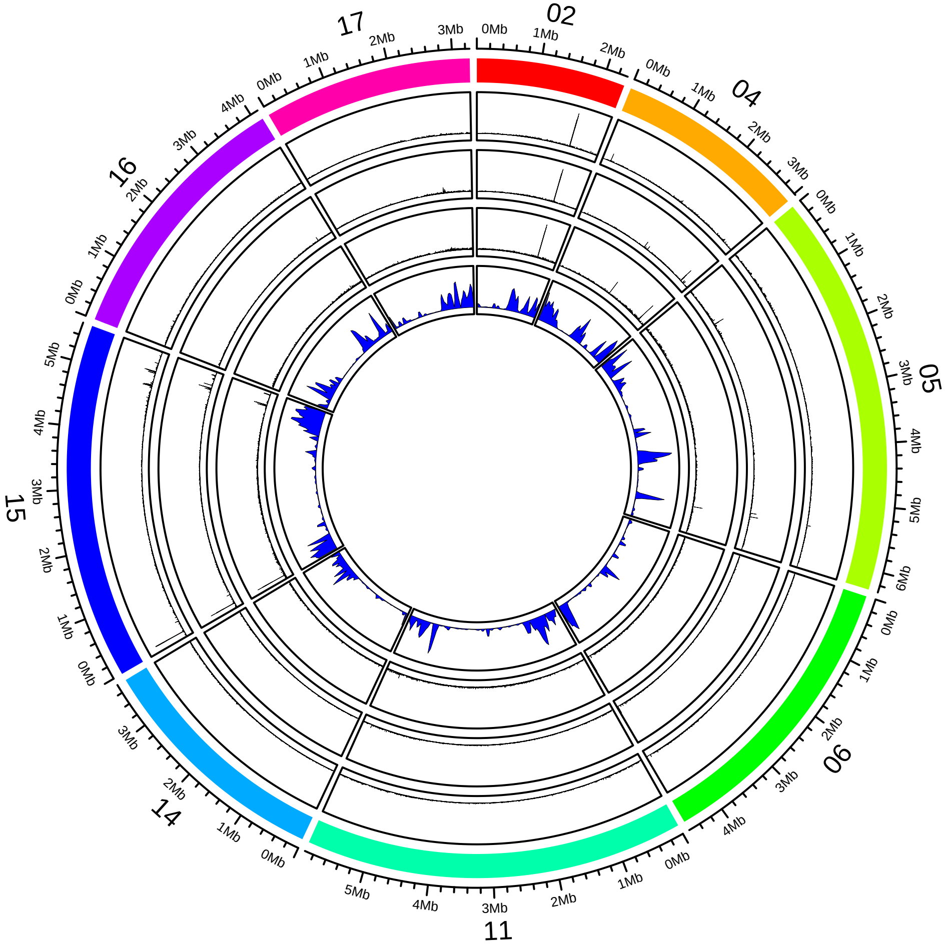

### San_Andrea_lt2Mb_mini021_022_023_repeatWind10k.pdf

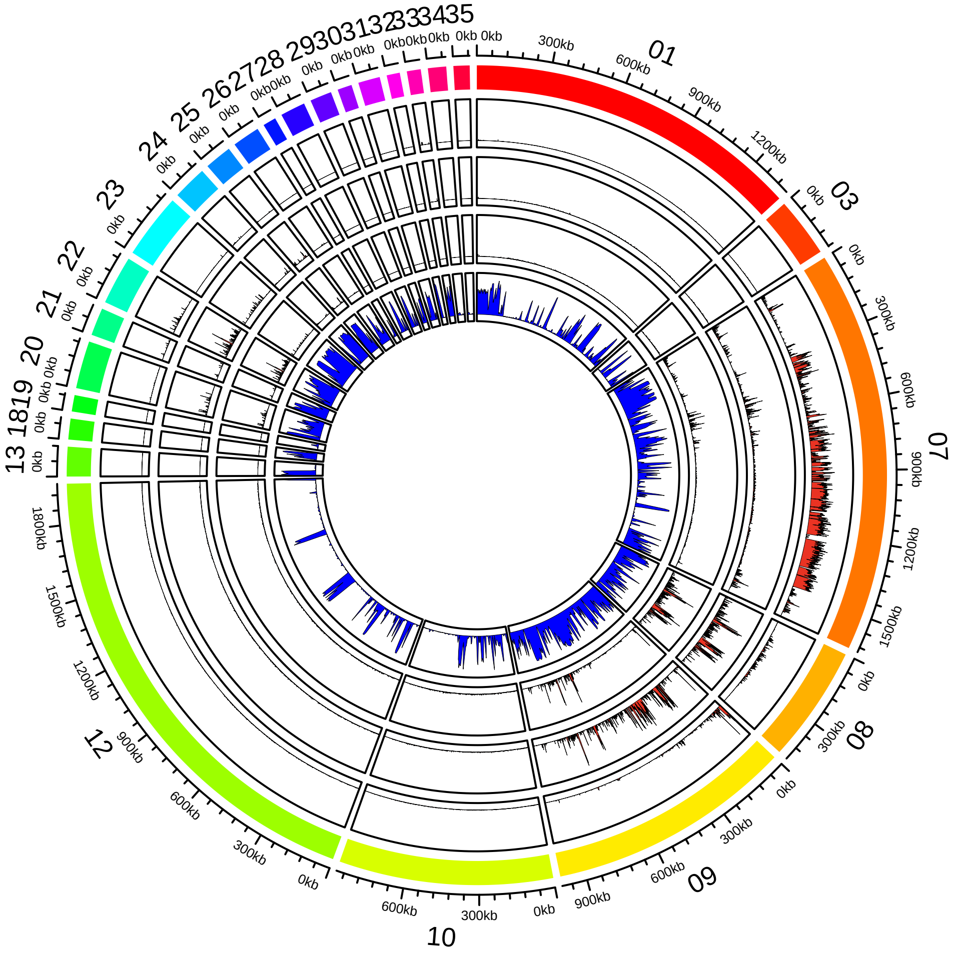
